# Dbf4-dependent kinase promotes meiotic DNA end resection through cyclin-dependent kinase 12 and DNA-2 in *Caenorhabditis elegans*

**DOI:** 10.64898/2026.06.17.732919

**Authors:** Anne Barbara Köhle, Angela Graf, Laura Kvedarauskaite, Lea Höglinger, Fränze Mueller, Felix Pinter, Alexander Schleiffer, Maria Rosaria Dello Stritto, Karl Mechtler, Markus Hartl, Joao Matos, Verena Jantsch, Antoine Baudrimont

## Abstract

Meiotic crossovers depend on the accurate processing of programmed DNA double-strand breaks by homologous recombination, which involves generating by resection the single-stranded DNA ends required for the subsequent strand invasion step. How regulatory kinases control DNA end resection in animal meiosis remains poorly understood. Here, we identify the *Caenorhabditis elegans* Dbf4-dependent kinase (DDK) complex, composed of CDC–7 and DBF–4, and identify CDK–12 as a candidate substrate. DDK promotes efficient meiotic DNA end resection, while crossover formation remains largely preserved in its absence. Loss of DDK function leads to SPO-11-dependent chromosome bridges at diakinesis that arise from recombination intermediates formed early in recombination. In addition to chromosome bridges, we observed elevated heterologous recombination that occurs independently of the canonical or alternative end-joining pathways. Interestingly, CDC–7 and DBF–4 localize dynamically during prophase I and load onto DNA in a hierarchical manner that depends on the BRC–1–BRD-1 complex. Epistasis analysis placed CDC–7 and CDK–12 within the DNA–2-dependent branch of long-range DNA resection. Together, our findings reveal an unidentified DDK/CDK–12/DNA–2 signaling axis that promotes efficient meiotic DNA resection to avoid the generation of aberrant recombination outcomes.

## Introduction

The two meiotic cell divisions produce haploid genomes in the gametes. For accurate segregation of parental homologs at meiosis I, a physical connection must be established between them to properly align chromosomes on the metaphase plate (GERTON AND HAWLEY 2005). During the prolonged prophase of meiosis I, a series of events leads to the formation of this connection via crossovers. Programmed meiotic DNA double-strand breaks (DSBs) are repaired by homologous recombination using one of the sister chromatids of the parental homolog as the repair template (HUNTER 2015). At the Spo11-mediated DSB, resection generates 3□ overhanging ends that form joint molecules via inter-homolog invasion, DNA synthesis, and capture of the second end of the DSB. These repair intermediates can be processed into crossovers via resolvases or into non-crossovers via nucleases along with the Bloom–topoisomerase–RMI1/2 complex, which dismantles joint molecules by decatenation (HUNTER 2015; GARTNER AND ENGEBRECHT 2022). In parallel, alternative repair pathways, including non-homologous end joining and microhomology-mediated end joining, can process meiotic DSBs, in particular when homologous recombination is compromised; under wild-type conditions, these pathways are generally suppressed to preserve genome integrity (HUNTER 2015; CECCALDI *et al*. 2016).

A core step of homologous recombination is the generation of 3– single-stranded DNA (ssDNA) overhanging ends. These are initially coated by the ssDNA-binding protein RPA (**r**eplication **p**rotein **A**), which is subsequently replaced by the RAD51 and DMC1 recombinases to promote strand invasion (SUGIYAMA AND KOWALCZYKOWSKI 2002; ZICKLER AND KLECKNER 2015). In yeast, resection is viewed as a two-step process in which the MRX (Mre11–Rad50–Xrs2) complex, together with Sae2/Ctip, cleaves Spo11-bound DNA to mediate short-range resection (MIMITOU AND SYMINGTON 2008; CANNAVO AND CEJKA 2014). The Mre11 endonuclease cleaves the DNA and degrades ssDNA toward the DSB via 3– to 5– exonuclease activity. Consecutive long-range resection is generated by the 5– to 3– nuclease activity of Exo1 (for review, see (KIM *et al*. 2025a)).

In worms, a similar mechanism is likely envisioned (CHIN AND VILLENEUVE 2001; YIN AND SMOLIKOVE 2013; KOURY *et al*. 2018; BELAN *et al*. 2021); however, a prominent role for DNA–2 in long-range resection has been proposed (HICKS *et al*. 2022), similar to the requirement for Dna2 in yeast mitotic resection (CECCALDI AND CEJKA 2025). In mouse spermatocytes, MRN–CTIP initiates and extends resection; EXO1 also contributes (KIM *et al*. 2025b) and other nucleases might be involved.

The complex chromosome transactions that occur during the prolonged prophase I, including chromatin loop organization, homolog pairing, DSB induction and processing into crossovers, and kinetochore emergence, must be coordinated with cell cycle progression and chromosome morphogenesis. Regulatory events are often controlled by post-translational modifications (KAR AND HOCHWAGEN 2021). Reversible phosphomodification is widely used to modulate protein–protein interaction surfaces, thus governing meiotic progression and implementing feedback control. Widely conserved signaling kinases with roles during prophase are also involved in *Caenorhabditis elegans* meiosis, including ATM-1, ATR-1, CDK-1, CDK-2, CHK-2, MPK-1, and Polo kinases (PLKs) (ARUR *et al*. 2011; HARPER *et al*. 2011; LABELLA *et al*. 2011; ROSU *et al*. 2013; STAMPER *et al*. 2013; WOGLAR *et al*. 2013; KIM *et al*. 2015; MACHOVINA *et al*. 2016; NADARAJAN *et al*. 2016; YIN *et al*. 2016; NADARAJAN *et al*. 2017; PATTABIRAMAN *et al*. 2017; FERRANDIZ *et al*. 2018; ACHACHE *et al*. 2019; GARCIA-MUSE *et al*. 2019; BRANDT *et al*. 2020; CASTELLANO-POZO *et al*. 2020; GUO *et al*. 2022; HAVERSAT *et al*. 2022; LASCAREZ-LAGUNAS *et al*. 2022; YU *et al*. 2023; ZHANG *et al*. 2023).

In yeast, the cell cycle Dbf4-dependent Cdc7 kinase (DDK) is a versatile kinase consisting of a catalytic subunit, Cell division cycle 7-related Ser/Thr kinase (Cdc7), and a regulatory subunit, Dumbbell-forming factor 4 (Dbf4). Yeast DDK operates at numerous stages of meiosis, including replication, DSB induction, crossover recombination, mono-polar attachment of sister kinetochores during the first meiotic division, and cohesion cleavage (VALENTIN *et al*. 2006; MATOS *et al*. 2008; SASANUMA *et al*. 2008; WAN *et al*. 2008; BRUCK AND KAPLAN 2009; NATSUME *et al*. 2013; MURAKAMI AND KEENEY 2014; CHEN *et al*. 2015; ARGUNHAN *et al*. 2018; ZHENG *et al*. 2018; HE *et al*. 2020).

Identified DDK substrates include the Mcm2–7 helicase, which is phosphorylated to drive replication and assist cohesin loading (VALENTIN *et al*. 2006; BRUCK AND KAPLAN 2009; NATSUME *et al*. 2013; ZHENG *et al*. 2018). The DSB-promoting protein Mer2 is phosphorylated to recruit the DSB-inducing protein Spo11 (SASANUMA *et al*. 2008; WAN *et al*. 2008; MURAKAMI AND KEENEY 2014). Furthermore, DDK phosphorylation of Msh4 ensures crossover recombination, and phosphorylation of the Zip1 synaptonemal complex protein drives synapsis between homologous chromosomes (CHEN *et al*. 2015; HE *et al*. 2020). Together with the yeast Polo kinase Cdc5, DDK mediates synapsis disassembly after crossover formation (ARGUNHAN *et al*. 2018). DDK-dependent phosphorylation of the Monopolin Lrs4 subunit facilitates homologous bi-orientation during metaphase I (LO *et al*. 2008; MATOS *et al*. 2008), and phosphorylation of Rec8 by DDK triggers its cleavage by separase (KATIS *et al*. 2010). An essential role for DDK in meiosis has been well established in yeast. However, we still lack insight into whether DDK activity is also essential for successful meiosis in animals: so far, only an involvement in the efficient recruitment of the DSB machinery has been shown in mice (DERELI *et al*. 2024).

Here, we report on the identification of the Dbf4-dependent kinase complex (DBF–4 and CDC–7) in *C. elegans* and identify CDK–12 as a potential substrate during the meiotic cell cycle. Cytogenetic analyses revealed that loading of the DDK complex onto chromosomes during prophase I is hierarchical and depends on the BRC–1–BRD-1 complex and DDK activity. Functional characterization of DDK mutants uncovered the formation of thin DNA bridges between chromosomes at diakinesis. Our data suggest that these bridges result from inefficient resection, form between chromatids of homologous and non-homologous chromosomes, and depend on both MSH-6 (mismatch recognition) and non-homologous end joining. Crossover recombination is largely unaffected; nevertheless, we detected translocations in the progeny that had arisen independently of the canonical and alternative end-joining pathways. Genetic analyses further indicated that CDC–7 acts in conjunction with CDK–12 and DNA–2 to mediate efficient DNA end resection. Together, these findings establish a previously uncharacterized role for DDK and CDK–12 in meiotic resection and faithful recombination that prevents chromosome rearrangements and segregation errors.

## Results

### Identification of the *C. elegans* Dbf4-dependent kinase with its regulatory subunit and the potential target, CDK–12

To investigate whether DDK plays a role in the meiotic cell cycle, we employed functionally tagged versions of CDC–7 (*cdc*–*7(jf218[cdc*–*7::HA]* and *cdc*–*7(syb6737[cdc*–*7::GFP]*, see Table S1 for strain viability) to perform native pull-downs and proximity labeling assays in triplicate (Figure 1A). By determining the overlap between these two complementary biochemical assays (Tables S2 and S3), we identified the following proteins with over 2-fold enrichment with 95% confidence: CDC–7 (our bait), the uncharacterized open reading frame F55H2.7, and components of the BRC–1–BRD-1 complex (Figure 1A). Bioinformatic analysis revealed that F55H2.7 belongs to the DBF-4 protein family (Figure S1A, see Materials and Methods). We will refer to F55H2.7 as DBF–4 hereafter. Indeed, AlphaFold (ABRAMSON *et al*. 2024) plausibly predicted the tertiary structure of the *C. elegans* DDK complex bound to ATP (Figures 1B and S1B).

**Figure 1.**
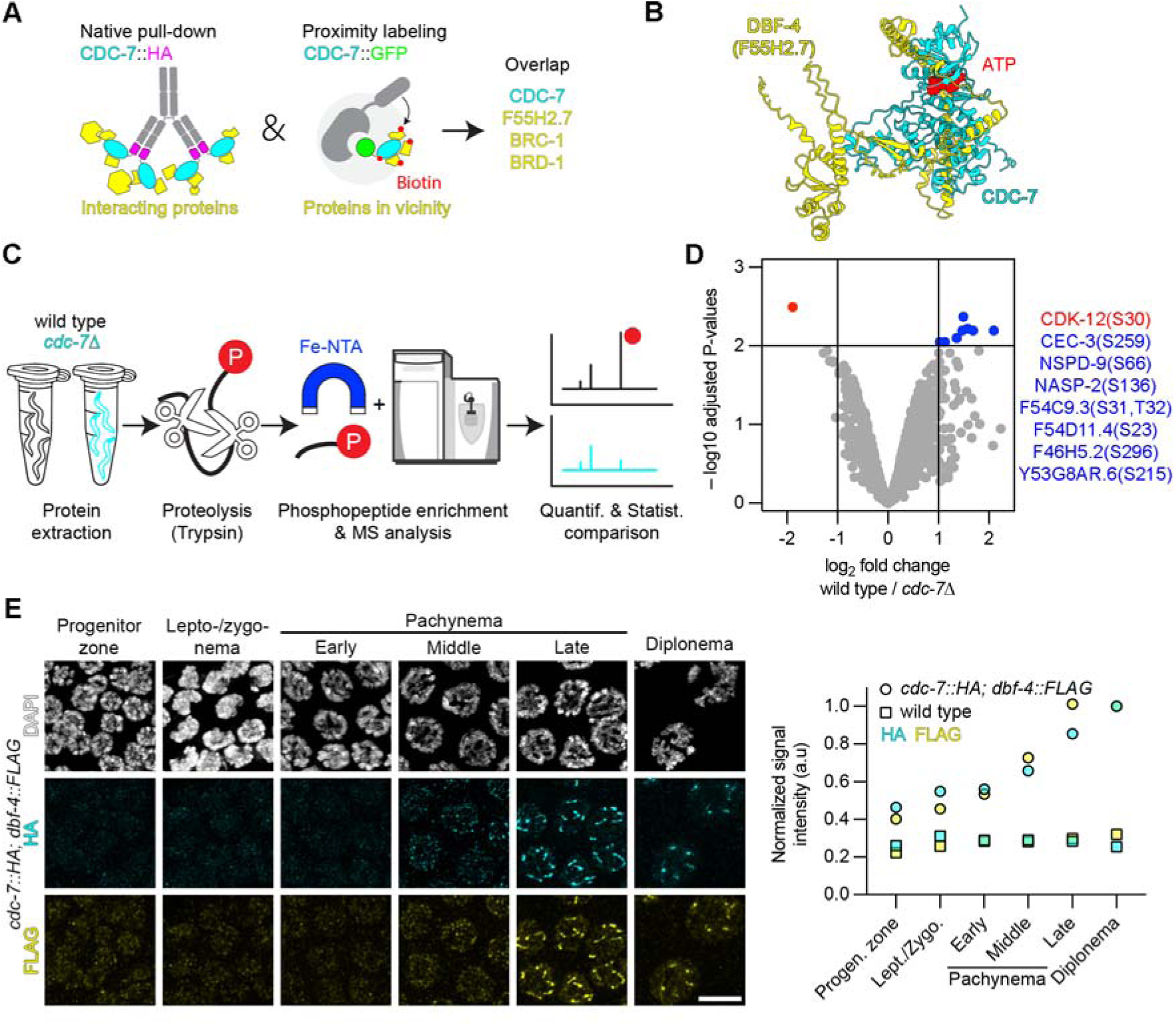
CDC–7-associated proximity labeling and phosphoproteomics reveal DDK complex members, candidate recruiters, and potential substrates during meiosis. **A.** Schematic diagram of the experimental strategy combining native pull-down of CDC–7::HA and proximity-dependent biotin labeling using CDC–7::GFP to identify interacting and proximal proteins. Overlapping candidates include F55H2.7, BRC–1, and BRD-1. **B.** AlphaFold prediction of the structure of the CDC–7–DBF–4 (F55H2.7) complex, highlighting the ATP-binding pocket. CDC–7 is shown in cyan, DBF–4 in yellow, and ATP in red (iPTM: 0.79). **C.** Workflow for quantitative phosphoproteomics. Protein extracts from wild-type and *cdc*–*7* mutant animals were subjected to tryptic digestion, Fe-NTA-based phosphopeptide enrichment, and mass spectrometry (MS), followed by quantitative and statistical analysis. **D.** Volcano plot of phosphopeptide abundance comparing the wild type with *cdc*–*7* mutants. Significantly enriched candidates are highlighted, including CDK–12(S30). **E.** Left, inset of representative nuclei in the zone of immunofluorescence of CDC–7::HA (cyan) and DBF–4::FLAG (yellow) localization across meiotic prophase, with DAPI counterstaining (white). Right, quantification of normalized signal intensity with arbitrary units (a.u.). Scale bar, 5 µm.

To identify targets of the Dbf4-dependent kinase, we set up a phosphomapping screen to compare phosphorylation levels after Fe-NTA column enrichment between the wild-type and *cdc*–*7(knu709)* deletion allele (CURREY AND LIACHKO 2019), hereafter referred to as *cdc*–*7*Δ (Figure 1C). We found a striking 3.7-fold reduction in a CDK–12 phosphopeptide at Ser30 (Figure 1D, red; Table S4). Comparative phosphoproteomics also identified phosphorylation sites that were upregulated over 2-fold upon DDK depletion (Figure 1D, blue).

To investigate the localization of both CDC–7 and DBF–4 in the *C. elegans* germline, we used genome-edited functional lines tagged with HA and FLAG epitopes (*cdc*–*7(jf218[cdc*–*7::HA]); dbf*–*4(jf247[dbf*–*4::FLAG])*; Table S1). Both CDC–7::HA and DBF–4::FLAG displayed nuclear enrichment, with numerous dim foci among a nuclear haze throughout the progenitor zone to early pachynema (Figure 1E). From mid-pachynema, the DDK complex formed dense aggregates culminating in stretches in late pachynema, which then retracted to form smaller stretches in diplonema. CDC–7 and DBF–4 not only displayed similar localization dynamics but also co-localized (Figure S1C). Furthermore, we found that CDC–7 colocalizes with the pro-crossover marker MSH-5 in late pachynema (KELLY *et al*. 2000; YOKOO *et al*. 2012) (Figure S1D).

In summary, we have identified known components of the *C. elegans* DDK complex and showed that they are expressed in the germline in distinct patterns. Furthermore, we identified a CDK–12 phosphopeptide that is significantly downregulated in *cdc*–*7*Δ and might, therefore, qualify as a potential substrate.

### *cdc*–*7* and *dbf*–*4* mutants display similar phenotypes, and their embryos are mostly viable

We next wanted to understand whether mutants in DDK complex genes would have similar phenotypes. Therefore, we compared embryonic lethality and the high incidence of male (Him) phenotype (a proxy for chromosome non-disjunction (HODGKIN *et al*. 1979)) across different alleles of DDK complex components (Figure 2A). The deletion allele *cdc*–*7*Δ and the *dbf*–*4* premature stop allele, *jf255* (hereafter, *dbf*–*4^STOP^*), which should not alter F55H2.8 transcript levels, both showed low embryonic lethality that, nevertheless, differed significantly from the wild type, as well as a Him phenotype (Figure 2A).

**Figure 2.**
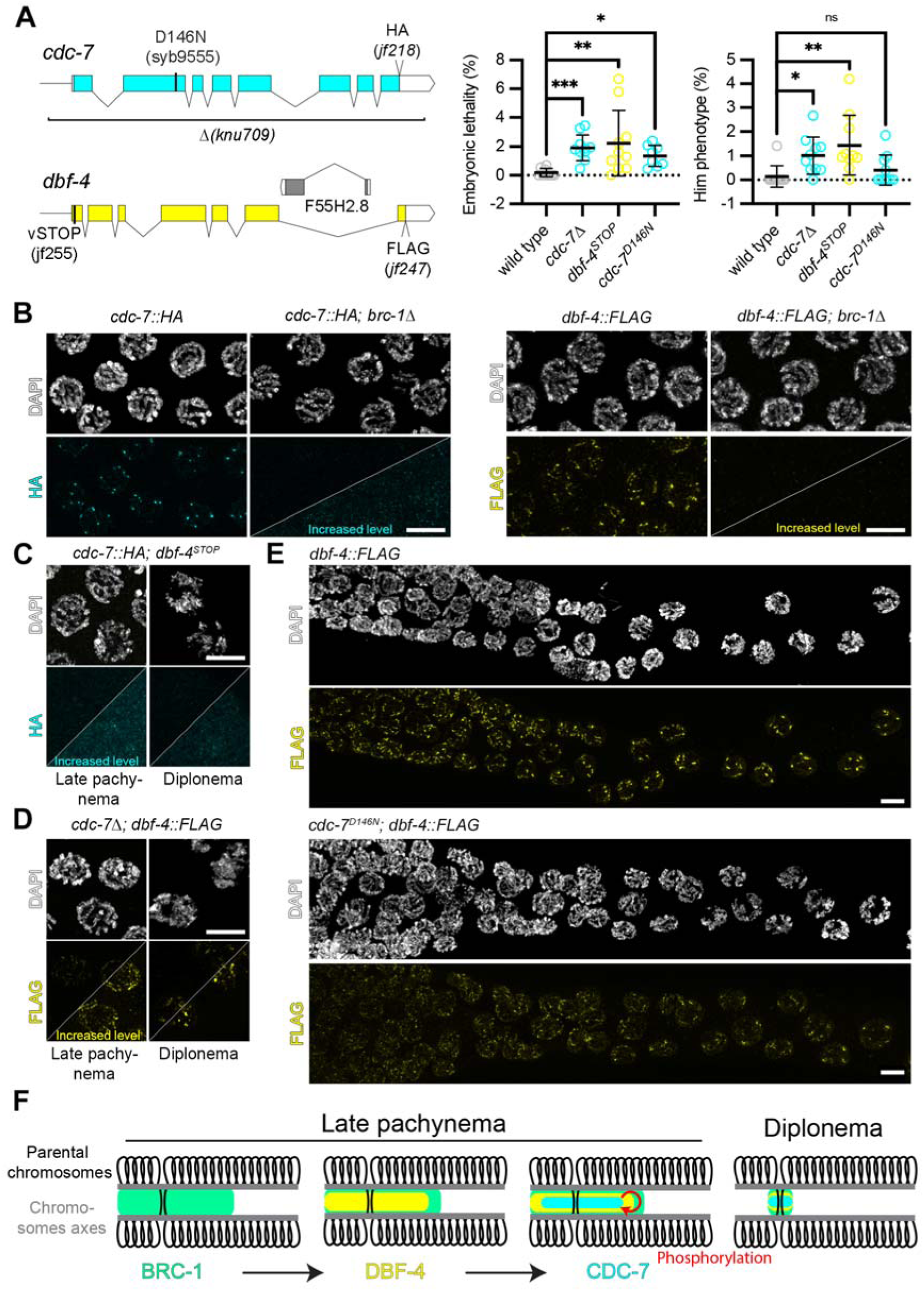
DDK mutants have subtle phenotypes, and DDK localization is hierarchical and BRC–1 dependent. **A.** Schematic representation of the *cdc*–*7* and *dbf*–*4* alleles used in this study, indicating *cdc*–*7*Δ (allele *knu709)*, catalytically dead *cdc*–*7^D146N^*(allele *syb9555*), and tagged *cdc*–*7::HA* (allele *jf218*). For *dbf*–*4*, the stop allele (*jf255*) and FLAG-tagged allele (*jf247*) are shown. Quantification of embryonic lethality and Him phenotype (%) for the indicated genotypes. Data are the mean ± SD. Statistical significance was determined using the Mann–Whitney U test (\**P* < 0.05, \*\**P* < 0.01, \*\*\**P* < 0.001; ns, not significant). **B.** Immunodetection of CDC–7::HA (cyan) and DBF–4::FLAG (yellow) with DAPI counterstaining (white) in the wild-type and *brc*–*1*Δ backgrounds during late pachynema. Lower-diagonal panels show increased signal intensity for HA and FLAG immunodetection. Scale bar, 5 µm. **C.** Immunodetection of CDC–7::HA (cyan) in the *dbf*–*4^STOP^* mutant at the specified stages. Lower-diagonal panels show increased signal intensity for HA immunodetection. Scale bar, 5 µm. **D.** Immunodetection of DBF–4::FLAG (yellow) with DAPI counterstaining (white) in the *cdc*–*7*Δ mutant for the specified stages. Lower-diagonal panels show increased signal intensity for HA immunodetection. Scale bar, 5 µm. **E.** Immunodetection of DBF–4::FLAG (yellow) during late pachynema to early diplonema in wild-type and *cdc*–*7^D146N^* backgrounds. Scale bar, 5 µm. **F.** Model illustrating the hierarchical loading of BRC–1, DBF–4, and CDC–7 as foci and signaling along chromosome axes during late pachynema and diplonema.

Additionally, we confirmed that CDC–7 contains the required residues for a kinase (KUNG AND JURA 2019; KWON *et al*. 2019): (1) a lysine within the N-terminal lobe (VAIK motif), (2) the catalytic aspartate (HRD motif), and (3) an aspartate required for metal-ion binding (DFG motif) (Figure S2A,B). We mutated the catalytic aspartate to asparagine (D146N) using CRISPR–CAS9 to create a putative catalytically inactive CDC–7 (allele *syb9555*, hereafter *cdc*–*7^D146N^*) (Figure 2A).

The *cdc*–*7^D146N^*mutant displayed similar levels of embryonic lethality to both the *cdc*–*7*Δ and *dbf*–*4*^STOP^ alleles (Figure 2A). When assessing the subcellular localization of catalytically inactive CDC–7, we found that the immunofluorescence signal was slightly reduced compared with the wild type (Figure S2C). Subcellular fractionation followed by immunoblotting of CDC–7::HA and CDC–7^D146N^::HA showed a slight increase in catalytically inactive CDC–7 in the cytosolic fraction compared with wild type (Figure S2D); however, the mutant protein remained predominantly nuclear and was present at wild-type levels (also here, a slight shift to the nuclear soluble fraction can be observed). This is consistent with the mutant retaining structural integrity despite a potential loss of catalytic activity.

Together, these observations suggest that *cdc*–*7* and *dbf*–*4* mutants, including the *cdc*–*7^D146N^* allele, exhibit similar levels of embryonic lethality, consistent with roles for the encoded proteins within the same complex.

### DDK loading is hierarchical, with BRC–1 at the base

Both the immunoprecipitation and proximity labeling experiments (Figure 1A; Tables S2 and S3) identified the BRC–1–BRD-1 complex as a prominent CDC–7 interactor/proximity hit. *C. elegans* BRC–1 has a similar dynamic subcellular localization pattern in the gonad to the DDK complex: it is expressed throughout the germline and is concentrated to the synaptonemal complex from mid-pachynema onwards and is then concentrated to the short arm of paired bivalents (JANISIW *et al*. 2018; LI *et al*. 2018). Therefore, we aimed to decipher the relationship between DDK and the BRC–1–BRD-1 complex, as well as interdependencies between CDC–7 and DBF–4 regarding their respective subcellular localizations.

Immunodetection of either CDC–7::HA or DBF–4::FLAG revealed that BRC–1 is essential for subcellular localization, as no signal for either was detected in *brc*–*1*Δ (Figure 2B). CDC–7::HA signal was lost in the *dbf*–*4^STOP^*mutant, indicating that DBF–4 is necessary for CDC–7 localization (Figure 2C). In contrast, in *cdc*–*7*Δ germlines, DBF–4::FLAG staining was reduced but still detectable in prophase I, suggesting that CDC–7 might reinforce DBF–4 localization (Figure 2D). To further explore this relationship, we examined DBF–4 localization in the putative catalytically inactive *cdc*–*7^D146N^*mutant.

As in the deletion allele, DBF–4 signal was dimmer in the *cdc*–*7^D146N^*mutant and failed to form dense stretches in late pachynema (Figure 2E). Upon entry to diplonema, DBF–4 foci became visible in *cdc*–*7^D146N^*, but with reduced intensity compared with the wild type. This suggests that CDC–7 catalytic activity might be necessary for robust DBF–4 localization and for the formation of DBF–4 stretches in late pachynema and dense foci in diplonema, which coincides with retraction to the chromosome short arm of numerous chromosome axis and SC components (NABESHIMA *et al*. 2005; MARTINEZ-PEREZ *et al*. 2008; SATO-CARLTON *et al*. 2017; HURLOCK *et al*. 2020). Indeed, the CDC–7 signal in late pachynema and diplonema follows that of the SYP-1 synapsis protein, which retracts to the short arm of the forming bivalent (Figure S2F).

In summary, the hierarchy of protein loading starts with BRC–1, which is required for the loading of DBF–4, followed by CDC–7 (Figure 2F). In turn, kinase activity promotes stable loading of the complex; later, retraction of DDK to the short arm appears to be mediated by the kinase activity.

As the BRC–1–BRD-1 complex is necessary for chromosomal loading of the DDK complex, and both *brc*–*1* and *brd-1* mutants display similar low levels of embryonic lethality (LI *et al*. 2018) as in DDK complex mutants (Figure 2A), we wanted to learn whether DDK complex mutants are sensitive to exogenous DNA damage, as previously reported for *brc*–*1* mutants (BOULTON *et al*. 2004). To test this, we measured embryonic lethality in young hermaphrodites after gamma irradiation over 3 days. Embryos derived from irradiated meiocytes from late pachynema (day 1), early to late pachynema (day 2), and in the progenitor zone to zygonema (day 3; Figure S3A). In irradiated *brc*–*1* mutant worms, viability was significantly reduced (as reported by (BOULTON *et al*. 2004)); this was not at all the case for the DDK mutants (Figure S3B). Therefore, we conclude that, despite the requirement for BRC–1 for loading the DDK complex, *cdc*–*7* mutants do not phenocopy all defects observed in *brc*–*1* mutants. Thus, even if BRC–1 is required for DDK loading, the two factors have non-redundant functions in meiosis.

### SPO-11–dependent chromosome bridges arise at an early stage of meiotic recombination in *cdc*–*7* mutants

Consistent with the high hatch rates of the mutants, the end products of meiotic recombination largely consisted of six connected chromosome pairs (bivalents) (reflected by the six DAPI-positive chromosome bodies in diakinesis nuclei, as in the wild type). In *cdc*–*7*Δ, no change in the number of DAPI bodies at diakinesis was detected, whereas in both *cdc*–*7^D146N^* and *dbf*–*4^STOP^* mutants, we observed a small but significant reduction of DAPI bodies at this stage (Tables S5 and S6).

However, closer inspection of the putative bivalents revealed thin bridges at diakinesis in the *cdc*–*7*Δ*, cdc*–*7^D146N^*, and *dbf*–*4^STOP^*mutants (Figure 3A). These DNA bridges between DAPI bodies were present in approximately 15% of nuclei at diakinesis in the DDK mutants, but in only 0.83% of nuclei in the wild type (Figure 3B; *P* values are in Table S7). The high number of bridges at diakinesis is in stark contrast with the low embryonic lethality in these mutants (around 2%; Figure 2A). In animal cells, the LEM3/ANKLE1 nuclease is a last-resort nuclease that removes bridges during anaphase in mitosis and meiosis (HONG *et al*. 2018a; HONG *et al*. 2018b). Therefore, we wanted to determine whether this nuclease is responsible for removing the bridges and, thereby, contributing to the near-normal hatch rates in the mutants. Indeed, embryonic lethality increased in the *cdc*–*7*Δ*; lem-3* double mutant to 15% (Figure 3C; *P* values are in Table S8). We conclude that LEM-3 removes most of the bridges observed at diakinesis and, therefore, restores the viability of DDK mutants.

**Figure 3.**
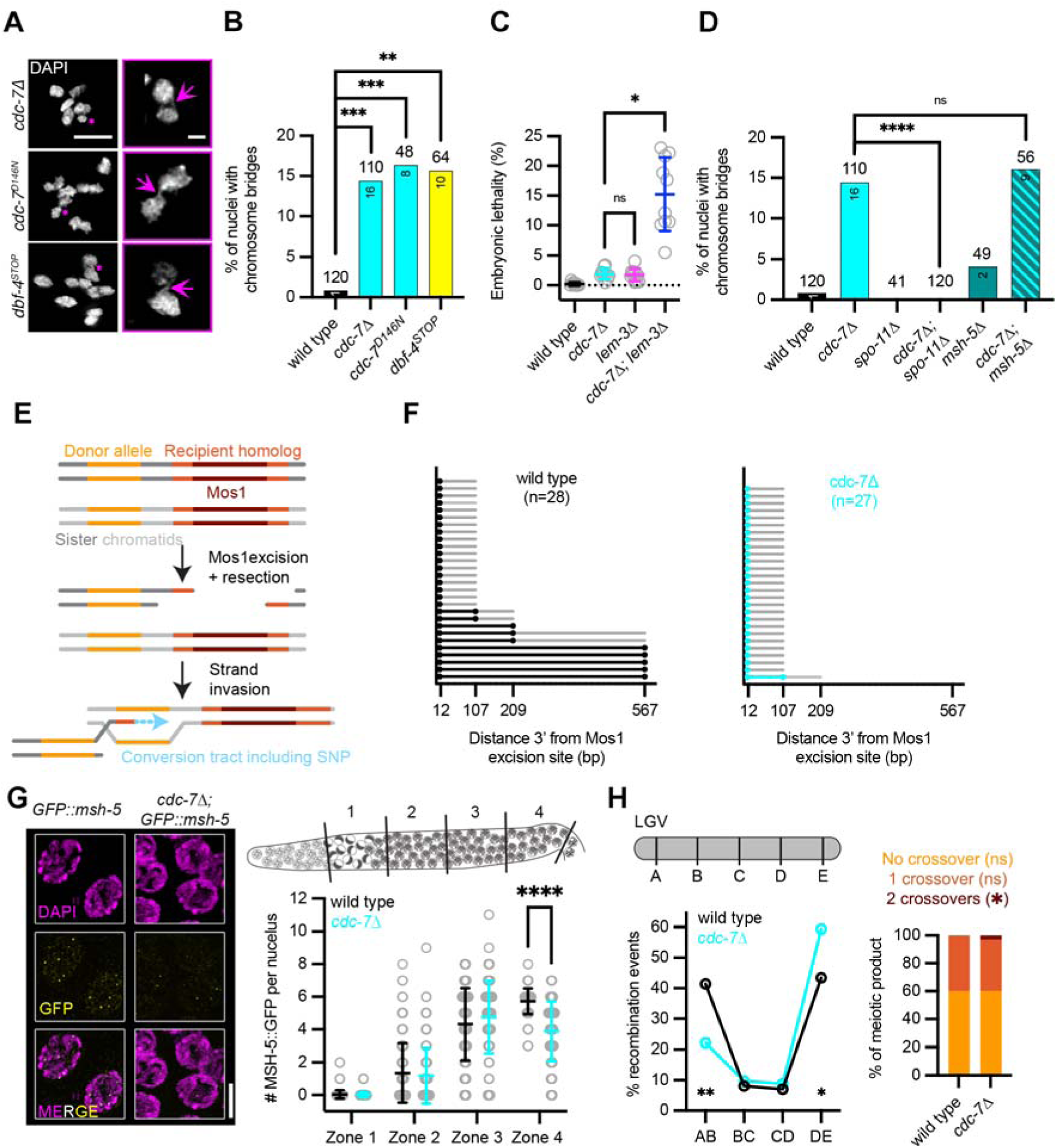
Chromosome bridges in *cdc*–*7*Δ are *spo-11* dependent and coincide with short DNA resection tracks with largely normal crossover recombination. **A.** DAPI-stained diakinesis nuclei of the indicated genotypes. An asterisk in the maximum intensity projection (left) marks the DAPI body shown in the inset. Insets highlight chromosome bridges (magenta arrows). Scale bar, 5 µm for main images and 1 µm for insets. **B.** Percentage of nuclei with chromosome bridges at diakinesis (−1 oocyte) for the indicated genotypes. Numbers of nuclei counted are indicated above the bars and numbers of bridges observed inside the bars. Statistical significance was determined using Fisher’s exact test (\*\**P* < 0.01, \*\*\**P* < 0.001). **C.** Embryonic lethality for the indicated genotypes. Data are the mean ± SD. Statistical significance was determined using the Mann–Whitney U test (\**P* < 0.05; ns, not significant). **D.** Percentage of nuclei with chromosome bridges at the −1 diakinesis for the indicated genotypes. Numbers of nuclei counted are indicated above the bars and numbers of bridges observed inside the bars. Statistical significance was determined using Fisher’s exact test (\*\*\*\**P* < 0.0001; ns, not significant). Counts for the wild type and cdc–7Δ in parts C and D are identical to those in part B. **E.** Schematic diagram of the Mos1 excision assay (TORAASON *et al*. 2021). Following Mos1 excision and resection, strand invasion into the homolog generates a conversion tract containing single nucleotides polymorphisms (SNPs). **F.** Distribution of conversion tract lengths, measured as the distance from the Mos1 excision site in base pair (bp) in the wild type and *cdc*–*7*Δ. **G.** Left, immunodetection of MSH-5::GFP (yellow) with DAPI counterstaining (magenta) in wild-type and *cdc*–*7*Δ animals. Right, number of MSH-5::GFP foci per nucleus detected across meiotic zones. Data are the mean ± SD. Statistical significance was determined using the Mann–Whitney U test (\*\*\*\**P* < 0.0001). Scale bar, 5 µm. **H.** Top left, genetic map of linkage group V (LGV), showing the intervals scored for recombination. Quantification of the recombination frequency (%) across intervals (bottom left) and distribution of crossover classes (right) in wild-type and *cdc*–*7*Δ animals. Statistical significance was determined using Fisher’s exact test (\**P* < 0.05; \*\**P* < 0.01).

We next investigated whether the bridges result from meiotic DSBs introduced by SPO-11 (DERNBURG *et al*. 1998). Analysis of the *spo-11*Δ*; cdc*–*7*Δ double mutant revealed that bridge formation at diakinesis is meiosis specific and requires SPO-11 activity (Figure 3D; DAPI body counts are in Table S6 and *P* values are in Table S9; *P* values for bridges are in Table S10). Surprisingly, bridges were present in the *cdc*–*7*Δ*; msh-5*Δ double mutant (Figure 3D), suggesting that they are formed before stabilization of the MSH-5-driven recombination intermediate (KELLY *et al*. 2000; YOKOO *et al*. 2012). Therefore, we reasoned that the bridge formation occurs upstream of MSH-5, and may include steps such as resection and D-loop formation or D-loop extension through DNA synthesis. To interfere with DNA synthesis, we depleted nucleotides by exposing worms to hydroxyurea (Figure S4A) and then examined bridges at diakinesis 42 hours (h) later, including in nuclei that had been in early pachynema during the treatment (Figure S4B). A strong reduction in bridge formation at diakinesis in treated *cdc*–*7*Δ mutants indicates that the bridges originate from a recombination intermediate upstream of D-loop extension (Figure S4C,D).

To explore the recombination steps upstream of strand invasion, we examined the loading of RPA-1 (ssDNA-binding factor) (WOLD 1997; HEFEL *et al*. 2021)) and RAD-51 (strand-exchange recombinase) (BAUMANN *et al*. 1996; ALPI *et al*. 2003; COLAIÁCOVO *et al*. 2003)) in the *cdc*–*7* mutant (Figure S4E). We found a general decrease in RPA-1 foci during meiotic progression (Figure S4F,G), which may reflect poor resection. The increased number of RAD-51 foci from middle to late pachynema (Figure S4H,I) may indicate the persistence or additional formation of recombination intermediates.

These findings suggest that chromosome bridges in *cdc*–*7* mutants arise from SPO-11-dependent recombination intermediates formed before the MSH-5 stabilization step. Those bridges are mainly resolved by LEM-3 during anaphase.

### DNA resection is impaired in *cdc*–*7* mutants despite largely preserved crossover outcomes

Reconstitution experiments suggesting a direct involvement of BRC1 in the resection process (Ceppi et al. 2024; Salunkhe et al. 2024), along with our results showing that (1) BRC–1–BRD-1 are in proximity to or form protein complexes with CDC–7 and (2) DDK localization depends on the BRC–1–BRD-1 complex prompted us to further evaluate whether resection is impaired in *cdc*–*7* mutants. We employed an assay that measures intersister/intrachromatid repair (ICR) (TORAASON *et al*. 2021) at an induced DSB generated by excision of the Mos1 transposable element at critical time windows in meiosis I prophase (Figure 3E). In this assay, Mos1 excision and resection around the DNA break enables strand invasion to occur on a strand donor allele that carries nucleotide polymorphisms. Detection of such an event and sequencing of the locus allow the extent of resection to be determined (as reflected by the extent of donor allele single-nucleotide polymorphism (SNP) incorporation). Notably, in cdc–7Δ worms, the ICR assay detected only short resection tracts, suggesting that the DDK complex supports resection (Figure 3F). This provides direct evidence that DDK promotes meiotic DNA end resection.

This result motivated us to measure the rate of successful recombination in the absence of CDC–7. We reasoned that if resection is defective, then crossovers might form less efficiently (as reported in yeast) (MOREAU *et al*. 1999; NEALE *et al*. 2005; HARTSUIKER *et al*. 2009; MILMAN *et al*. 2009; ZAKHARYEVICH *et al*. 2010; GARCIA *et al*. 2011; KEELAGHER *et al*. 2011; CANNAVO AND CEJKA 2014; DESHPANDE *et al*. 2016; MIMITOU *et al*. 2017). To test this hypothesis, we examined the localization of the pro-crossover factor MSH-5 (KELLY *et al*. 2000; YOKOO *et al*. 2012). In *cdc*–*7* mutants, we observed a significant reduction in the number of MSH-5 foci in late pachynema (wild type: 5.72 ± 0.79, *n* = 140 nuclei, *cdc*–*7*Δ: 3.90 ± 1.81, *n* = 126 nuclei, *P* = 3.63 × 10^−21^, Mann–Whitney U test with correction for multiple comparisons; Figure 3G, zone 4), despite the presence of a comparable number of MSH-5 foci in earlier stages. This suggests that in *cdc*–*7* mutants, crossover formation might be compromised or MSH-5 is less stable at crossover sites, as previously reported for the MutSγ complex in yeast, where Cdc7 masks a degron sequence in Msh4 (HE *et al*. 2020).

The reduced number of MSH-5 foci did not lead to a marked increase in univalent formation at diakinesis (Table S6). Therefore, we wanted to determine the extent of genetic exchange on chromosome V through a recombination assay (DAVIS *et al*. 2005; ROCKMAN AND KRUGLYAK 2009). For this, the *cdc*–*7* mutation was engineered into the closely related Hawaiian ecotype, and we measured recombination in the oocytes of heterozygous (Bristol/Hawaii) hermaphrodite *cdc*–*7*Δ animals (see Materials and Methods).

Overall, the recombination landscape remained largely similar to in the wild type, with one crossover per chromosome pair, enrichment on chromosome arms, and depletion in chromosome centers (ROCKMAN AND KRUGLYAK 2009; KAUR AND ROCKMAN 2014) (Figure 3H). Notably, we observed a shift in the position of crossovers toward the right arm of linkage group V, at the expense of those on the left arm (Figure 3H, left). The overall frequency of genetic exchange was comparable to in the wild type, although we detected seven double crossovers in the mutant (2.6% in *cdc*–*7*Δ; Figure 3H, right, Table S11).

In summary, our results show that CDC–7 promotes efficient DNA end resection during meiosis, as indicated by the shorter resection tracts in *cdc*–*7* mutants. Additionally, there are fewer MSH-5 foci in late pachynema, suggesting impaired stabilization of crossover intermediates. However, crossover frequency and distribution are mostly unchanged, indicating that crossover outcomes are resilient to reduced resection.

### CDC–7 works in conjunction with DNA–2 to regulate long-range DNA resection

DNA resection after DSB formation is essential for successful crossover formation. As in all organisms studied so far, in *C. elegans* short-range resection is mediated by the MRN complex, whereas long-range resection involves the exonucleases EXO-1 and DNA–2 (CHIN AND VILLENEUVE 2001; HICKS *et al*. 2022). Given that our previous results suggest that the DDK complex is involved in resection, we examined the diakinesis chromosomes in *exo-1* and *dna*–*2* mutants. For this, we employed a deletion allele of the *exo-1* gene and an auxin-induced degron *dna*–*2* allele (*dna*–*2^AID^*for the specific depletion of DNA–2 during meiosis (HICKS *et al*. 2022)). Consistent with our observations in *cdc*–*7* mutants, where we detected DNA bridges in diakinesis chromosomes (Figure 4A,B), we found fine DNA bridges in 30% of *exo-1* diakinesis nuclei and bridges in 54% of *dna*–*2^AID^* depleted nuclei (Figure 4C,D). The presence of bridges is consistent with a defect in long-range resection.

**Figure 4.**
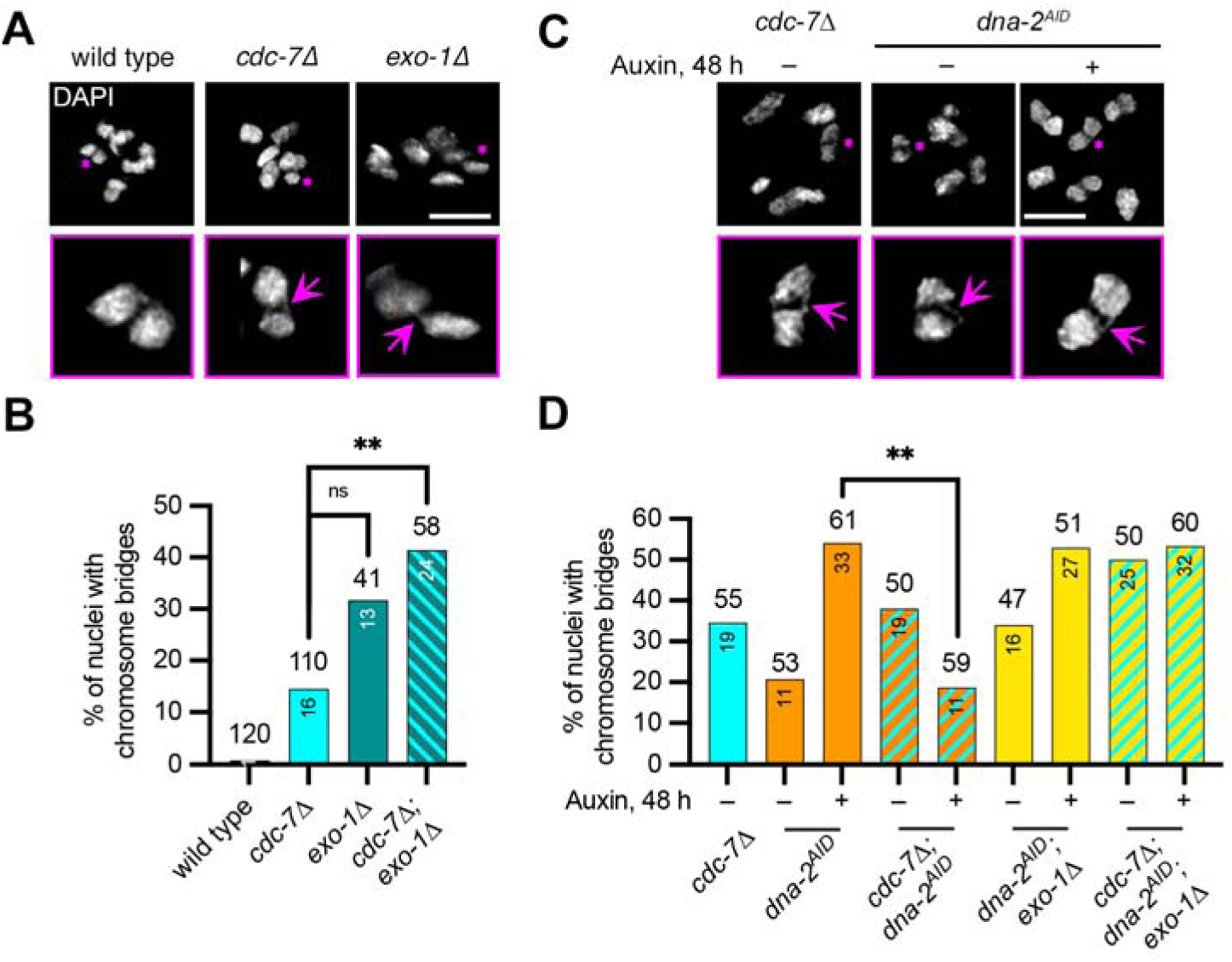
CDC–7 channels resection through DNA–2 while limiting EXO-1-dependent processing. **A.** DAPI-stained diakinesis nuclei for the indicated genotypes. An asterisk in the maximum intensity projection (top) marks the DAPI body shown in the inset (bottom). The *cdc*–*7*Δ DAPI body shown is identical to the one presented in Fig. 3A. Insets highlight chromosome bridges (magenta arrows). Scale bars, 5 µm for main images and 1 µm for insets. **B.** Percentage of nuclei with chromosome bridges at the −1 diakinesis for the indicated genotypes. Data for the wild type and *cdc*–*7*Δ are as shown in Fig. 3B. Numbers of nuclei counted are indicated above the bars and numbers of bridges observed inside the bars. Statistical significance was determined using Fisher’s exact test (\*\**P* < 0.01; ns, not significant). **C.** DAPI-stained diakinesis nuclei for the indicated genotypes without (−) or with (+) auxin depletion for 48 h post gravid stage. An asterisk in the maximum intensity projection (top) marks the DAPI body shown in the inset (bottom). Insets highlight chromosome bridges (magenta arrows). Scale bars, 5 µm for the main image and 1 µm for insets. **D.** Percentage of nuclei with chromosome bridges at the −1 diakinesis for the indicated genotypes without (−) or with (+) auxin depletion for 48 h post gravid stage. Numbers of nuclei counted are indicated above the bars and numbers of bridges observed inside the bars. Statistical significance was determined using Fisher’s exact test (\*\**P* < 0.01).

We then crossed the *exo-1*Δ allele into the *cdc*–*7*Δ mutant to test the genetic interaction of these mutations. In the double mutant *cdc*–*7*Δ*; exo-1*Δ, the number of bridges was significantly increased compared with in the single mutant *cdc*–*7*Δ, suggesting that CDC–7 and EXO-1 might operate in different pathways that contribute to efficient long-range resection (Figure 4B; diakinesis DAPI body counts are in Table S6 and *P* values are in Table S12; *P* values for the bridges are in Table S13).

Next, we combined the *dna*–*2^AID^* allele with *cdc*–*7*Δ. Surprisingly, in the *cdc*–*7*Δ mutant upon depletion of DNA–2 by the addition of auxin for 48 h post-L4 stage (which allows for exclusive protein depletion during meiotic prophase), the formation of bridges was significantly reduced (Figure 4D; DAPI body counts are in Table S6 and *P* values are in Table S14; *P* values for the bridges are in Table S15) and we did not observe an additive effect as observed for the combined *cdc*–*7; exo-1* mutant (Figure 4D). This result supports the view that in the absence of CDC–7 and DNA–2, EXO-1 might become the predominant long-range resection pathway and, thus, compensates for the lack of DNA–2. Since EXO-1 did not compensate for the lack of DNA–2, it appears that CDC–7 may inhibit EXO-1 or promote the dominance of the DNA–2 pathway. Together, these results place CDC–7 within the DNA–2-dependent branch of long-range resection.

To further test whether loss of CDC–7 might boost EXO-1 activity, we first confirmed that the *exo-1*Δ*; dna*–*2^AID^* double mutant displays the same level of bridges as the single *dna*–*2^AID^* mutant (Figure 4D). In the *cdc*–*7*Δ*; exo-1*Δ*; dna*–*2^AID^* triple mutant, the number of bridges was similar to levels seen in the *dna*–*2^AID^; exo-1* double mutant and *dna*–*2^AID^* single mutant, which display bridges in 50% of diakinesis nuclei (Figure 4D). Therefore, we concluded that the reduction of bridges in *dna*–*2; cdc*–*7* mutants is dependent on EXO-1. (Please note that the AID regime leads to the analysis of diakinesis of older hermaphrodites and that the number of bridges is already elevated in unchallenged conditions).

Briefly, we present evidence that chromosome bridges are found in long-range resection mutants as well as in *brc*–*1* mutants (Figure S3C,D) and suggest that they result from poor resection. Consistently, bridge levels are not increased in the *brc*–*1; cdc*–*7* double mutant (Figure S3C,D), supporting the view that BRC–1 is required for CDC–7 loading. Evidence from our epistasis experiments suggests that the DDK complex is involved in long-range resection and appears to inhibit EXO-1, while working in conjunction with DNA–2.

### Chromosome bridges in DDK mutants arise from alternative repair pathways and occur within and between bivalents

To further investigate the formation of bridges in the DDK mutant, we first asked whether the homologous chromosome is necessary for this. To this end, we combined the *cdc*–*7*Δ mutant with a deletion mutant of the central element of the synaptonemal complex (*syp-2*Δ), which abolishes synaptonemal complex extension and, therefore, prevents the homolog from being available for repair (Figure 5A; DAPI body counts are in Table S6 and *P* values are in Table S16; *P* values for the bridges are in Table S17). Bridges between the DAPI bodies of nuclei at diakinesis were still found in the *cdc*–*7*Δ*; syp-2*Δ double mutant and at a similar level (18%) as found in the *cdc*–*7*Δ single mutant (14%, Figure 5A). Therefore, we conclude that bridge formation does not require the homolog.

**Figure 5.**
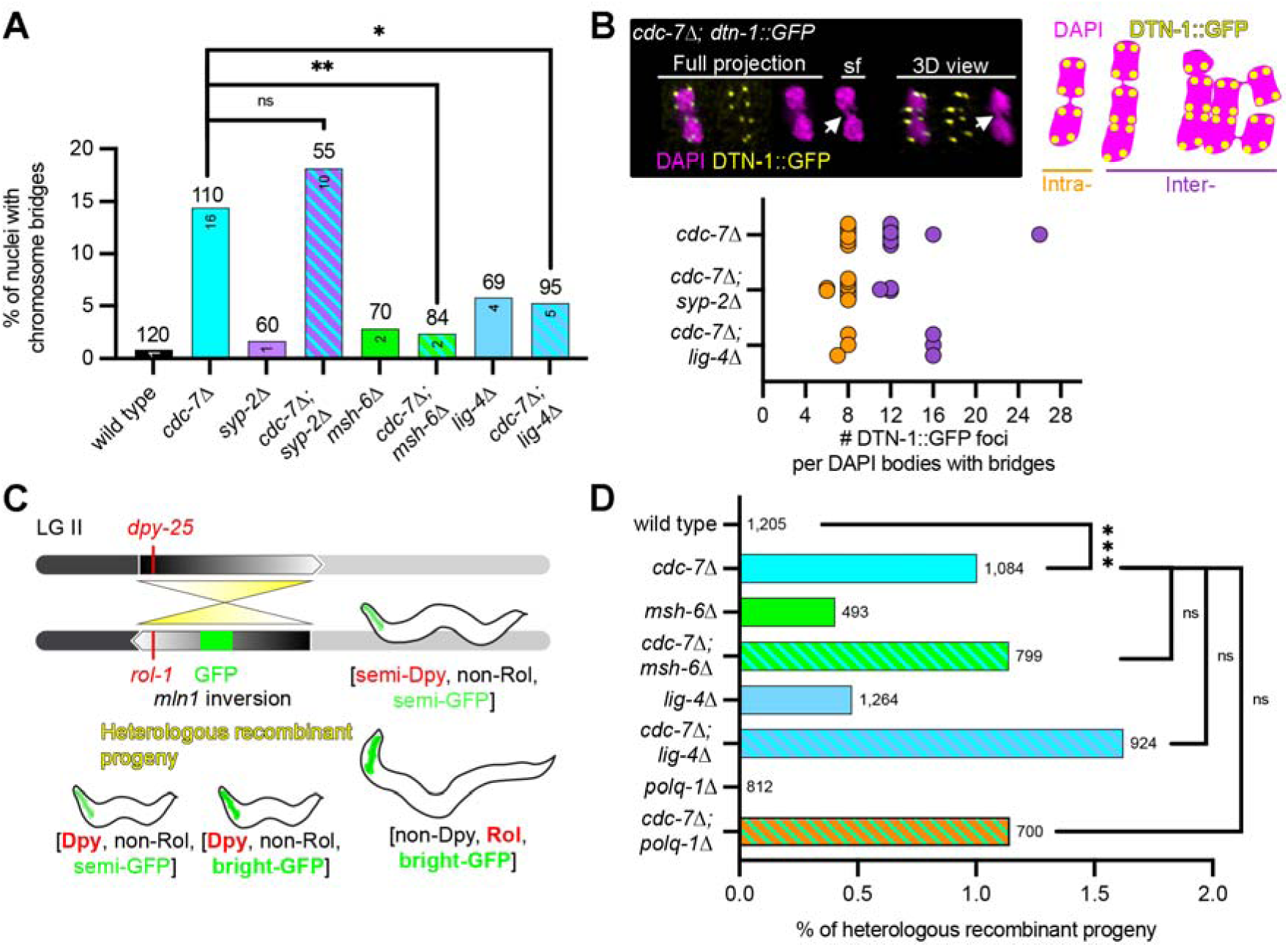
Mismatch repair factor MSH-6 and non-homologous end joining participate in bridge formation but do not account for elevated heterologous recombination in *cdc*–*7*Δ. **A.** Percentage of nuclei with chromosome bridges at the −1 diakinesis for the indicated genotypes. Numbers of nuclei counted are indicated above the bars and numbers of bridges observed inside the bars. Data for the wild type and *cdc*–*7*Δ are as shown in Fig. 3B. Statistical significance was determined using Fisher’s exact test (\**P* < 0.05, \*\**P* < 0.01; ns, not significant). **B.** Left, immunodetection of DTN-1::GFP (yellow) with DAPI counterstaining (magenta) in the *cdc*–*7*Δ mutant. Representative full projection, single focal plane (sf), and 3D views are shown. Right, schematic diagrams of possible classifications for intra- and inter-chromosomal foci. Bottom, quantification of DTN-1::GFP foci per DAPI body containing bridges for the indicated genotypes. **C.** Schematic diagram of the heterologous recombination assay on linkage group II (LG II) (LEÓN-ORTIZ *et al*. 2018). The *mIn1* inversion separates *dpy-25* and *rol-1* markers and enables detection of recombination events based on GFP intensity and marker segregation. Representative recombinant progeny classes are indicated. **D.** Percentage of heterologous recombination events for the indicated genotypes. Numbers next to bars indicate the number of progeny scored. Statistical significance was determined using a Chi-square test (\*\*\**P* < 0.001; ns, not significant).

This is consistent with our previous results that revealed that bridge formation does not depend on homologous recombination, since bridges also occur in *msh-5*Δ animals (Figure 3D). We next examined the contribution of alternative repair pathways to bridge formation. We found that both mismatch repair (*msh-6*Δ, (KRUISSELBRINK *et al*. 2008; LEÓN-ORTIZ *et al*. 2018)) and non-homologous end joining (*lig-4*Δ, (ADAMO *et al*. 2010)) are required because the *cdc*–*7*Δ*; msh-6*Δ and *cdc*–*7*Δ*; lig-4*Δ double mutants display a strong reduction in bridges between DAPI bodies in the −1 oocyte (2% and 5%, respectively; Figure 5A). Together, these results indicate that bridge formation between DAPI bodies in nuclei arises from alternative DNA repair pathways that act independently of homologous recombination, with both mismatch repair and non-homologous end joining contributing to the generation or stabilization of these structures.

To better understand the nature of these bridges, we used a strain expressing GFP-tagged DTN-1, a telomeric double-stranded DNA-binding protein (Yamamoto et al., 2021). At diakinesis, this marker appears as eight GFP-positive foci per DAPI body, corresponding to chromosome ends (Figure 5B, top left), which enabled us to distinguish bridges occurring within a single DAPI body representing a bivalent (8 DTN-1::GFP foci) from those connecting DAPI-positive structures representing more than one bivalent (> 8 DTN-1::GFP foci; Figure 5B, top right). In *cdc*–*7*Δ worms, we observed a nearly equal distribution of intra- and inter-bivalent body bridges: among 60 diakinesis nuclei analyzed, 14 (23%) displayed bridges, of which 6 (43%) were in inter-bivalent structures (Figure 5B, bottom).

Using the same DTN-1::GFP reporter assay, we next examined bridge-associated structures in *cdc*–*7*Δ*; syp-2*Δ diakinesis nuclei. In the absence of synapsis, individual univalents are expected to display four DTN-1::GFP foci corresponding to their chromosome ends. Of the 55 nuclei analyzed, 14 (25%) displayed bridges, with 11 of these bridge-associated structures (79%) containing at most eight DTN-1::GFP foci (Figure 5B, bottom), consistent with bridges involving at least one univalent-associated structure. Finally, in *cdc*–*7*Δ*; lig-4*Δ worms, the bridge frequency was strongly reduced (6 out of 107 nuclei; 6%), and the remaining bridge-associated structures showed no obvious bias in DTN-1::GFP focus distribution (Figure 5B, bottom).

Together, these results indicate that bridge formation is not restricted to a specific chromosomal context. Rather, they can occur between chromatids belonging to a bivalent and also between chromatids belonging to multiple bivalents.

### Heterologous recombination is increased in DDK mutants

Increased heterologous recombination has been reported in *brc*–*1* mutants (LEÓN-ORTIZ *et al*. 2018). Since DDK loading requires the BRC–1–BRD-1 complex, we asked whether a similar defect could exist in DDK mutants. Therefore, we assessed heterologous recombination in the *cdc*–*7*Δ mutant using the assay established by (LEÓN-ORTIZ *et al*. 2018) (Figure 5C). Whereas no heterologous recombination events were detected in the wild type, we observed a significant increase in the *cdc*–*7*Δ mutant (1.0%; Figure 5D; *P* values are in Table S17). We next asked whether the alternative repair pathways implicated in bridge formation also contribute to increased heterologous recombination in *cdc*–*7*Δ. To address this, we introduced the *msh-6*Δ and *lig-4*Δ alleles into the reporter system, with or without *cdc*–*7*Δ. Levels of heterologous recombination in both *cdc*–*7*Δ*; msh-6*Δ and *cdc*–*7*Δ*; lig-4*Δ double mutants were comparable to those observed in the *cdc*–*7*Δ single mutant (1.1% and 1.6% respectively; *P* values are in Table S18). Since alternative end joining is a frequently employed repair route in the germline (KAMP *et al*. 2020), we also tested whether the non-homologous recombination events are POLQ-1-dependent (Figure 5D). Levels of heterologous recombination in the *cdc*–*7*Δ*; polq-1*Δ double mutant were comparable to those in the *cdc*–*7*Δ single mutant (1.14% and 1.0%, respectively; Figure 5D; *P* values are in Table S18), indicating that the increase in heterologous recombination observed upon loss of CDC–7 is independent of POLQ-1. These results indicate that heterologous recombination is elevated in *cdc*–*7* mutants but is independent of mismatch repair, non-homologous end joining, and alternative end joining. This suggests that these chromosome translocations are not derived from the unscheduled use of alternative repair pathways, unlike the bridges we observed in the *cdc*–*7* mutants.

### Phosphorylated CDK–12 regulates long-range resection and suppresses the chromosome aberrations in *cdc*–*7* mutants

We found a 3.7-fold reduction in the CDK–12 Ser30 phosphopeptide in *cdc*–*7* mutants compared with the wild type (Figure 1D). In *C. elegans*, CDK–12 is best known as the major germline RNA polymerase II C-terminal domain (CTD) Ser2 kinase and as a regulator of genome stability through transcriptional control of DNA damage response genes (BOWMAN *et al*. 2013; SARKAR *et al*. 2023). (For experimental details of CDK–12 activity in the context of RNA polymerase II and for subcellular localization of CDK–12, see supplemental Figure S5 and Supplemental Text.) Therefore, we hypothesized that during meiosis in *C. elegans*, the DDK complex promotes resection through a CDK–12/DNA–2 signaling axis. To test this model, we generated phosphomutant alleles of CDK–12 at Ser30. A non-phosphorylatable variant allele (*syb9527*, hereafter *cdk*–*12^S30A^*) was obtained by alanine substitution, whereas a phosphomimetic variant allele (*syb9536*, hereafter *cdk*–*12^S30D^*) was generated by aspartic acid substitution.

We first examined bridge formation in diakinesis chromosomes in the *cdk*–*12^S30A^*mutant: *cdk*–*12^S30A^*animals displayed levels of bridges comparable to those observed in the *cdc*–*7*Δ mutant (15.0% and 14.5%, respectively), and no additive effect was detected in the *cdc*–*7*Δ*; cdk*–*12^S30A^*double mutant (16.2%, Figure 6A; DAPI body counts are in Table S6 and *P* values are in Table S19; *P* values for bridges are in Table S20). These results indicate that CDC–7 and CDK–12 act in the same pathway to regulate bridge formation.

**Figure 6.**
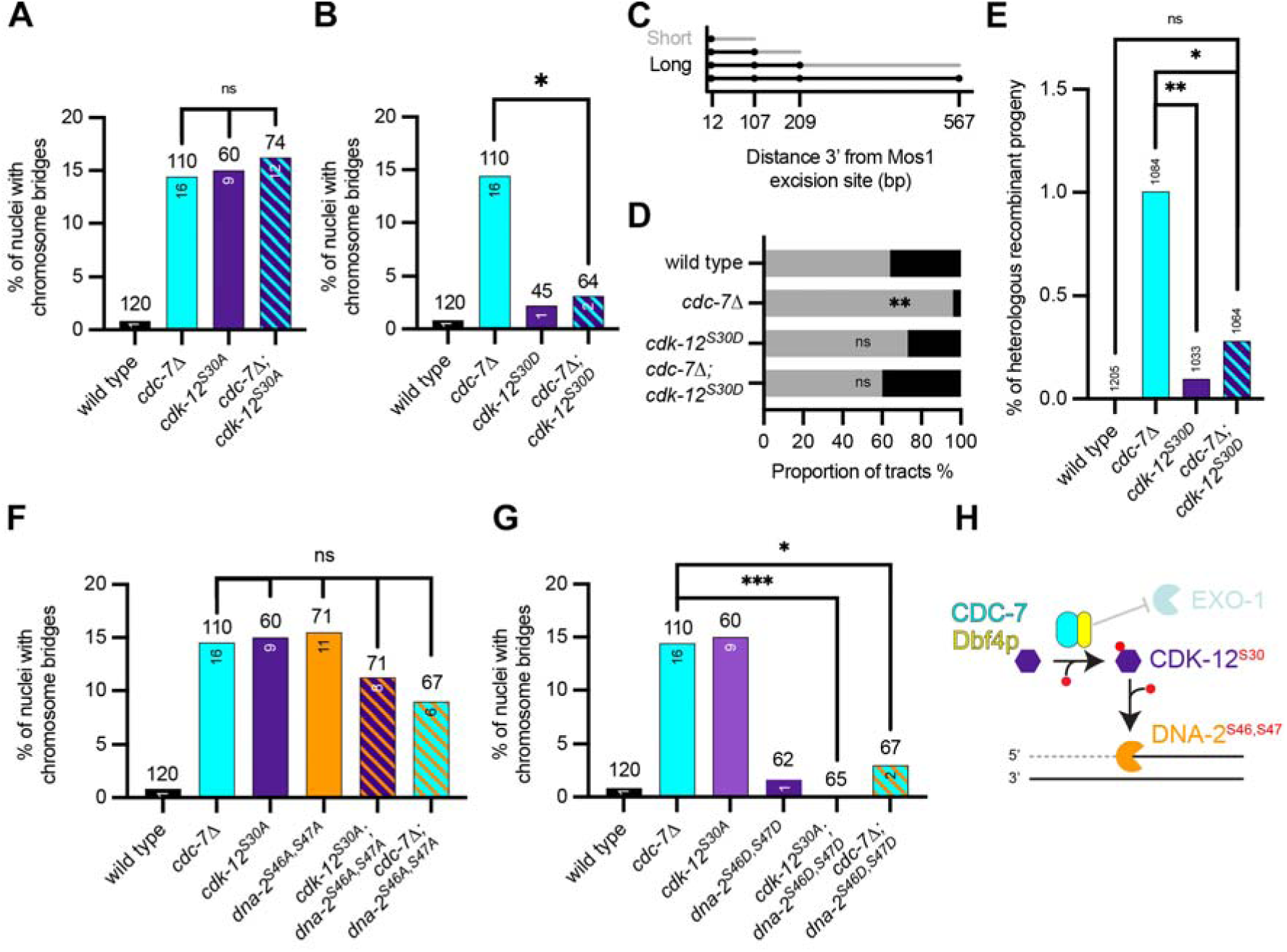
CDK–12 and DNA–2 act downstream of CDC–7 to promote resection and limit aberrant recombination outcomes. **A.** Percentage of nuclei with chromosome bridges at the −1 diakinesis for the indicated genotypes. Statistical significance was determined using Fisher’s exact test (ns, not significant). Data for the wild type and *cdc*–*7*Δ are as shown in Fig. 3B. **B.** Percentage of nuclei with chromosome bridges at the −1 diakinesis for the indicated genotypes. Numbers of nuclei counted are indicated above the bars and numbers of bridges observed inside the bars. Statistical significance was determined using Fisher’s exact test (\**P* < 0.05). Data for the wild type and *cdc*–*7*Δ are as shown in Fig. 3B. **C.** Schematic diagram defining classification of conversion tracts as short or long based on distance from the Mos1 excision site (bp). **D.** Proportion of short and long conversion tracts for the indicated genotypes. Statistical significance was determined using Fisher’s exact test (\*\**P* < 0.01; ns, not significant). **E.** Frequency of heterologous recombination for the indicated genotypes. Statistical significance was determined using a Chi-square test (\**P* < 0.05, \*\**P* < 0.01; ns, not significant). Data for the wild type and *cdc*–*7*Δ are as shown in Fig. 5B. **F.** Percentage of nuclei with chromosome bridges at the −1 diakinesis for the indicated genotypes. Numbers of nuclei counted are indicated above the bars and numbers of bridges observed inside the bars. Statistical significance was determined using Fisher’s exact test (ns, not significant). Data for the wild type and *cdc*–*7*Δ are as shown in Fig. 3B. **G.** Percentage of nuclei with chromosome bridges at the −1 diakinesis for the indicated genotypes. Numbers of nuclei counted are indicated above the bars and numbers of bridges observed inside the bars. Statistical significance was determined using Fisher’s exact test (\**P* < 0.05, \*\*\**P* < 0.001). Data for the wild type and *cdc*–*7*Δ are as shown in Fig. 3B. **H.** Model illustrating a CDC–7/CDK–12/DNA–2 regulatory axis. CDC–7 promotes CDK–12 phosphorylation, which, in turn, activates DNA–2 to drive long-range resection while limiting EXO-1-dependent processing.

We next analyzed the phosphomimetic *cdk*–*12^S30D^*allele. In *cdk*–*12^S30D^*worms, bridge formation at diakinesis was not significantly different from in the wild type (2.2% and 0.8%, respectively, Figure 6B; DAPI body counts are in Table S6 and *P* values are in Table S21; *P* values for bridges are in Table S22). Notably, in the *cdc*–*7*Δ*; cdk*–*12^S30D^* double mutant, bridge formation was strongly reduced, with numbers indistinguishable from in the wild type (3.1%, Figure 6B; *P* values for bridges are in Table S22). These results indicate that CDK–12 phosphorylation is sufficient to suppress the bridge formation phenotype of *cdc*–*7*Δ mutants, consistent with CDK–12 acting downstream of DDK.

To further corroborate this, we introduced the *cdk*–*12^S30D^* allele into the ICR reporter strain to assess its impact on DNA resection. Based on the previously described assay (TORAASON *et al*. 2021) (Figure 3E), reads were categorized as short or long according to the length of conversion tracts (Figure 6C). In the *cdc*–*7*Δ mutant, the proportion of long conversion tracts was significantly reduced compared with the wild type (4% vs 36%, Figure 6D; Fisher’s exact test, *P* = 0.0052). In contrast, both the *cdk*–*12^S30D^*single mutant and, more prominently, the *cdc*–*7*Δ*; cdk*–*12^S30D^*double mutant displayed proportions of long conversion tracts comparable to the wild type (27% and 40%, respectively, Figure 6D; Fisher’s exact test, *P* = 0.2101 and *P* = 0.5722, respectively). These results indicate that loss of CDK–12 phosphorylation underlies the resection defect observed in *cdc*–*7* mutants.

We next revisited the increase in heterologous recombination observed in the *cdc*–*7* mutant and reasoned that its origin may lie upstream in defective resection. Therefore, we measured heterologous recombination in *cdk*–*12^S30D^*and *cdc*–*7*Δ*; cdk*–*12^S30D^*mutants. In both cases, heterologous recombination was significantly reduced compared with the *cdc*–*7*Δ mutant (0.09% and 0.28%, respectively, vs 1.00% in *cdc*–*7*Δ, Figure 6E; *P* values are in Table S23). Together, these results indicate that the increase in heterologous recombination observed in the *cdc*–*7* mutant is due to defective CDK–12 phosphorylation.

In summary, these findings demonstrate that constitutive CDK–12 phosphorylation restores long-range resection and suppresses elevated heterologous recombination in the *cdc*–*7* mutant.

### A DDK/CDK**–**12/DNA**–**2 signaling axis promotes long-range resection

We next tested whether CDK–12 acts upstream of DNA–2, potentially through direct phosphorylation. In budding yeast mitosis, DDK and CDK phosphorylate Dna2 at multiple residues, including S246 and S247 ((GALANTI *et al*. 2024); Figure S6A). In an independent proteome-wide phosphoscreen, we identified three phosphorylation sites on *C. elegans* DNA–2: S46, S47, and T383 (Figure S6B). The phosphopeptide containing the N-terminal sites S46 and S47 was identified with high confidence (PEP = 1.3 x 10^-21^; Table S24), but the exact site localization (S46 versus S47) could not be unambiguously resolved. T383 was identified with lower confidence (PEP = 3.36 x 10^-3^; Table S24). Sequence comparison revealed that both yeast Dna2 phosphosites S246 and S247 and *C. elegans* DNA–2 phosphosites S46 and S47 lie within a conserved SSP motif (Figure S6C).

Therefore, we generated a phosphomutant allele targeting S46 and S47 (*jf285*, termed *dna*–*2^S46A, S47A^*) and a phosphomimetic allele (*jf286*, termed *dna*–*2^S46D, S47D^*). In the *dna*–*2^S46A, S47A^* mutant, bridge levels (15.5%) were comparable to those observed in the *cdc*–*7*Δ and *cdk*–*12^S30A^*mutants (14.5% and 15.0%, respectively) and significantly elevated relative to the wild type (Figure 6F; DAPI body counts are in Table S6 and *P* values are in Table S25; *P* values for bridges are in Table S26). No additive effects were observed in *cdk*–*12^S30A^* and *dna*–*2^S46A, S47A^* single mutants or the *cdc*–*7*Δ*; dna*–*2^S46A, S47A^* double mutant, indicating that these factors may act in a linear pathway.

In contrast, no bridges were detected in the *dna*–*2^S46D, S47D^* mutant (Figure 6G). Moreover, the phosphomimetic *dna*–*2^S46D, S47D^* allele suppressed bridge formation in both the *cdk*–*12^S30A^*and *cdc*–*7*Δ backgrounds (2.5% and 0%, respectively, Figure 6G; DAPI body counts are in Table S6 and *P* values are in Table S27; *P* values for bridges are in Table S28). These results support a model in which DDK and CDK–12 act in a signaling cascade to phosphorylate DNA–2 and, thereby, promote long-range resection during meiosis.

Together, these findings define a DDK/CDK–12/DNA–2 signaling axis that promotes long-range resection during meiosis (Figure 6H). In this model, CDC–7-dependent phosphorylation of CDK–12 enables downstream phosphorylation of DNA–2 to drive efficient resection, while DDK concurrently limits EXO-1 activity, thereby ensuring proper pathway balance and preventing aberrant repair outcomes.

## Discussion

For our study, it proved advantageous that CDC–7 kinase is not essential for DNA replication initiation in worms because an alternative mechanism involving DNSN-1/MUS-101 exists (XIA *et al*. 2023). This enabled us to scrutinize DDK mutant phenotypes in meiosis. We identified the DDK complex as a regulator of efficient meiotic DNA end resection in *Caenorhabditis elegans*. CDC–7 and DBF–4 were found to form a functional complex that localized dynamically during prophase I and loaded hierarchically downstream of the BRC–1–BRD-1 complex. Loss of DDK function led to SPO-11-dependent chromosome bridges at diakinesis, arising from recombination intermediates that were formed before MSH-5-mediated recombination steps. These were largely resolved by the LEM-3/ANKL1 nuclease during anaphase. Consistent with defective resection, *cdc*–*7* mutants displayed shortened resection tracts and altered RPA-1 and RAD-51 dynamics. Genetic analyses placed CDC–7 within the DNA–2-dependent branch of long-range resection and indicated an inhibitory role for CDC–7 toward EXO-1. Chromosome bridge formation depended on non-homologous end joining and occurred both within and between bivalents, while other chromosome aberrations, such as those caused by heterologous recombination, were elevated independently of end-joining pathways. Phosphomimetic CDK–12 and DNA–2 mutants restored resection and suppressed aberrant recombination phenotypes in *cdc*–*7* mutants. Together, these findings support a model in which DDK promotes meiotic DNA end resection via a CDK–12/DNA–2 signaling axis that, when active, dominates EXO-1-mediated resection (Figure 6H).

Our work uncovers a previously unrecognized role for DDK in meiotic DNA end resection in animals. Our analysis places DDK upstream of long-range resection through CDK–12-dependent regulation of DNA–2. This is consistent with studies showing that in yeast Dna2 activity is modulated by phosphorylation and cooperates with CDK signaling to promote mitotic resection (CHEN *et al*. 2011; GALANTI *et al*. 2024). However, our findings define a distinct regulatory architecture in which DDK acts through CDK–12 to control DNA–2 during meiotic prophase. BRCA1–s involvement in DNA end resection has been demonstrated convincingly *in vitro* and *in vivo* (CEPPI *et al*. 2024; SALUNKHE *et al*. 2024; TORAASON *et al*. 2024). Here, we place BRC–1–BRD-1 at the base of DDK–CDK–DNA–2-mediated resection. Consistent with this, we detected as-yet unreported DNA bridges in *brc*–*1* mutants. It will be interesting to examine whether BRCA1 also plays a role in vertebrate meiotic resection.

Although *cdc*–*7* mutants displayed reduced resection tract length and fewer MSH-5 foci in late pachynema, crossover numbers and global recombination landscape are largely preserved. This suggests that meiotic recombination is sufficiently robust to moderate perturbations in resection, consistent with previous observations that crossover designation is tightly regulated and buffered to ensure obligate crossover formation (KIRKPATRICK *et al*. 2000; MARTINI *et al*. 2006; COLE *et al*. 2012) or that resection tracks in DDK mutants remain above a critical threshold to cause recombination failure (HUANG *et al*. 2025). One possible explanation is that only a subset of DSBs requires extensive resection to mature into crossovers, whereas others can proceed through alternative processing routes, although the risk for generating chromosome aberrations is markedly elevated. Poorly resected intermediates are at risk of forming translocations, as demonstrated by the loss of Exo1 and Sgs1 at an induced DSB in mitosis (KIMBLE *et al*. 2023). We have now identified DDK as a risk factor for translocation formation. Interestingly, DDK-dependent translocations were independent of end-joining mechanisms, suggesting that they might be generated by the canonical homologous recombination machinery. We can only infer this indirectly because, despite its sensitivity, the heterologous recombination assay we used is not suitable for mutants with inviable progeny (such as *spo-11*) or for class I crossover mutants (such as *msh-5* or *cosa-1*).

Several questions remain regarding how DDK regulates DNA end resection during meiosis. In budding yeast, long-range resection is largely driven by Exo1, whereas Dna2 contributes in parallel or under specific conditions (MIMITOU AND SYMINGTON 2008; CEJKA 2015). In contrast, our data suggest a distinct regulatory architecture in *C. elegans* in which DNA–2 plays a central role and EXO-1 activity is both redundant and somehow inhibited by DDK. How this inhibition is achieved at the molecular level remains unclear, and determining whether EXO-1 is directly or indirectly regulated by DDK-dependent signaling will require further investigation. In mouse meiotic resection, it is still not fully established which long-range resection nuclease is most important. Surprisingly, in the mouse, the MRN complex might not only start resection but also extend it through iterative nicking steps or alternatively by recruiting another resecting nuclease (KIM *et al*. 2025b), such as DNA2; so far, DDK involvement has not been shown.

Although our epistasis analyses support a model in which CDK–12 regulates DNA–2 activity, the direct phosphorylation events and their biochemical consequences remain to be defined. Finally, it will be important to determine how DDK-dependent resection is coordinated with crossover designation and whether similar regulatory mechanisms operate in other metazoans.

## Materials and Methods

### F55H2.7 sequence similarity search

F55H2.7 is an uncharacterized protein with no domain annotation in the UniProt database (accession: P34467, (THE UNIPROT 2023)). A NCBI blast search within the UniProt reference proteomes hit several nematode DBF4-type domain containing proteins, among these a protein from *Plectus sambesii* with a highly significant E-value of 1e-12 (UniProt:A0A914V5G5, region:28-286 (ALTSCHUL *et al*. 1997)). A blast search with the *Plectus sambesii* hit region identified numerous vertebrate and invertebrates DBF4 proteins with a significant E-value of 1e-4 and confirmed the phylogenetic relationship to the DBF4 protein family. Additional evidence was provided by a Foldseek search with the structure model of F55H2.7 derived from AphFoldDB, in which human DBF4 was the best hit outside the nematode homologs in the AFDB-proteome database with an E-value of 7.96e-6 (VARADI *et al*. 2022; VAN KEMPEN *et al*. 2024).

### Nematode strains, strain construction, and culture conditions

All strains listed are derivatives of N2 Bristol (Table S29) and were cultivated under normal conditions (BRENNER 1974) except for N2 Hawaii, which was used for the recombination test. CRISPR editing was done as previously described (PAIX *et al*. 2015). Guide and repair templates and genotyping primers are listed in Tables S30 and S31, respectively.

### Cytological preparation of gonads and immunostaining

Immunofluorescence was performed as previously described (MARTINEZ-PEREZ AND VILLENEUVE 2005). L4 hermaphrodites were incubated at 20°C for 16–20 h. Gonads were then dissected from young adults into 1× phosphate-buffered saline (PBS), fixed in 1% formaldehyde for 5 min at room temperature, and frozen in liquid nitrogen. After postfixation in ice-cold methanol, nonspecific binding sites were blocked by incubation in PBS containing 1% bovine serum albumin for at least 1 h. Antibodies were diluted in 1× PBST (1× PBS containing 0.1 % Tween 20) and incubated overnight at 4°C (for primary antibodies) or 2 h at room temperature (for secondary antibodies). After washing in PBST, samples were mounted in VECTASHIELD Antifade (Vector Laboratories Inc., Burlingame, CA; catalog number VECH-1000) containing 4L6-diamidino-2-phenylindole (DAPI; 2 mg/mL).

For analysis of diakinesis nuclei, gonads were prepared as described above and DAPI stained without antibody incubation. To visualize DTN-1::GFP in diakinesis oocytes, the same procedure was followed without primary or secondary antibody.

To visualize MSH-5::GFP, germlines were prepared as previously described (DELLO STRITTO *et al*. 2022). Briefly, 10–15 young adult worms were dissected on poly-L-lysine-coated slides in 1× egg buffer containing 0.1% Tween-20 and immediately frozen in liquid nitrogen. After freeze-cracking, slides were incubated in methanol at −20°C for 1 min and then fixed in 4% paraformaldehyde in 100 mM K_2_HPO_4_ (pH 7.4) for 15 min. Samples were then washed three times in 1× PBST and mounted with DAPI.

Primary and secondary antibodies are listed in Table S32.

### Microscopy and evaluation

Three-dimensional (3D) stacks of images were acquired using a DeltaVision Ultra High Resolution microscope equipped with 100×/1.40 oil-immersion objective lenses and a custom softWoRx software package. Images acquired with the DeltaVision were deconvolved using the softWoRx deconvolution algorithm. Maximum intensity projections of the deconvolved images were generated in ImageJ after adjusting the maxima and performing background subtraction with a rolling ball radius of 50 pixels. Where specified, images of gonads consist of multiple stitched images. This was necessary because of the field-of-view limitation at high magnifications. Manual stitching of images to build the entire gonads was performed in Adobe Photoshop. Fluorescence levels for stitched images were adjusted to each other in Adobe Photoshop to correct for the microscope’s auto-adjustment settings.

### Chromosome bridge quantification

Chromosome bridges were scored in 3D-deconvolved image stacks of −1 diakinesis nuclei. Image stacks were examined using ImageJ (SCHNEIDER *et al*. 2012) by scrolling through individual *z*-planes. To confirm bridge structures and assess their spatial organization, images were additionally visualized in three dimensions using Napari (SOFRONIEW *et al*. 2026).

### Image segmentation and intensity quantification

Nuclear signals were quantified using a custom Python pipeline based on Cellpose (STRINGER *et al*. 2021). The analysis was restricted to DAPI channel images, which were used for nucleus segmentation. The images were processed as either single-plane 2D images or RGB images, with the DAPI signal extracted from the blue channel and the corresponding signal of interest from the green channel. Prior to segmentation, images were converted to floating point and normalized using percentile-based scaling (1st–99th percentiles). Nuclei were segmented using the Cellpose generalist model with a fixed diameter of 35 pixels, flow threshold of 0.3, and cell probability threshold of 2.0. Segmentation masks were generated for each image and used to define regions of interest (ROIs). For each segmented nucleus, mean intensity values were extracted from the corresponding green channel image. DAPI and green images were paired based on their filename structure to ensure correct channel matching. Measurements were performed on a per-nucleus basis. The pixel size was set to 0.0642 µm in both the *x* and *y* dimensions and used for downstream spatial scaling where required. All measurements were exported as tabulated data, including ROI identifiers and associated intensity values, for downstream analysis. All parameters were kept constant across samples.

### Focus detection and data processing

For MSH-5 focus quantification, the gonad was divided into four zones of equal length, starting from early meiotic prophase (leptonema–zygonema), as defined by SUN-1(S8Pi) staining, and extending to the onset of diplonema. MSH-5 foci were manually counted in each nucleus. Data were plotted and statistical analyses were performed using GraphPad Prism (version 11.0).

### Embryonic viability assays

L4 worms were prepicked, incubated overnight at 20°C, and transferred to a fresh plate the next day and each of the following 3 days. Eggs and the number of hatched worms were counted. Embryonic viability (%) was calculated as (number of hatched eggs/total number of laid eggs) × 100. The high incidence of males (Him) phenotype was derived from the same dataset and calculated as (number of males/total number of hatched progeny) × 100. Unless specified otherwise, 10 worms per condition were used and data are presented as the mean viability ± SD.

### Nucleus isolation and protein fractionation from large *C. elegans* cultures

Nucleus isolation and cellular fractionation were done as previously described (SILVA *et al*. 2014). Briefly, large cultures of *C. elegans* were prepared by seeding 20 100-mm NGM plates with 1 mL OP50 bacteria (obtained by centrifuging a 2-L overnight *Echericia coli* culture and resuspending the pellet to a final volume of 40 mL). Between 5000 and 6000 *C. elegans* embryos were seeded onto each 100-mm plate and incubated at 20°C for 3 days. Young adult worms were collected and transferred to 50-mL tubes by washing the plates with M9 solution and pelleting the worms by gravity for 15 min. Most of the supernatant was then removed and fresh M9 solution added. This washing step was repeated three times. The final wash was performed using NP buffer (10 mM HEPES-KOH (pH 7.6), 1 mM EGTA, 10 mM KCl, 1.5 mM MgCl_2_, 0.25 mM sucrose, 1 mM phenylmethylsulfonyl fluoride, and 1 mM dithiothreitol (DTT)) containing protease inhibitors (Roche, catalog number 11836170001), and worms were pelleted by centrifugation at 600 *g* for 2 min.

Nuclei were isolated from a 1-mL sample of the worm pellet. Worms were disrupted using a cooled metal Wheaton tissue grinder, and the resulting suspension was filtered first through a 100-μm mesh and then through a 40-μm mesh. The filtered solution was centrifuged at 300 *g* for 2 min at 4°C, and the supernatant (containing nuclei) was centrifuged at 2500 *g* for 10 min at 4°C. The resulting supernatant was used as the cytosolic fraction, and the pellet as germline nuclear fraction. To separate soluble and DNA-bound nuclear protein fractions, we used a Qproteome Nuclear Protein Kit (QIAGEN, catalog number 37582) according to the manufacturer’s instructions.

### Native immunoprecipitation

At least 750 µg pooled soluble and insoluble nuclear fractions was used for immunoprecipitation. In parallel, extracts from wild-type worms were prepared and served as controls. CDC–7::HA was immunoprecipitated using anti-HA magnetic beads (Pierce, #88836). For all experiments (performed in triplicate), beads were pre-equilibrated in buffer D (20% glycerol, 0.2 mM EDTA pH 8.0, 150 mM KCl, 20 mM HEPES-KOH pH 7.9, 0.2% Triton X-100) supplemented with protease inhibitor cocktail (Roche). Protein extracts were incubated with the beads for 4 h at 4°C with rotation. Beads were then washed three times for 10 min in buffer D, followed by five additional washes in buffer D lacking Triton X-100. Proteins bound to the beads were analyzed by mass spectrometry.

### TurboID-based proximity labeling and streptavidin pull-down

Proximity labeling experiments were performed using TurboID-mediated biotinylation followed by streptavidin pull-down, as adapted from an established protocol (VELEZ-AGUILERA *et al*. 2024). Two strains were used: a strain expressing TurboID fused to CDC–7::GFP and a control strain expressing TurboID alone (see Table S29). Experiments were performed in biological triplicate. Nuclear fractions were prepared as described above. The nuclear pellet was resuspended in 800 µL lysis buffer (6 M guanidinium, 50 mM Tris-HCl pH 8.0, 100 mM NaCl, and 5 mM DTT, supplemented with protease inhibitor cocktail (Roche)) and homogenized by vortexing until fully solubilized. Samples were clarified by centrifugation at 13,000 *g* for 20 min at room temperature, and the supernatant was transferred to a fresh tube and centrifuged again under the same conditions. An aliquot was retained as the input control. Streptavidin magnetic beads (Pierce, #88817) were pre-equilibrated by washing three times in pre-wash buffer (50 mM Tris-HCl pH 7.5, 100 mM NaCl). Clarified lysates were incubated with 20 µL equilibrated beads for 1 h at room temperature with rotation. Following incubation, an aliquot of the supernatant was collected as the flow-through sample. Beads were washed sequentially to remove non-specifically bound proteins: twice rapidly and twice for 3 min in lysis buffer lacking DTT and protease inhibitors, followed by two washes in pre-wash buffer and two washes in sterile water. After washing, beads were resuspended in 350 µL water and subjected to on-bead tryptic digestion for mass spectrometry analysis.

### Preparation of frozen worm powder for phosphoproteomic analysis

For phosphomapping experiments, wild-type and *cdc*–*7*Δ worms were collected in biological triplicates from large-scale cultures as described above, and frozen in NP buffer. Frozen samples were thawed rapidly and immediately dispensed dropwise into liquid nitrogen to generate frozen worm beads. Beads were collected using a pre-chilled sieve, transferred to 15-mL tubes, and maintained in liquid nitrogen until grinding. Frozen worm beads were pulverized using a Freezer/Mill 6870 (SPEX SamplePrep) under liquid nitrogen. Grinding chambers, impactors, and plugs were pre-cooled before use. Samples were processed using five grinding cycles, each consisting of 1-min pre-cooling, 2-min run time, and 1-min cooling, at a cycle strength of 12. The resulting worm powder was transferred to pre-cooled 50-mL Falcon tubes and stored at −80°C until phosphoproteomic analysis. Three independent biological replicates were prepared for each genotype.

### Mass spectrometry analysis

#### On bead digest

Beads were resuspended in 60µl of 100 mM ammonium bicarbonate (ABC), supplemented with 600 ng of lysyl endopeptidase (Lys-C, Fujifilm Wako Pure Chemical Corporation) and incubated for 4 h on a Thermo-shaker with 1200 rpm at 37°C. The supernatant was transferred to a fresh tube and reduced with 2 mM Tris 2-carboxyethyl phosphine hydrochloride (TCEP, Sigma) for 30 min at 60°C and alkylated in 8 mM S-methyl methanethiosulfonate (MMTS, Fluka) for 30 min at room temp protected from light. Subsequently, the sample was digested with 600 ng Trypsin (Trypsin Gold, Promega) at 37°C over night. The digest was acidified by addition of 10 µl of 10% trifluoroacetic acid (TFA, Pierce). A similar aliquot of each sample was analyzed by LC-MS/MS.

#### nanoLC-MS/MS Analysis

The nano HPLC system (UltiMate 3000 RSLC nano system) was coupled to an Orbitrap Exploris 480 mass spectrometer, equipped with a FAIMS pro interface, equipped with a Nanospray Flex ion source (all parts Thermo Fisher Scientific).

Peptides were loaded onto a trap column (PepMap Acclaim C18, 5 mm × 300 μm ID, ìm particle size, 100 Å pore size, Thermo Scientific) at a flow rate of 25 μl/min using 0.1% TFA as mobile phase. After 10 min, the trap column was switched in line with the analytical column (PepMap Acclaim C18, 500 mm × 75 μm ID, 2 μm particles, 100 Å, Thermo Scientific). Peptides were eluted using a flow rate of 230 nl/min, starting with the mobile phases 98% A (0.1% formic acid in water) and 2% B (80% acetonitrile, 0.1% formic acid) and linearly increasing to 35% B over the next 120 min, followed by an increase to 95% B in 5 min, a 5-min hold at 95% B and ramping down in 2 min to the starting conditions of 98% A and 2% B for equilibration at 30°C.

The Orbitrap Exploris 480 mass spectrometer was operated in data-dependent mode, performing a full scan (m/z range 350-1200, resolution 60,000, normalized AGC target 100%) at 3 different compensation voltages (CV −45, −60, −75), followed each by MS/MS scans of the most abundant ions for a cycle time of 0.9 seconds (CV −45, −60) or 0.7 sec (CV −75). MS/MS spectra were acquired using HCD collision energy of 30, isolation width of 1.0 m/z, orbitrap resolution of 30.000, normalized AGC target 200%, minimum intensity of 25.000 and maximum injection time of 100 ms. Precursor ions selected for fragmentation (include charge state 2-6) were excluded for 45 s. The monoisotopic precursor selection filter and exclude isotopes feature were enabled.

#### Data Processing protocol

For peptide identification, the RAW files were loaded into Proteome Discoverer (version 2.5.0.400, Thermo Scientific). All MS/MS spectra were searched using MSAmanda v2.0.0.19924 (DORFER *et al*. 2014). The peptide mass tolerance was set to ±10 ppm and fragment mass tolerance to ±10 ppm, the maximum number of missed cleavages was set to 2, using tryptic enzymatic specificity without proline restriction. Peptide and protein identification was performed in two steps. For an initial search the RAW-files were searched against the uniprot_reference_*C._elegans*_2022-12-19.fasta database (19,834 sequences; 8,141,223 residue), supplemented with common contaminants and sequences of proteins of interest using beta-methylthiolation on cysteine as a fixed modification. The result was filtered to 1 % FDR on protein level using the Percolator algorithm (KÄLL *et al*. 2007) integrated in Proteome Discoverer. A sub-database of proteins identified in this search was generated for further processing. For the second search, the RAW files were searched against the created sub-database using the same settings as above and considering the following additional variable modifications: oxidation on methionine, deamidation on asparagine and glutamine, phosphorylation on serine, threonine, and tyrosine, glutamine to pyro-glutamate conversion at peptide N-terminal glutamine, and acetylation on protein N-terminus. The localization of post-translational modification sites within peptides was performed using the tool ptmRS, based on phosphoRS (TAUS *et al*. 2011). Identifications were filtered again to 1 % FDR on protein and PSM level, additionally, an Amanda score cut-off of at least 150 was applied. Proteins were filtered to be identified by at least 2 PSMs in at least 1 sample. Protein areas have been computed in IMP-apQuant (DOBLMANN *et al*. 2019) by summing up unique and razor peptides. Resulting protein areas were normalized using iBAQ,(SCHWANHÄUSSER *et al*. 2011), and sum normalization was applied to normalize across samples. Match-between-runs (MBR) was applied for peptides with high confident peak area that were identified by MS/MS spectra in at least one run. Statistical significance of differentially expressed proteins was determined using limma (SMYTH 2004).

#### Worm sample preparation and protein digestion for TMT-based proteomics

Frozen *C. elegans* worm powder was provided in twelve 50 mL Falcon tubes and stored at −80L°C prior to processing. Samples comprised three biological replicates each of wild type (N2), *cdc-7(knu709)*, *cdc-7(tm4391)* and *Dbf4p(stop)* strains, with 0.5 mg material assigned per TMTpro channel. Worm powder was lyophilized overnight and subsequently resuspended in 1 mL 50 mM HEPES buffer (pH 7.3). Proteins were extracted using methanol/chloroform precipitation by addition of 3 mL methanol, followed by incubation for 4 h at −20L°C, addition of 1 mL chloroform, and further incubation for 1 h at −20L°C. Samples were centrifuged for 10 min at 4,000 × g, the supernatant was removed, and protein pellets were stored at −20L°C.

Pellets were solubilized in 400 µL lysis buffer containing 100 mM DTT, 100 mM Tris and 4% SDS. Samples were sonicated for 20 min in a water bath followed by probe sonication (70% amplitude, 0.5 s pulse, 30 s duration) repeated three times.

Protein digestion was performed using a filter-aided sample preparation (FASP) workflow. Samples were loaded onto FASP filters in freshly prepared 8 M urea buffer (50 mM HEPES, pH 7.3) to minimize urea-derived peptide modifications.

Approximately 200 µg of protein per sample was loaded by repeated centrifugation at 14,000 × g for 20 min at room temperature. Filters were washed twice with 200 μL urea buffer, then alkylated with 100 mM iodoacetamide in 8 M urea buffer for 20 min in the dark at room temperature. Excess reagent was removed by repeated washing with urea buffer (2x) and subsequently with 50 mM HEPES buffer (pH 7.3)(3x). Proteins were digested overnight at 37L°C using 1:100 trypsin in 50 µL HEPES buffer. Peptides were recovered by centrifugation and additional washing steps, resulting in final peptide volumes of approximately 100 µL for subsequent TMT labeling.

#### TMTpro labeling and quality control analysis

TMTpro labeling was performed with up to 200 µg of peptide per sample. TMTpro reagents were equilibrated from −20L°C to room temperature immediately prior to use and dissolved in 20 µL anhydrous acetonitrile by repeated vortexing. Individual peptide samples were transferred directly into the corresponding TMTpro labeling reagent according to the predefined channel allocation. For samples containing excess peptide material, input amounts were normalized to 200 µg by adjusting sample and buffer volumes prior to labeling.

To assess labeling efficiency and reporter-ion bias, additional control samples were prepared, including a pooled reference sample generated by combining 5 µL from each labeled sample and selected individual samples for labeling-efficiency testing. An additional bias assessment was conducted to evaluate reporter-ion balance relative to label-free housekeeping protein abundances.

#### High-pH reversed-phase peptide fractionation

For high-pH reversed-phase fractionation, peptide samples corresponding to a total volume of 333.2 µL (∼334 µg peptide material) were first desalted using 50 mg Sep-Pak C18 cartridges (Waters) according to the manufacturer’s instructions. Following desalting, samples were concentrated in a SpeedVac concentrator until approximately 10% acetonitrile remained, then lyophilized. Dried peptides were resuspended in 75 µL 10 mM ammonium formate, and the complete sample volume was injected for fractionation.

High-pH fractionation was performed using an XBridge Peptide BEH C18 column (130 Å pore size, 3.5 µm particle size, 4.6 × 250 mm; Waters, part no. 186003570). Buffer A consisted of 10 mM ammonium formate, and buffer B of 90% acetonitrile. Peptides were separated using a gradient from 4.5% to 45% acetonitrile under high-pH conditions. A total of 40 fractions were collected and concatenated into 20 pooled whole-proteome fractions using the following pooling scheme: 1–20, 2–21, 3–22, 4–23, 5–24, 6–25, 7–26, 8–27, 9–28, 10–29, 11–30, 12–31, 13–32, 14–33, 15–34, 16–35, 17–36, 18–37 and 19–38. Fractions were subsequently concentrated in a SpeedVac concentrator and lyophilized. Prior to LC–MS analysis, fractions were resuspended in 50 µL of 0.1% TFA and and a small aliquot analyzed on an HPLC-UV system for quality control. LC-MS analysis was performed prior to phospho-peptide enrichment for whole-proteome-level normalization.

#### Phosphopeptide enrichment

Phosphopeptide enrichment was performed using the High-Select Fe-NTA Magnetic Phosphopeptide Enrichment Kit (Thermo Fisher Scientific; cat. no. A52283/A52284). The 20 high-pH reversed-phase fractions were first concatenated into three final pools: pool 1 (fractions 1, 3, 5, and 6), pool 2 (fractions 2, 7, and 8), and pool 3 (fractions 4, 9, and 10). Pooled fractions were concentrated in a SpeedVac concentrator and resuspended in 50 µL loading buffer consisting of 80% acetonitrile and 0.1% TFA. Fe-NTA magnetic beads (10 µL per sample) were equilibrated according to the manufacturer’s instructions before peptide samples were added. Binding was performed for 30 min at room temperature using overhead rotation. Following incubation, flow-through fractions were collected, and beads were washed three times with 20 µL binding/wash buffer and once with 20 µL MS-grade water. Phosphopeptides were eluted twice with 20 µL elution buffer containing 5% ammonium hydroxide. Eluates were concentrated to dryness in a SpeedVac concentrator and resuspended in 20 µL 0.1% TFA prior to LC–MS analysis.

#### Data-dependent acquisitions (DDA)

LC-MS/MS analysis was performed using an Orbitrap Exploris 480 with high-field asymmetric ion mobility spectrometry (FAIMS) interface (FAIMS Pro, Thermo Fisher Scientific, Waltham, Massachusetts, United States) coupled with an EASY-Spray source and Vanquish Neo UHPLC system (Thermo Fisher Scientific). A trap column PepMap C18 (5 mm × 300 μm ID, 5 μm particles, 100 Å pore size) (Thermo Fisher Scientific, Waltham, Massachusetts, United States) and an analytical column PepMap (length 50cm, diameter 75 μm, particle size 3 μm) (Thermo Fisher Scientific) were employed for separation. The column temperature was set to 50°C. Sample loading was performed using 0.1% trifluoroacetic acid in water with a flow rate of 25 uL/min. Mobile phases used for separation were as follows: A 0.1% formic acid (FA) in water; B 80% acetonitrile, 0.1% FA in water. Peptides were eluted using a flow rate of 230 nL/min, with the following gradient: from 2% to 35% phase B in 180 min, from 35% to 95% phase B in 1 min, followed by a washing step at 95% for min, and re-equilibration of the column.

Full MS scans were acquired in the Orbitrap at 60,000 resolution across an m/z range of 350–1,200 using profile mode, a normalized AGC target of 300%, and automatic maximum injection time settings. FAIMS compensation voltages (CVs) of −55 V and −70 V were applied in separate experiments. Precursor selection was performed using peptide mode with an intensity threshold of 25,000 and charge-state filtering restricted to precursor ions carrying charges between +2 and +6. Dynamic exclusion was enabled with exclusion after one scan for 45 s using a ±10 ppm exclusion window. Isotope exclusion and single charge-state precursor selection were enabled. MS/MS spectra were acquired using HCD fragmentation with normalized collision energy of 34%. Precursors were isolated using a 0.7 m/z isolation window and analyzed in the Orbitrap at 45,000 resolution. Fragment spectra were acquired from m/z 110 using centroid mode, a normalized AGC target of 200%, and a fixed maximum injection time of 200 ms. Data-dependent acquisition was operated in cycle-time mode with a total cycle time of 1 s.

#### Quantitative phosphoproteomic data processing and statistical analysis

Further computational analysis was performed using Python and the in-house developed Python library MsReport (0.0.29) (HOLLENSTEIN 2026). Protein-normalized phosphopeptide intensities were normalized across samples using the ModeNormalizer from MsReport to compensate for global effects not corrected by the protein normalization. The ModeNormalizer method involves calculating log2 protein ratios for all pairs of samples and determining normalization factors based on the modes of all ratio distributions. Statistical analysis was performed using the linear models for microarray analysis (limma) v.3.54.2 (RITCHIE *et al*. 2015) package in R. Moderated *t*-statistics were calculated using the limma-trend method, and multiple testing correction was applied using the Benjamini–Hochberg method. The Python library XlsxReport (0.1.0) (HOLLENSTEIN AND HARTL 2025) was used to create formatted Excel files summarizing the results of the proteomics experiments (Table S4). Raw mass spectrometry data and processed phosphoproteomic datasets have been deposited to the ProteomeXchange Consortium via the PRIDE (PEREZ-RIVEROL *et al*. 2025) partner repository under the dataset identifier PXD079177.

### Auxin depletion

For auxin depletion experiments, worms were prepicked at the L4 stage and grown at 20°C for 16–20 h. They were then transferred to NGM plates containing ethanol (wild type) or 4 mM 3-indoleacetic acid (auxin; Sigma-Aldrich, catalog number I2886) and grown for 48 h.

### Homologous recombination assay

Meiotic recombination frequencies and locations were determined using unique SNPs, as previously described (DAVIS *et al*. 2005; ROCKMAN AND KRUGLYAK 2009). CRISPR/Ca9 was used to generate the *cdc*–*7(knu709)* deletion in the CB4856 Hawaiian background *cdc*–*7(jf262).* Subsequently, mutants and controls were crossed to produce a hybrid, as previously described (VELKOVA *et al*. 2021). All SNP positions used for this assay are listed in Table S33.

### Heterologous recombination assay

To examine the rate of heterologous recombination across different genetic backgrounds, we performed an assay using the *dpy25/mIn1[rol-1GFP] II C. elegans* strain (LEÓN-ORTIZ *et al*. 2018). To score heterologous recombination, a heterozygous parent carrying one normal copy of chromosome II marked with the semi-dominant *dpy-25* mutation and one copy with the *mln-1* inversion marked with a recessive *rol-e91* mutation and a semi-dominant GFP-expressing transgene was used to score recombinant progeny. In the absence of recombination within the *mln-1* inversion, the expected progeny would be 50% heterozygous *dpy-25/mIn1*, 25% homozygous *dpy-25/dpy-25*, and 25% homozygous *mIn1/mIn1*, all of which genotypes can be distinguished phenotypically. An illegitimate crossover between *mln-1* and the normal copy of chromosome II would produce different phenotypic combinations. We crossed different mutant strains into the *mln-1* strain and recorded illegitimate recombination events in the progeny.

### Intersister/intrachromatid repair (ICR) assay

ICR assays were performed as previously described (TORAASON *et al*. 2021). Parent hermaphrodites (P0) of each genotype were generated by crossing DLW14 hermaphrodites to wild-type or mutant males of interest to generate *unc-5(lib1)/+IV; krIs14/+V* males. F1 males were crossed with CB791 hermaphrodites to generate *unc-5(lib1)/unc-5(e791) IV; krIs14/+V* hermaphrodites.

P0 L4 hermaphrodites of the desired genotype were isolated 16–18 h before heat shock and maintained at 16°C. All worms were maintained on and regularly transferred to NGM plates seeded with OP50. Heat shock was performed at 34°C for 1 h, followed by recovery at 20°C for 9 h. P0 hermaphrodites were placed on individual small plates and transferred to fresh plates at 22, 34, and 46 h after heat shock. At 58 h after heat shock, P0 hermaphrodites were discarded. Plates with P0 hermaphrodites were maintained at 20°C, whereas plates containing F1 progeny were placed at 16°C. Plates containing F1 progeny were shifted to 25°C about 18 h prior to scoring. F1 progeny were scored for GFP fluorescence approximately 54–70 h after removal of the P0 hermaphrodite. F1 individuals exhibiting recombinant fluorescence patterns were isolated and lysed for subsequent sequencing.

### Materials and Methods (Supplemental Figures) Colocalization analysis

Colocalization analysis was performed using the Colocalization Finder plugin in ImageJ. Images were converted to 8-bit images prior to analysis. Intensity thresholds were set to a minimum of 40 for each channel, based on manual background signal estimation. The gonad was divided into seven zones along its length using a custom ImageJ script. Nuclei were manually selected in each zone, and a macro was used to extract Manders’ colocalization coefficients (M1 and M2) for each nucleus. Quantification was performed on maximum intensity projections. The Manders’ colocalization coefficient was used to quantify the proportion of signal overlap between channels above the defined thresholds.

### Focus detection and data processing

RPA-1 and RAD-51 foci were quantified using a custom ImageJ macro applied to manually curated nuclear ROIs defined with Cellpose. Within each nucleus, local intensity maxima were detected with the “Find Maxima” function using a fixed prominence threshold of 75. The total number of foci per nucleus was recorded. All measurements were exported as comma-separated values (CSV) files. Raw output files were processed with custom bash scripts to correct misaligned entries generated during export and to associate measurements with their corresponding nuclei. Cleaned datasets were parsed to extract foci per nucleus counts across all samples. Final datasets consisting of foci per nucleus counts were used for downstream statistical analysis. All processing steps were applied uniformly across samples.

### Hydroxyurea treatment

At 24 h after the L4 stage, worms were transferred to NGM plates containing 40 mM hydroxyurea and seeded with *E. coli* for 8 h. Worms were recovered and transferred to NGM plates seeded with *E. coli*. Each day, worms were transferred to fresh plates, and for 3 days the embryonic viability of the eggs laid was assessed.

### Irradiation

L4 hermaphrodites were picked and incubated at 20°C for 16–20 h. Following this, worms were irradiated at 60 Gy using a ^137^Cs source (Gammacell 3000). 10 worms per genotype were placed onto individual small NGM plates (one worm per plate) and scored in a 3-day viability assay.

## Acknowledgements

We thank N. Silva, D. Slade, and D. Libuda for sharing reagents and E. Ogris and members of the Jantsch laboratory, N. Silva and Petr Cejka for helpful discussions. We are grateful to D. Bradley for valuable input on identifying CDC–7 kinase domains. We thank J. Gotzmann for access to the Microscopy Facility and technical support. Some strains were provided by the Caenorhabditis Genetics Center, which is funded by the NIH Office of Research Infrastructure Programs (P40 OD010440). All LC-MS/MS mass spectrometry analyses were performed on instruments of the Vienna BioCenter Core Facilities (VBCF). We thank Elisabeth Roitinger and her team at the Proteomics Facility (GMI/IMBA/IMP) and the Mass Spectrometry Facility at Max Perutz Labs for their support. The study received funding from the Austrian Science Fund (FWF) grant PAT2512023, Grant-DOI 10.55776/PAT2512023 and the SFB project F 8805-B (VJ), Grant-DOI 10.55776/F88. ABK was funded by the Austrian Science Fund (FWF) doc.funds ‘Genome Instability’ Grant-DOI 10.55776/DOC195. F.M. was funded by the ESPRIT program project number ESP 566, Grant-DOI 10.55776/ESP566.

## Author contributions

Cell biology experiments: A.B., A.B.K., L.K., and L.H.

Bioinformatics analysis: A.S.

Biochemistry: A.B.

Mass spectrometry analysis: F.M., K.M. and M.H.

Worm strains: A.B., A.B.K., L.K., L.H., M.R.D.S., F.P., and A.G.

Project conception, data analysis: A.B., A.B.K, J.M. and V.J.

Writing the manuscript: A.B. and V.J.

## Data availability

All data needed to evaluate the conclusions in the paper are present in the paper and/or the Supplementary Materials. Workflows used to quantify the signal intensity and to count the foci (RPA-1/RAD-51) are archived on Zenodo under DOI 10.5281/zenodo.20138040 (BAUDRIMONT AND JANTSCH 2026).

## Supplemental Figures

**Figure S1.**
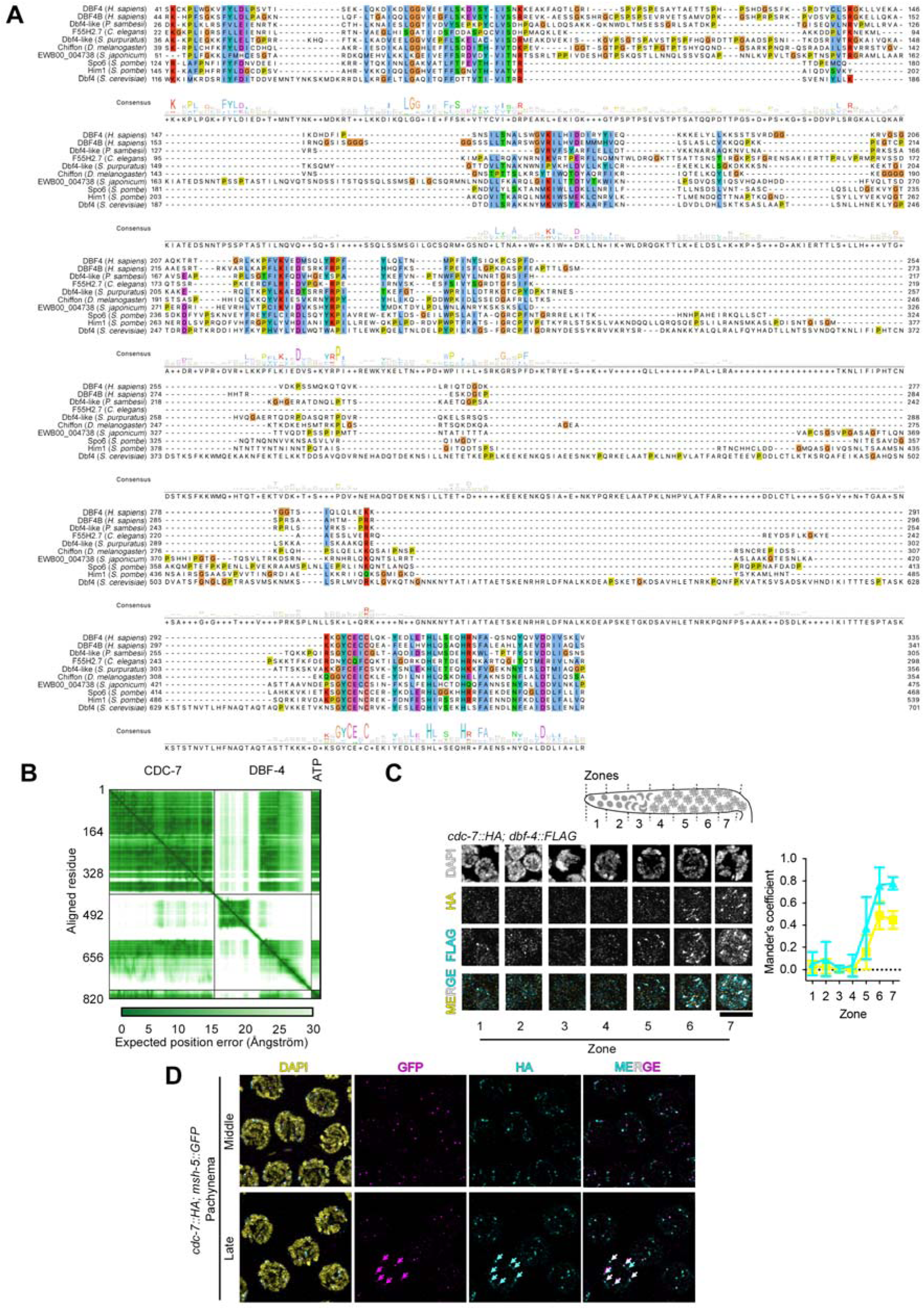
Colocalization of CDC–7 and DBF–4 in meiotic prophase and CDC–7 colocalization with MSH-5. **A.** Multiple sequence alignment of Dbf4 orthologs from vertebrates, invertebrates, and fungi. Conserved residues are highlighted according to amino acid properties. The predicted *Caenorhabditis elegans* Dbf4 ortholog F55H2.7 is aligned with Dbf4 proteins from *Homo sapiens*, *Pectus sambesii*, *Strongylocentrotus purpuratus*, *Drosophila melanogaster*, *Schistosoma japonicum*, *Schizosaccharomyces pombe*, and *Saccharomyces cerevisiae*. **B.** Predicted aligned error heatmap derived from AlphaFold modeling of the CDC–7–DBF–4 complex in the presence of ATP. Axes indicate residue positions across the full-length proteins. **C.** Top, schematic representation of the gonad divided into seven zones of equal length from the progenitor zone to late pachynema. Bottom left, immunodetection of CDC–7::HA (yellow) and DBF–4::FLAG (cyan) with DAPI counterstaining (white) across these zones (1–7). Bottom right, quantification of colocalization using Manders’ coefficient across zones. Scale bar, 5 µm. **D.** Immunodetection of CDC–7::HA (cyan) and MSH-5::GFP (magenta) with DAPI counterstaining (yellow) in middle and late pachynema. Cyan and magenta arrows indicate CDC–7::HA and MSH-5::GFP foci, respectively; white arrows in merged images indicate sites of colocalization between both markers. Scale bar, 10 µm.

**Figure S2.**
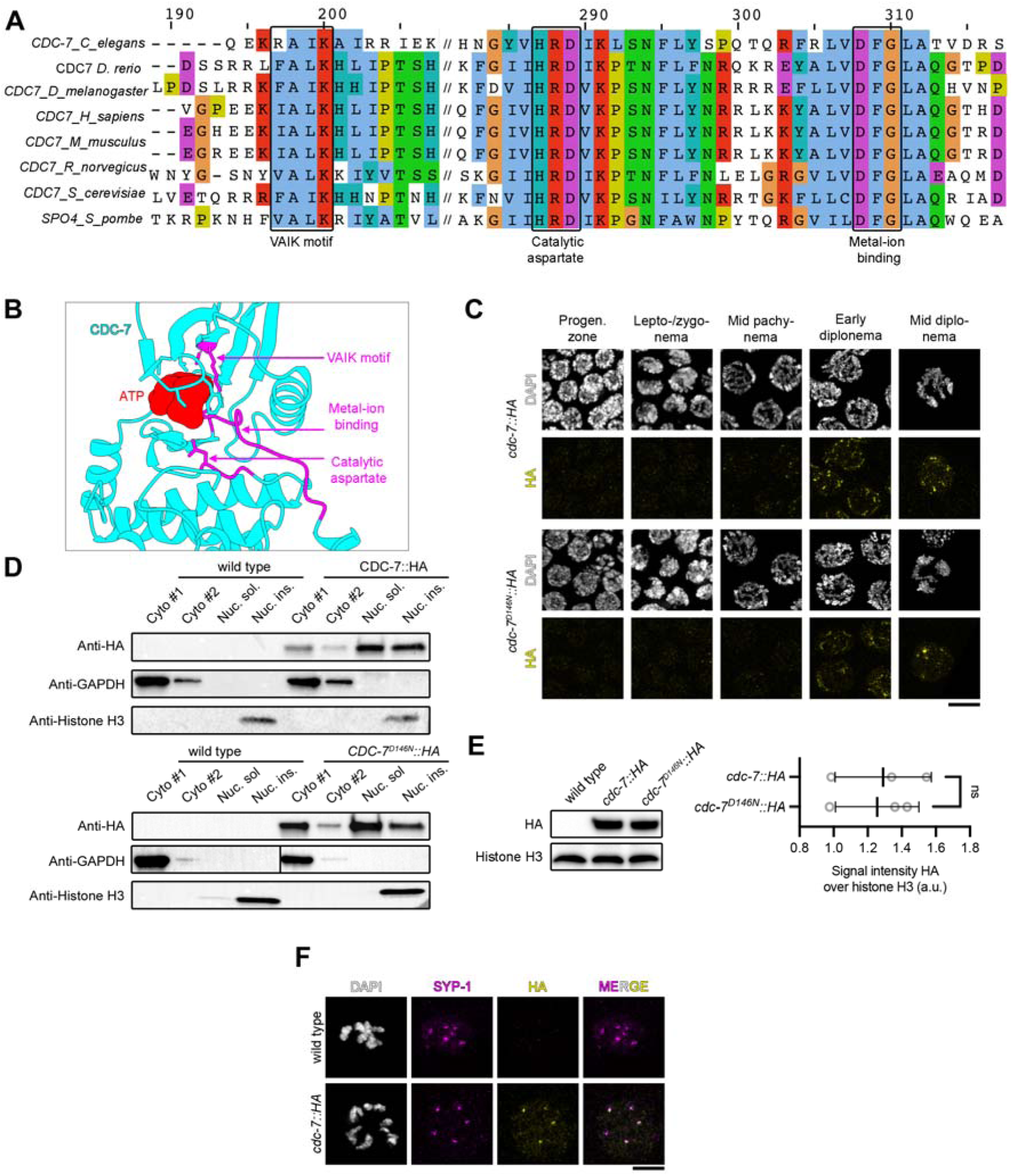
Conserved CDC–7 kinase features, catalytic activity-independent loading, and localization to chromosome short arms. **A.** Multiple sequence alignment of CDC7 orthologs from *Caenorhabditis elegans*, *Danio rerio*, *Drosophila melanogaster*, *Homo sapiens*, *Mus musculus*, *Rattus norvegicus*, *Saccharomyces cerevisiae*, and *Schizosaccharomyces pombe*. Conserved residues are highlighted, including the VAIK motif, catalytic aspartate, and metal-ion binding residues. **B.** Structural model of CDC–7, highlighting key functional elements. The ATP-binding pocket is shown in red, and the VAIK motif, catalytic aspartate, and metal-ion-binding region in pink. **C.** Immunodetection of CDC–7::HA (top) or CDC–7^D146N^::HA (bottom) (yellow) with DAPI counterstaining (white) across meiotic prophase stages. Scale bar, 5 µm. **D.** Subcellular fractionation followed by immunoblotting of CDC–7::HA and CDC–7^D146N^::HA germline nuclei: cytoplasmic (Cyto #1 and #2) and nuclear soluble (Nuc. sol.) and insoluble (Nuc. ins.) fractions. GAPDH and Histone H3 are cytoplasmic and nuclear markers, respectively. **E.** Whole-worm extracts from *cdc*–*7::HA* and *cdc*–*7D146N::HA* animals analyzed by immunoblotting against HA and Histone H3. Histone H3 was used for normalization. Left, representative immunoblot. Right, quantification of HA signal intensity normalized to Histone H3 from three independent experiments. Statistical significance was determined using the Mann–Whitney test. **F.** Immunodetection of CDC–7::HA (yellow) and SYP-1 (magenta) with DAPI counterstaining (white) at the diakinesis stage in wild-type and *cdc*–*7::HA* animals. Scale bar, 5 µm.

**Figure S3.**
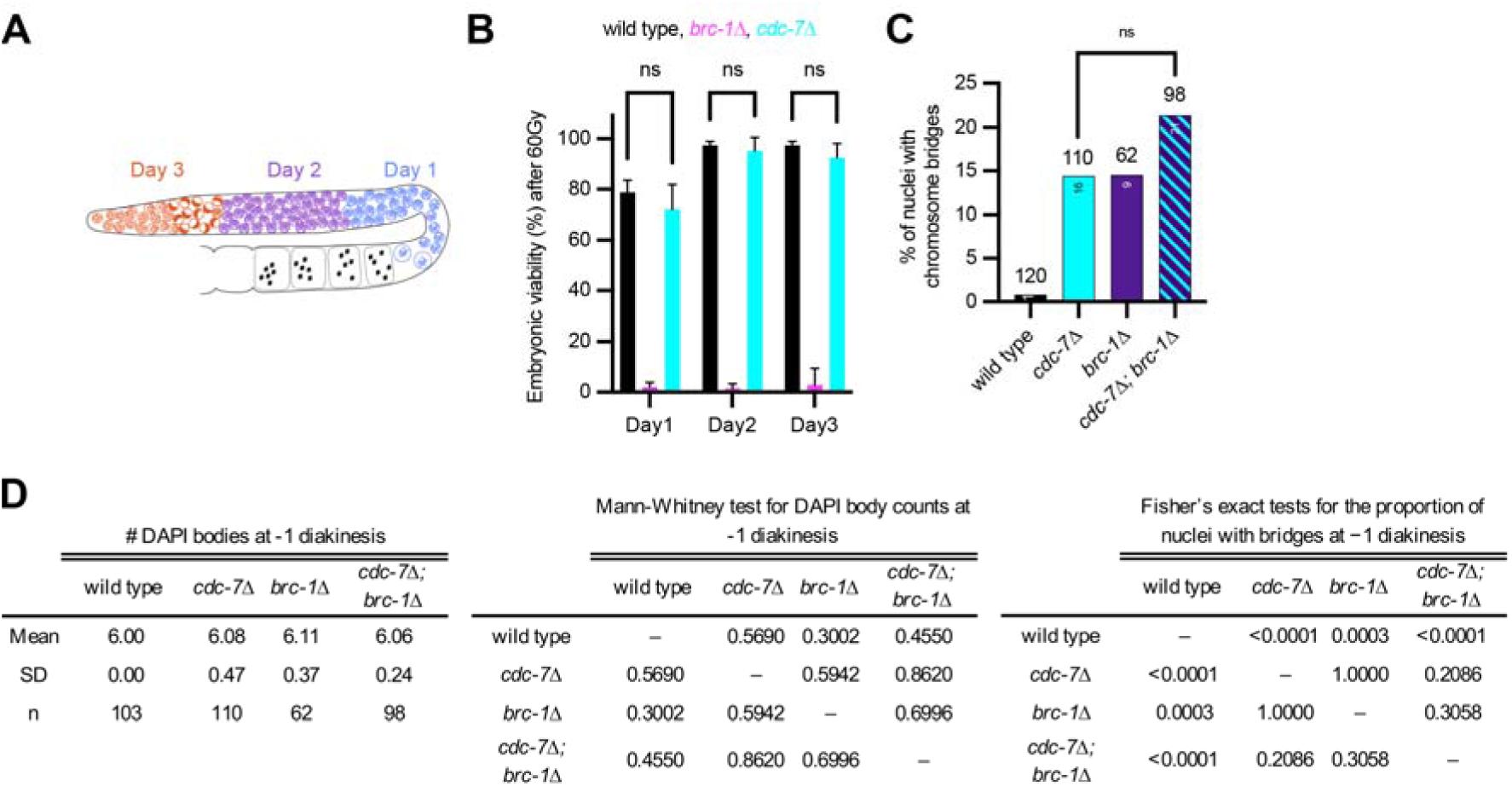
CDC–7 is irradiation insensitive, unlike BRC–1, and the double mutant exhibits chromosome bridges. **A.** Schematic diagram indicating the position of nuclei within the germline at egg laying (Days 1–3) following 60 Gy irradiation. **B.** Embryonic viability following 60 Gy irradiation for the indicated genotypes across the indicated time points. Data are the mean ± SD Statistical significance was determined using the Mann–Whitney U test (ns, not significant). **C.** Percentage of nuclei with chromosome bridges at the −1 diakinesis for the indicated genotypes. Numbers of nuclei counted are indicated above the bars and numbers of bridges observed inside the bars. Statistical significance was determined using the Mann–Whitney U test (ns, not significant). Data for the wild type and *cdc*–*7*Δ are as shown in Fig. 3B. **D.** Quantification of DAPI bodies at the −1 diakinesis stage (left) and corresponding pairwise statistical analyses for DAPI body counts (center) and the proportion of nuclei with bridges (right) for the indicated genotypes. *P* values for DAPI body counts were calculated using the Mann–Whitney U test. *P* values for bridge frequency were calculated using Fisher’s exact test.

**Figure S4.**
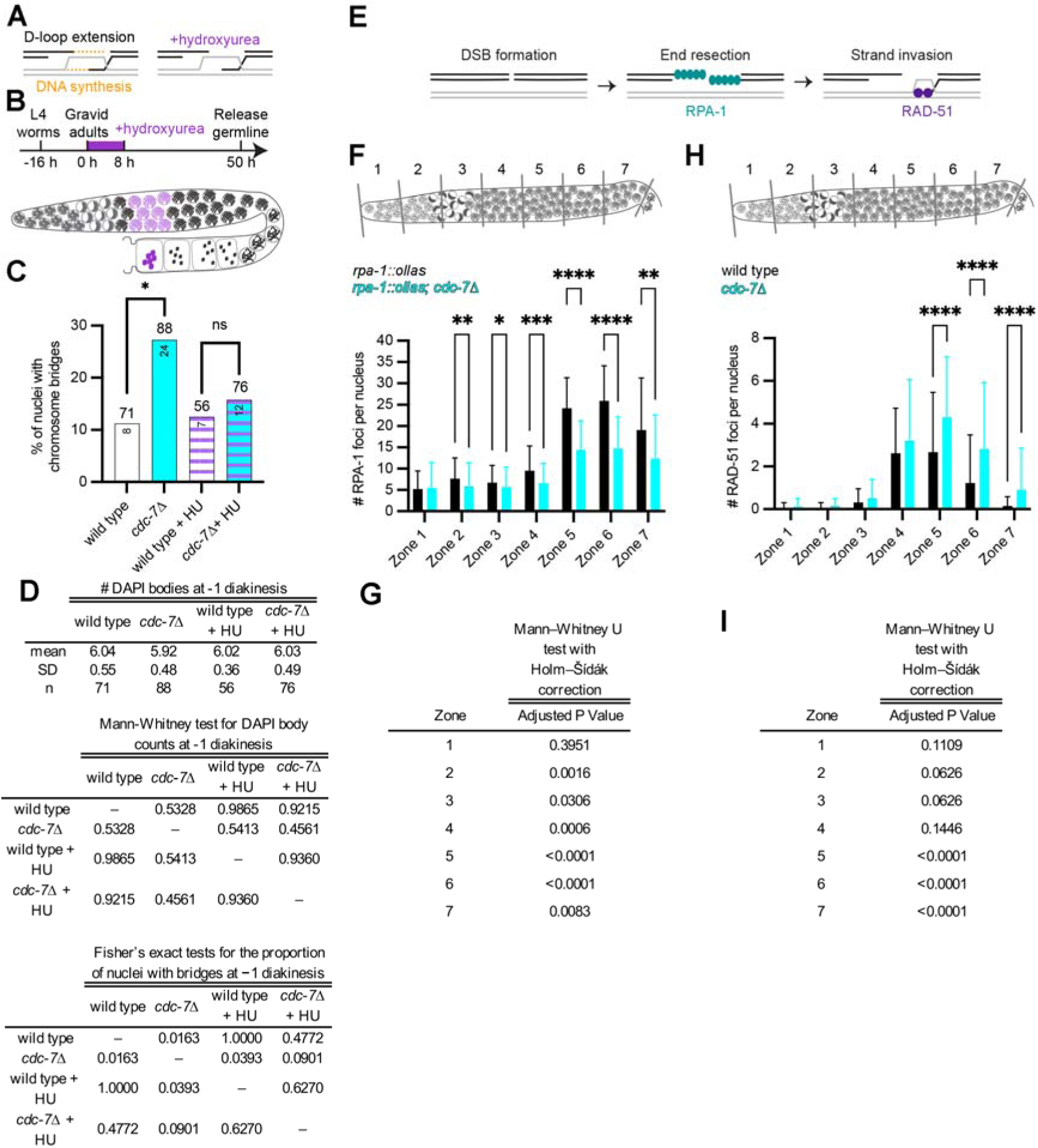
Bridge formation requires DNA synthesis, and the *cdc*–*7*Δ mutant has fewer RPA-1 foci and increased RAD-51 foci. **A.** Schematic diagram of D-loop extension and of DNA synthesis and its inhibition by hydroxyurea. **B.** Experimental design: gravid worms (16 h post L4) were treated with hydroxyurea for 8 h, and germlines were released 42 h later for analysis of diakinesis chromosomes (dark violet) in nuclei that had previously been located in early pachynema (light violet). **C.** Percentage of nuclei with chromosome bridges at the −1 diakinesis under the indicated conditions. Numbers of nuclei counted are indicated above the bars and numbers of bridges observed inside the bars. Baseline levels are elevated compared with other figures because older animals (3-day-old adults) were used. Statistical significance was determined using the Mann–Whitney U test (\**P* < 0.05). **D.** Quantification of DAPI bodies at the −1 diakinesis stage (top), corresponding pairwise statistical analyses for DAPI body counts (middle), and the proportion of nuclei with bridges (bottom) for the given genotypes under the indicated conditions. *P* values for DAPI body counts were calculated using the Mann–Whitney U test. *P* values for bridge frequency were calculated using Fisher’s exact test. **E.** Simplified schematic diagrams of recombination steps, including double-strand break (DSB) formation, end resection marked by RPA-1, and strand invasion marked by RAD-51. **F.** Quantification of RPA-1 foci per nucleus across germline zones 1–7, defined as equal-length segments from the progenitor zone to late pachynema (see schematic diagram), in *rpa-1::ollas* and *rpa-1::ollas; cdc*–*7*Δ germlines. Data are the mean ± SD. Statistical significance was determined using the Mann–Whitney U test (\**P* < 0.05, \*\**P* < 0.01, \*\*\**P* < 0.001, \*\*\*\**P* < 0.0001). **G.** Adjusted *P* values for the comparisons shown in **F**, as calculated using the Mann–Whitney U test with Holm–Šídák correction for multiple comparisons. **F.** Quantification of RAD-51 foci per nucleus across germline zones 1–7, defined as equal-length segments from the progenitor zone to late pachynema (see schematic diagram), in wild-type and *cdc*–*7*Δ germlines. Data are the mean ± SD. Statistical significance was determined using the Mann–Whitney U test (\*\*\*\**P* < 0.0001). **I.** Adjusted *P* values for the comparisons shown in **H**, as calculated using the Mann–Whitney U test withHolm–Šídák correction for multiple comparisons.

**Figure S5.**
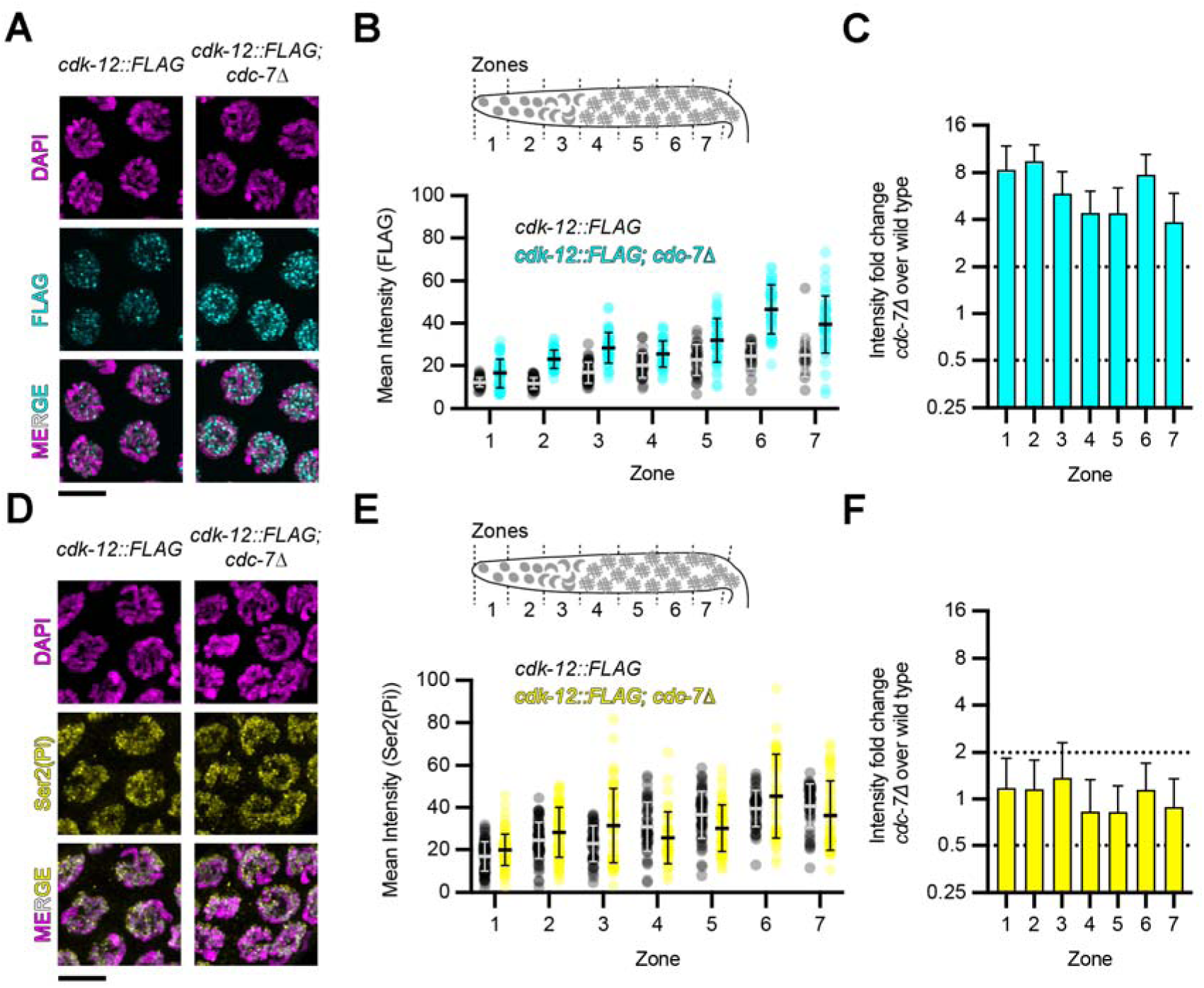
CDK–12 accumulation in *cdc*–*7*Δ is uncoupled from Ser2 phosphorylation levels. **A.** Immunodetection of CDK–12::FLAG (cyan) with DAPI counterstaining (magenta) in *cdk*–*12::FLAG* and *cdk*–*12::FLAG; cdc*–*7*Δ germline. Scale bar, 5 µm. **B.** Quantification of the mean intensity of the CDK–12::FLAG signal across germline zones 1–7, defined as equal-length segments from the progenitor zone to late pachynema (see schematic diagram), in *cdk*–*12::FLAG* and *cdk*–*12::FLAG; cdc*–*7*Δ germlines. Each point represents a nucleus; lines indicate the mean ± SD. **C.** Fold change in CDK–12::FLAG signal intensity in *cdc*–*7*Δ relative to the wild type across germline zones 1–7. **D.** Immunodetection of Ser2 phosphorylated RNA polymerase II (Ser2Pi; magenta) with CDK–12::FLAG (yellow) and DAPI counterstaining (white) in *cdk*–*12::FLAG* and *cdk*–*12::FLAG; cdc*–*7*Δ germlines. Scale bar, 5 µm. **E.** Mean intensity of the Ser2P signal across germline zones 1–7, defined as equal-length segments from the progenitor zone to late pachynema (see schematic diagram), in *cdk*–*12::FLAG* and *cdk*–*12::FLAG; cdc*–*7*Δ germlines. Each point represents a nucleus; lines indicate the mean ± SD. **F.** Fold change in Ser2P signal intensity in *cdc*–*7(knu709)* relative to the wild type across germline zones 1–7.

**Figure S6.**
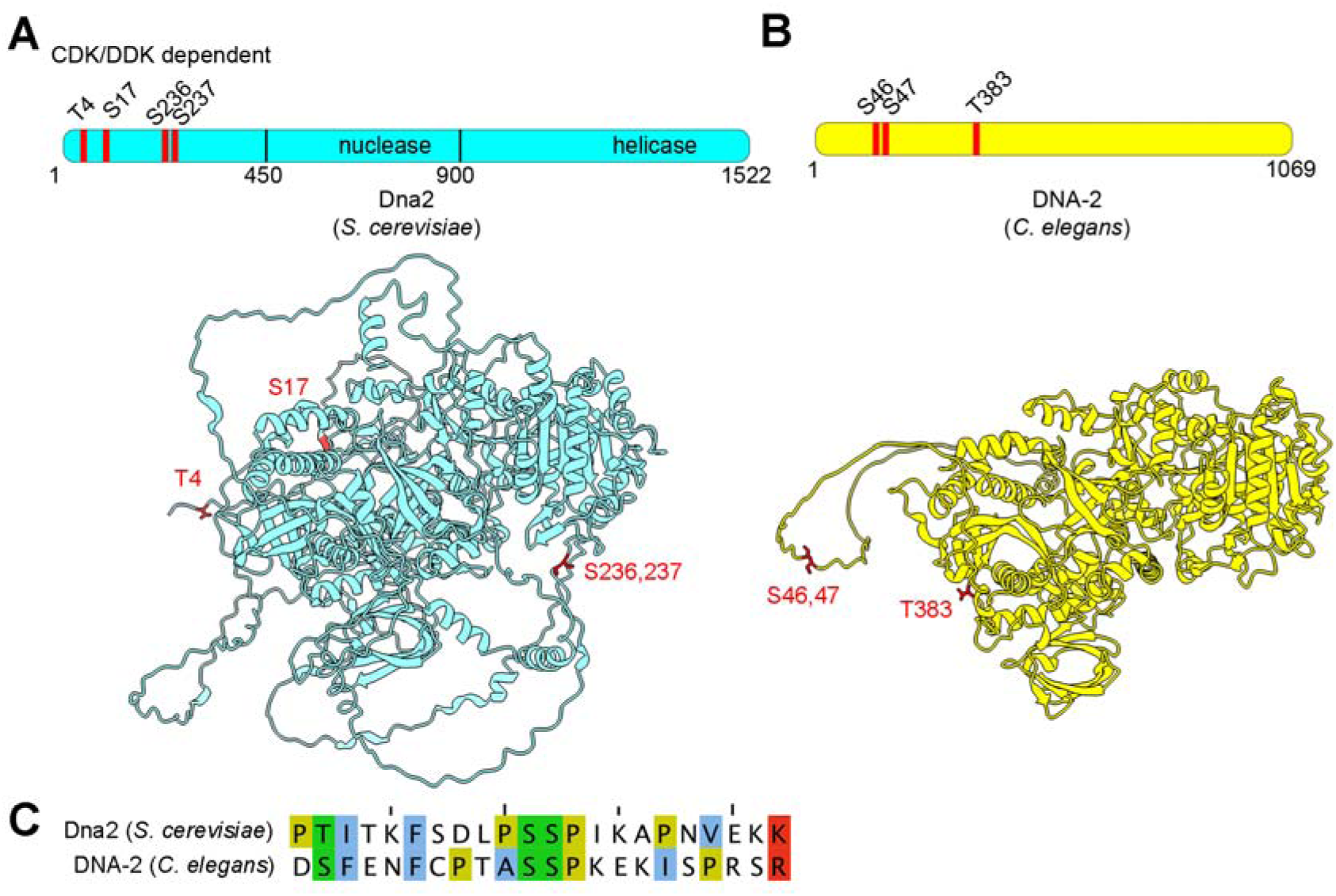
Identification and conservation of DNA–2 phosphorylation sites. **A.** Domain organization of Dna2 from *S. cerevisiae*, indicating conserved nuclease and helicase domains and reported CDK/DDK-dependent phosphorylation sites (T4, S17, S236, S237). The structural model predicted using AlphaFold highlights the position of these residues. **B.** Domain organization of DNA–2 from *C. elegans* showing candidate phosphorylation sites (S46, S47, and T383). The structural model predicted using AlphaFold highlights the position of these residues. **C.** Sequence alignment of Dna2 (*S. cerevisiae*) and DNA–2 (*C. elegans*) surrounding the conserved phosphorylation motifs.

## Supplementary Table

**Table S1.** Embryo hatch rate counts.

| Genotype | Viability (%) |
| --- | --- |
| Wild type | 99.9 ± 0.1 |
| *cdc 27(jf218[cdc 27::HA])* | 99.5 ± 0.6 |
| *cdc 27(syb6737[cdc 27::GFP])* | 99.1 ± 1.1 |
| *cdc 27(syb6737cdc 27::GFP); TurboID* | 99.0 ± 1.2 |
| *cdc 27(jf218[cdc 27::HA]); dbf 24(jf247[dbf 24::FLAG])* | 99.4 ± 1.1 |

**Table S2.** Spreadsheet of all proteins identified in triplicate CDC–7::HA immunoprecipitation experiments followed by mass spectrometry analysis.

**Table S3.** Spreadsheet of all proteins identified in triplicate CDC–7::GFP proximity ligation assays followed by mass spectrometry analysis.

**Table S4.** Spreadsheet of all phosphosites identified in triplicate measurements after Fe-NTA-based phosphopeptide enrichment followed by mass spectrometry analysis.

**Table S5.** Number of DAPI bodies at the −1 diakinesis for the indicated genotypes.

| Genotype | Average | SD | n |
| --- | --- | --- | --- |
| Wild type | 6.00 | 0.00 | 103 |
| *cdc 7(knu709[cdc 7Δ])* | 6.08 | 0.47 | 110 |
| *dbf 4(jf255[dbf 4<sup>STOP</sup>])* | 5.75 | 0.67 | 64 |
| *cdc 7(syb9555[cdc 7<sup>D146N</sup>])* | 5.59 | 0.81 | 49 |
| *spo-11(ok79[spo-11Δ])* | 11.94 | 0.24 | 51 |
| *cdc 7(knu709[cdc 7Δ]);*<br>*spo-11(ok79[spo-11Δ])* | 12.00 | 0.00 | 52 |
| *msh-5(me23[msh-5Δ])* | 11.72 | 0.53 | 54 |
| *cdc 7(knu709[cdc 7Δ]);*<br>*msh-5(me23[msh-5Δ])* | 11.40 | 0.92 | 47 |
| *exo-1(tm1842[exo-1Δ])* | 5.77 | 0.57 | 56 |
| *cdc 7(knu709[cdc 7Δ]);*<br>*exo-1(tm1842[exo-1Δ])* | 5.27 | 0.85 | 59 |
| *cdc 7(knu709[cdc 7Δ])*<br>*48 h EtOH* | 5.66 | 0.70 | 55 |
| *dna 2(iow11[dna 2<sup>AID</sup>])*<br>*48 h EtOH* | 5.54 | 0.73 | 50 |
| *dna 2(iow11[dna 2<sup>AID</sup>])*<br>*48 h auxin* | 5.79 | 0.71 | 61 |
| *cdc 7(knu709[cdc 7Δ]);*<br>*dna 2(iow11[dna 2<sup>AID</sup>])*<br>*48 h EtOH* | 5.52 | 0.74 | 50 |
| *cdc 7(knu709[cdc 7Δ]);*<br>*dna 2(iow11[dna 2<sup>AID</sup>])*<br>*48 h auxin* | 5.66 | 0.58 | 59 |
| *dna 2(iow11[dna 2<sup>AID</sup>]);*<br>*exo-1(tm1842[exo-1Δ])*<br>*48 h EtOH* | 6.04 | 0.72 | 47 |
| *dna22(iow11[dna22<sup>AID</sup>]);*<br>*exo-1(tm1842[exo-1Δ])*<br>48 h auxin | 5.71 | 0.81 | 51 |
| *cdc7(knu709[cdc7Δ]);*<br>*dna22(iow11[dna22<sup>AID</sup>]);*<br>*exo-1(tm1842[exo-1Δ])*<br>48 h EtOH | 5.60 | 0.76 | 50 |
| *cdc7(knu709[cdc7Δ]);*<br>*dna22(iow11[dna22<sup>AID</sup>]);*<br>*exo-1(tm1842[exo-1Δ])*<br>48 h auxin | 5.50 | 0.77 | 60 |
| *syp-2(ok307[syp-2Δ])* | 11.87 | 0.34 | 60 |
| *syp-2(ok307[syp-2Δ]);*<br>*cdc7(knu709[cdc7Δ])* | 10.55 | 1.12 | 55 |
| *msh-6(pk2504[msh-6Δ])* | 5.85 | 0.53 | 41 |
| *msh-6(pk2504[msh-6Δ]);*<br>*cdc7(knu709[cdc7Δ])* | 5.90 | 0.92 | 57 |
| *lig-4(ok716[lig-4Δ])* | 5.81 | 0.48 | 58 |
| *lig-4(ok716[lig-4Δ]);*<br>*cdc7(knu709[cdc7Δ])* | 5.81 | 0.61 | 94 |
| *cdk12(syb9527[cdk12<sup>S30A</sup>])* | 5.68 | 0.57 | 60 |
| *cdk12(syb9527[cdk12<sup>S30A</sup>]);*<br>*cdc7(knu709[cdc7Δ])* | 5.82 | 0.56 | 76 |
| *cdk12(syb9536[cdk12<sup>S30D</sup>])* | 5.96 | 0.21 | 45 |
| *cdk12(syb9536[cdk12<sup>S30D</sup>]);*<br>*cdc7(knu709[cdc7Δ])* | 5.98 | 0.38 | 64 |
| *dna22(jf285[dna22<sup>S46A, S47A</sup>])* | 5.58 | 0.73 | 71 |
| *cdc7(knu709[cdc7Δ]);*<br>*dna22(jf285[dna22<sup>S46A, S47A</sup>])* | 5.90 | 0.47 | 67 |
| *cdk12(syb9527[cdk12<sup>S30A</sup>]);*<br>*dna22(jf285dna22<sup>S46A, S47A</sup>])* | 5.70 | 0.62 | 71 |

| Genotype | Average | SD | *n* |
| --- | --- | --- | --- |
| *dna* 72(jf286[*dna* 72<sup>S46D, S47D</sup>]) | 5.81 | 0.47 | 62 |
| *cdc* 77(knu709[*cdc* 77Δ]);<br>*dna* 72(jf286[*dna* 72<sup>S46D, S47D</sup>]) | 5.88 | 0.38 | 64 |
| *cdk* 12(syb9527[*cdk* 12<sup>S30A</sup>]);<br>*dna* 72(jf286[*dna* 72<sup>S46D, S47D</sup>]) | 5.88 | 0.32 | 68 |

**Table S6.** *P* values for the Mann–Whitney U test for DAPI body counts at −1 diakinesis for the indicated genotypes.

|  | Wild type | *cdc 27*Δ | *cdc 27*<sup>D146N</sup> | *dbf 24*<sup>STOP</sup> |
| --- | --- | --- | --- | --- |
| Wild type | – | 0.5690 | 0.0626 | 0.0043 |
| *cdc 27*Δ | 0.5690 | – | 0.0223 | 0.0014 |
| *cdc 27*<sup>D146N</sup> | 0.0626 | 0.0223 | – | 0.3023 |
| *dbf 24*<sup>STOP</sup> | 0.0043 | 0.0014 | 0.3023 | – |

**Table S7.**
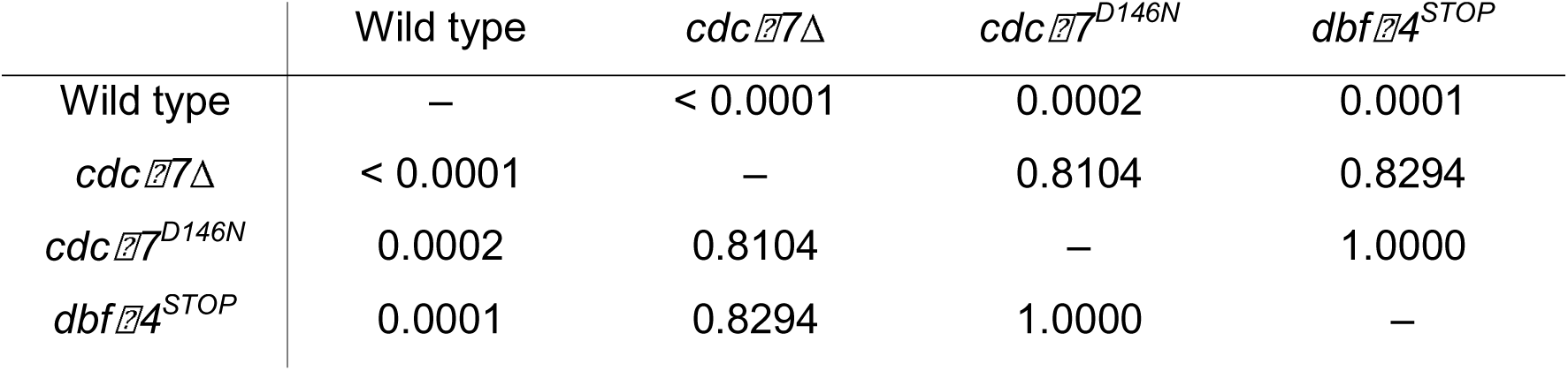
*P* values for pairwise Fisher’s exact tests for the proportion of nuclei with bridges at −1 diakinesis for the indicated genotypes.

| | Wild type | *cdc 27*Δ | *cdc 27*<sup>D146N</sup> | *dbf 24*<sup>STOP</sup> |
| --- | --- | --- | --- | --- |
| Wild type | — | < 0.0001 | 0.0002 | 0.0001 |
| *cdc 27*Δ | < 0.0001 | — | 0.8104 | 0.8294 |
| *cdc 27*<sup>D146N</sup> | 0.0002 | 0.8104 | — | 1.0000 |
| *dbf 24*<sup>STOP</sup> | 0.0001 | 0.8294 | 1.0000 | — |

**Table S8.**
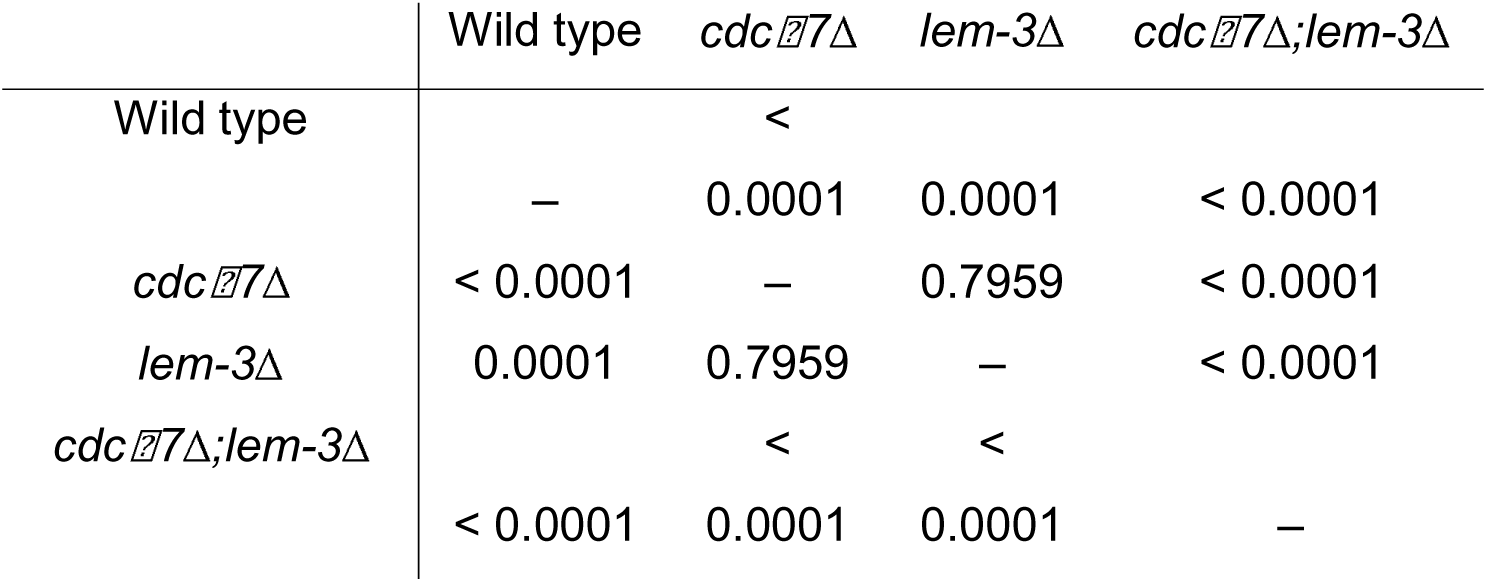
*P* values for the Mann–Whitney U test for embryonic lethality for the indicated genotypes.

**Table S9.** *P* values for the Mann–Whitney U test for DAPI body counts at −1 diakinesis for the indicated genotypes.

|  | Wild<br>type | *cdc 77Δ* | *spo-11Δ* | *cdc 77Δ;*<br>*spo-11Δ* | *msh-5Δ* | *cdc 77Δ;*<br>*msh-5Δ* |
| --- | --- | --- | --- | --- | --- | --- |
| Wild<br>type | – | 0.5690 | <<br>0.0001 | <<br>0.0001 | <<br>0.0001 | <<br>0.0001 |
| *cdc 77Δ* | 0.5690 | – | <<br>0.0001 | <<br>0.0001 | <<br>0.0001 | <<br>0.0001 |
| *spo-11Δ* | <<br>0.0001 | <<br>0.0001 | – | 0.6107 | 0.1057 | 0.0012 |
| *cdc 77Δ;*<br>*spo-11Δ* | <<br>0.0001 | <<br>0.0001 | 0.6107 | – | 0.0325 | 0.0002 |
| *msh-5Δ* | <<br>0.0001 | <<br>0.0001 | 0.1057 | 0.0325 | – | 0.0695 |
| *cdc 77Δ;*<br>*msh-5Δ* | <<br>0.0001 | <<br>0.0001 | 0.0012 | 0.0002 | 0.0695 | – |

**Table S10.**
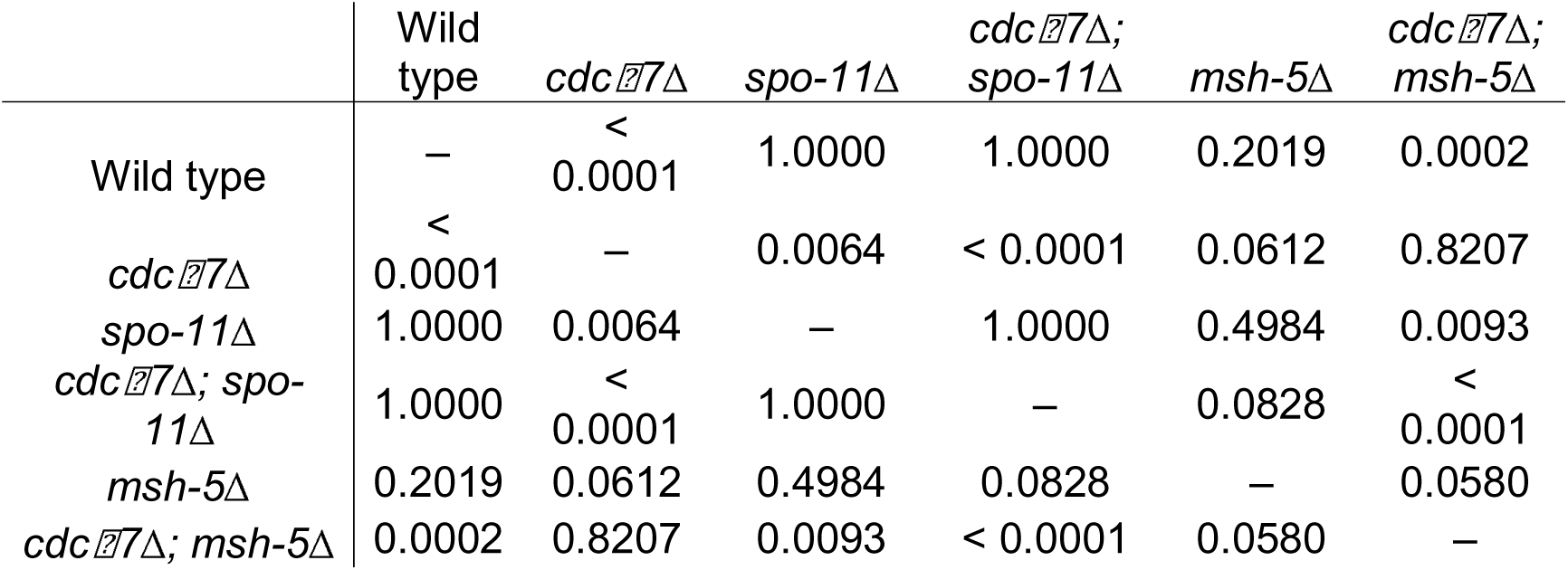
*P* values for pairwise Fisher’s exact tests for the proportion of nuclei with bridges at −1 diakinesis for the indicated genotypes.

**Table S11.** Number and percentage (in parenthesis) of non-crossovers (NCO), single crossovers (SCO), and double crossovers (DCO) in the indicated genotypes.

| Chromosome<br>V | NCO | SCO | DCO |
| --- | --- | --- | --- |
| Wild type | 150 (60.2%) | 99 (39.8%) | 0 (0%) |
| *cdc 27*Δ | 157 (59.5%) | 100 (37.9%) | 7 (2.7%) |

**Table S12.** *P* values for the Mann–Whitney U test for DAPI body counts at −1 diakinesis for the indicated genotypes.

|  | Wild<br>type | *cdc 27*Δ | *exo-1*Δ | *cdc 27*Δ;<br>*exo-1*Δ |
| --- | --- | --- | --- | --- |
| Wild<br>type | – | 0.5690 | 0.0412 | <<br>0.0001 |
| *cdc 27*Δ | 0.5690 | – | 0.0151 | <<br>0.0001 |
| *exo-1*Δ | 0.0412 | 0.0151 | – | 0.0015 |
| *cdc 27*Δ;<br>*exo-1*Δ | <<br>0.0001 | <<br>0.0001 | 0.0015 | – |

**Table S13.**
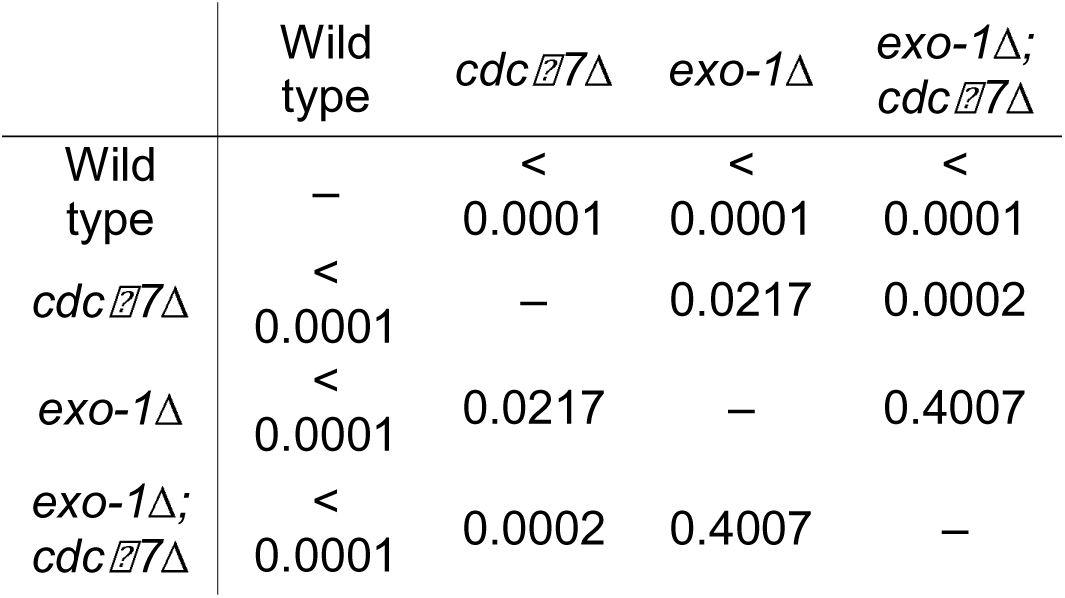
*P* values for pairwise Fisher’s exact tests for the proportion of nuclei with bridges at −1 diakinesis for the indicated genotypes.

|  | Wild<br>type | *cdc 7*Δ | *exo-1*Δ | *exo-1*Δ;<br>*cdc 7*Δ |
| --- | --- | --- | --- | --- |
| Wild<br>type | – | <<br>0.0001 | <<br>0.0001 | <<br>0.0001 |
| *cdc 7*Δ | <<br>0.0001 | – | 0.0217 | 0.0002 |
| *exo-1*Δ | <<br>0.0001 | 0.0217 | – | 0.4007 |
| *exo-1*Δ;<br>*cdc 7*Δ | <<br>0.0001 | 0.0002 | 0.4007 | – |

**Table S14.**
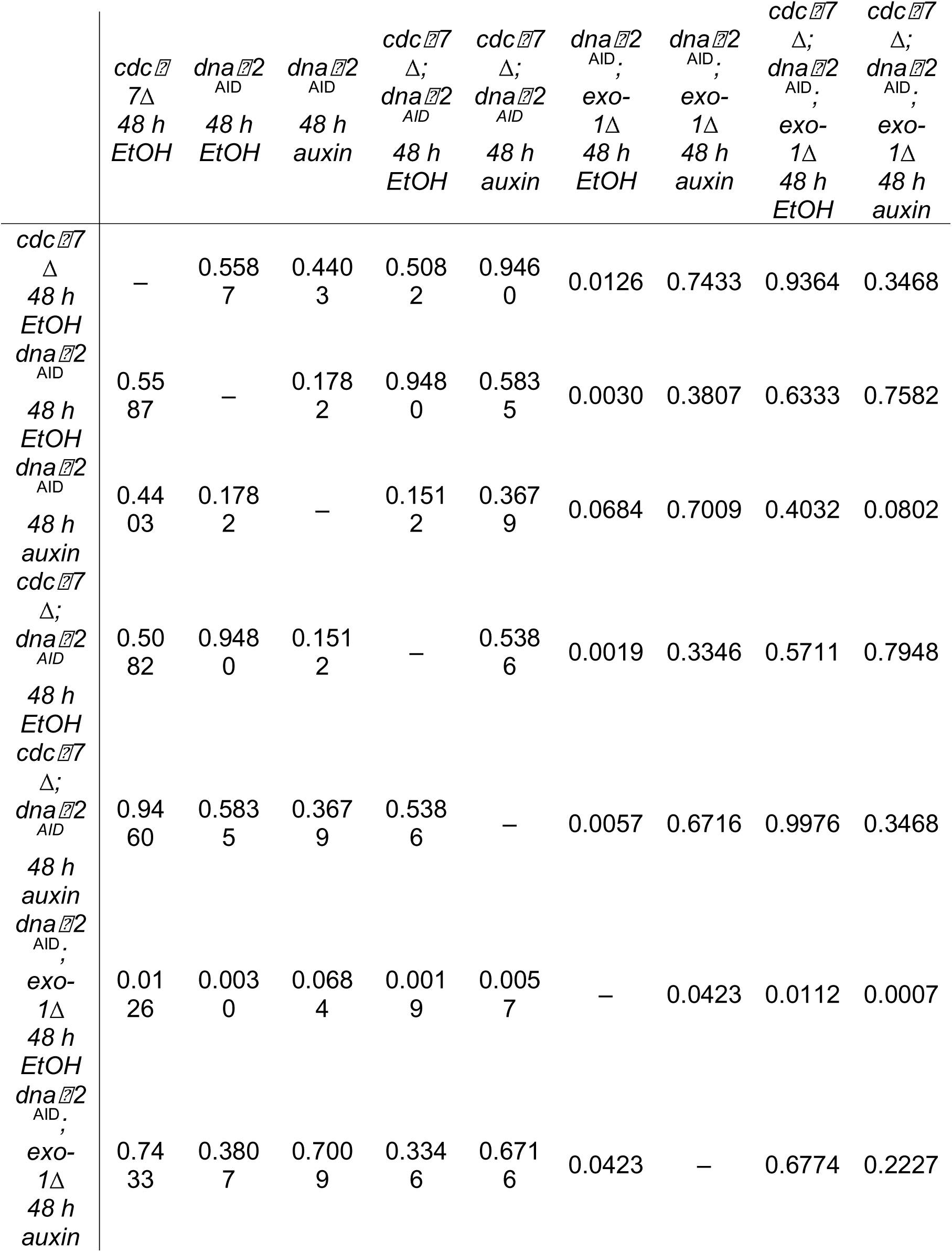
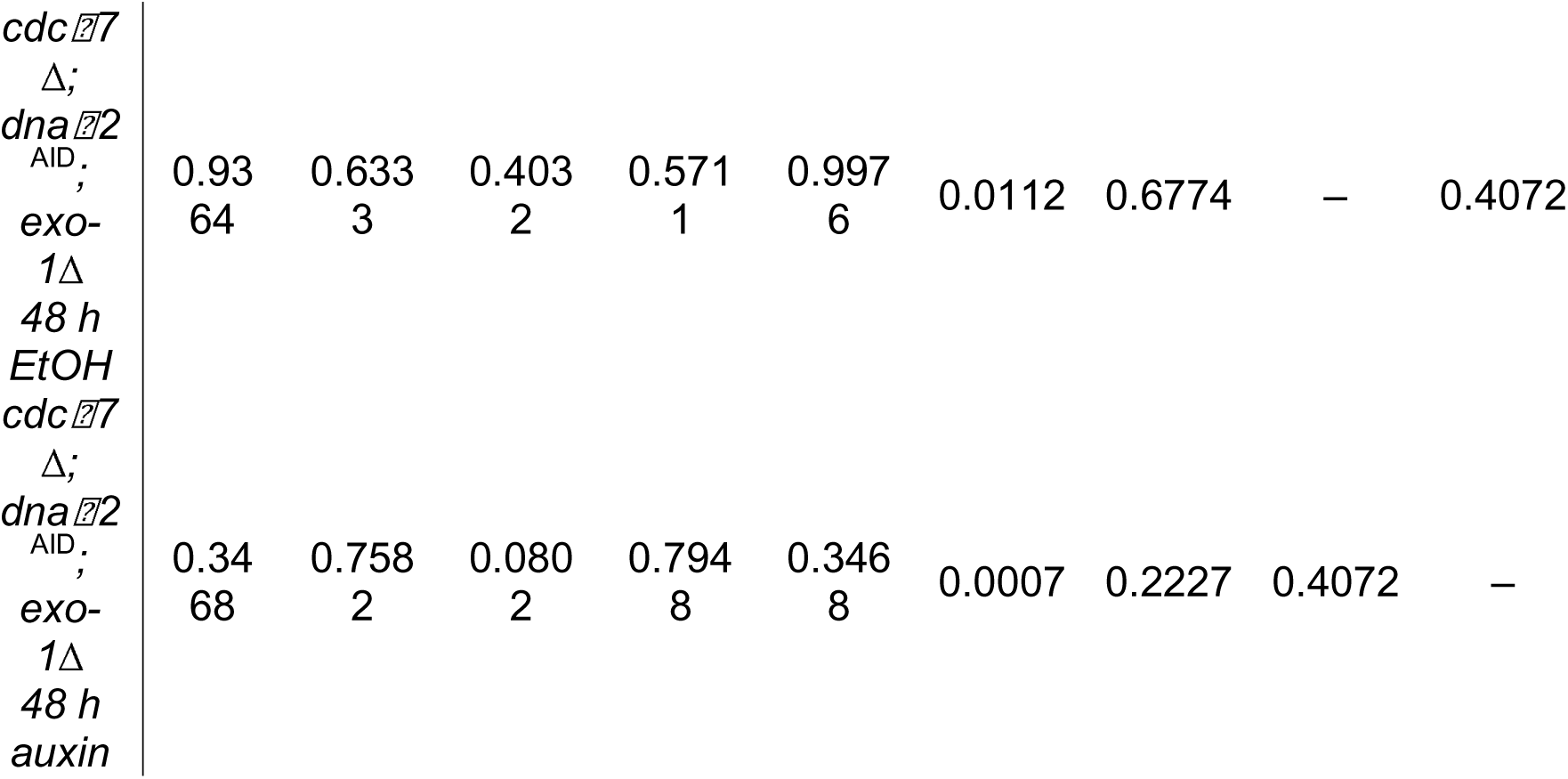
*P* values for the Mann–Whitney U test for DAPI body counts at −1 diakinesis for the indicated genotypes.

**Table S15.**
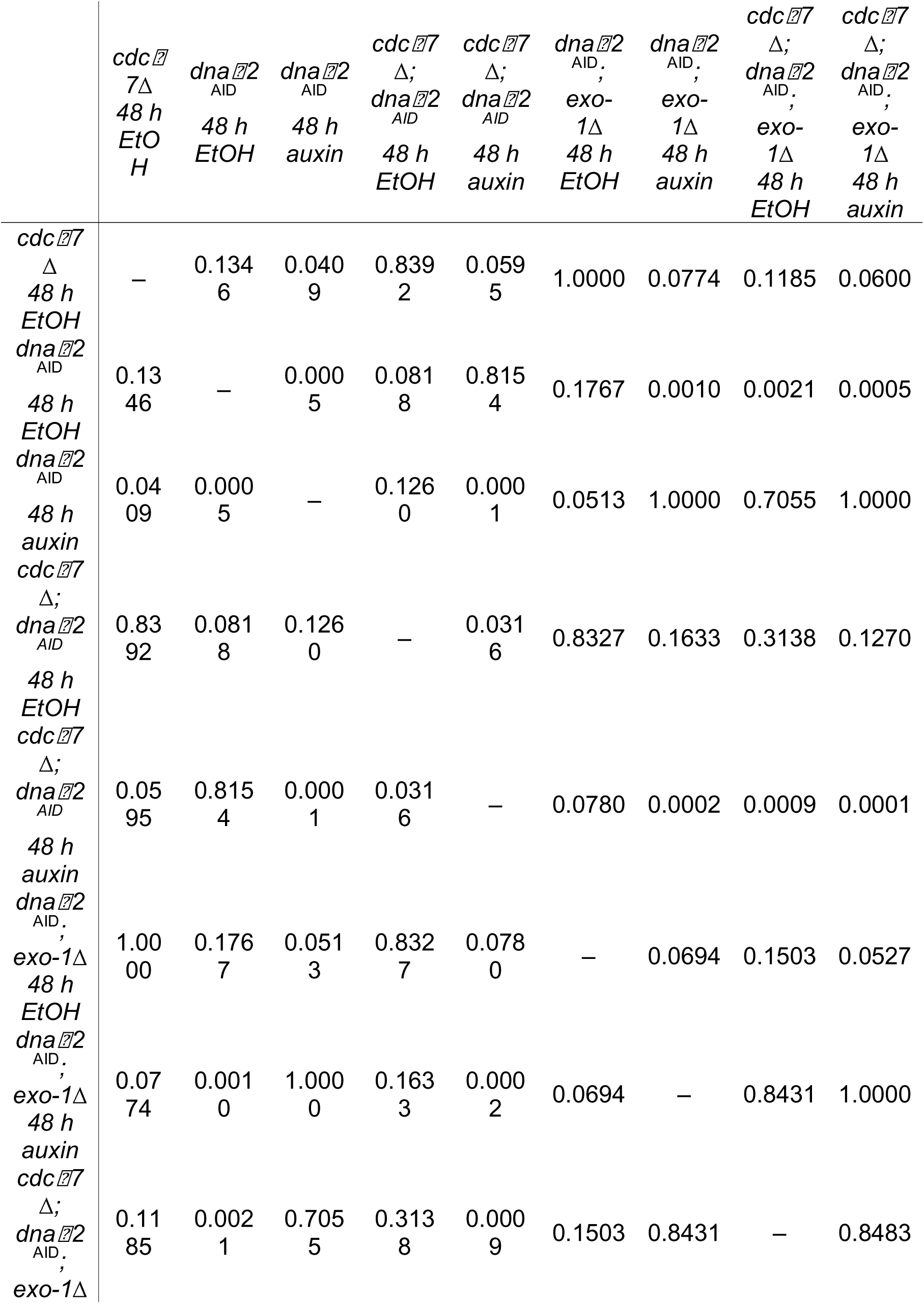
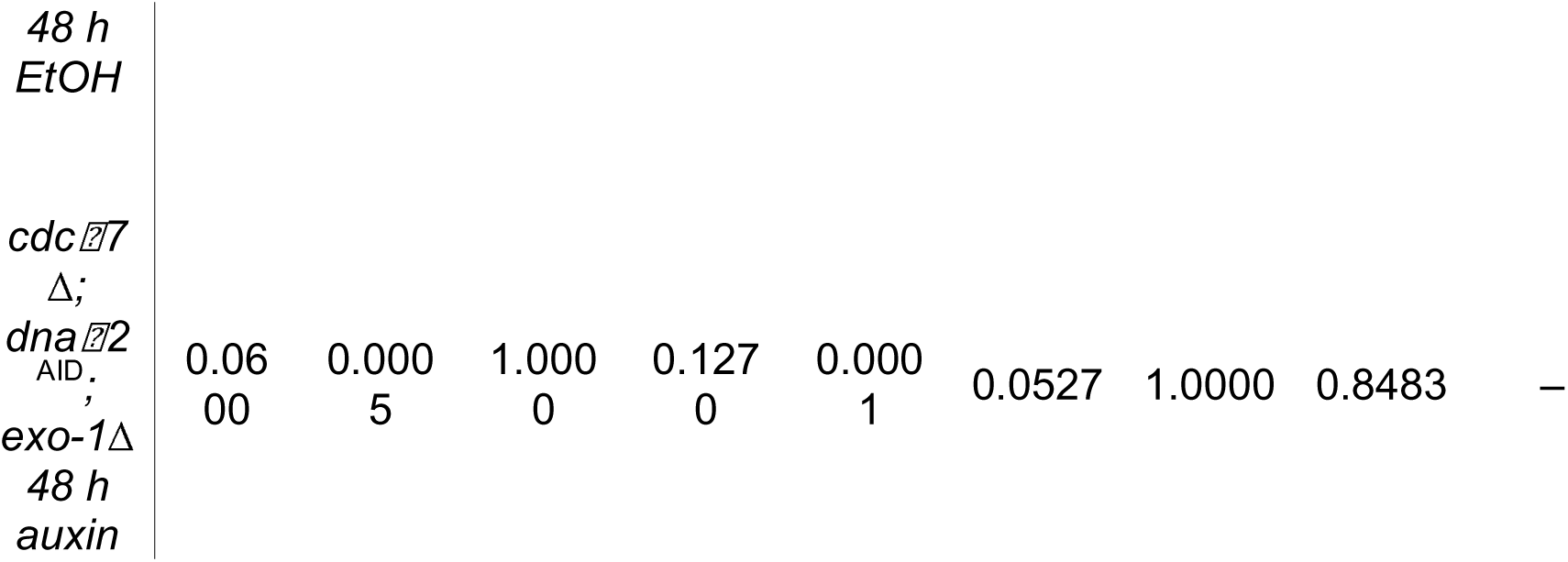
*P* values for pairwise Fisher’s exact tests for the proportion of nuclei with bridges at −1 diakinesis for the indicated genotypes.

**Table S16.** *P* values for the Mann–Whitney U test for DAPI body counts at −1 diakinesis for the indicated genotypes.

|  | Wild type | *cdc 77Δ* | *syp-2Δ* | *syp-2Δ;*<br>*cdc 77Δ* | *msh-6Δ* | *msh-6Δ;*<br>*cdc 77Δ* | *lig-4Δ* | *lig-4Δ;*<br>*cdc 77Δ* |
| --- | --- | --- | --- | --- | --- | --- | --- | --- |
| Wild type | – | 0.5690 | < 0.0001 | < 0.0001 | 0.1734 | 0.2734 | 0.1035 | 0.1574 |
| *cdc 77Δ* | 0.5690 | – | < 0.0001 | < 0.0001 | 0.0867 | 0.1312 | 0.0404 | 0.0552 |
| *syp-2Δ* | < 0.0001 | <<br>0.0001 | – | < 0.0001 | <<br>0.0001 | <<br>0.0001 | <<br>0.0001 | <<br>0.0001 |
| *syp-2Δ; cdc 77Δ* | < 0.0001 | <<br>0.0001 | < 0.0001 | – | <<br>0.0001 | <<br>0.0001 | <<br>0.0001 | <<br>0.0001 |
| *msh-6Δ* | 0.1734 | 0.0867 | < 0.0001 | < 0.0001 | – | 0.8242 | 0.9690 | 0.8807 |
| *msh-6Δ; cdc 77Δ* | 0.2734 | 0.1312 | < 0.0001 | < 0.0001 | 0.8242 | – | 0.7402 | 0.9132 |
| *lig-4Δ* | 0.1035 | 0.0404 | < 0.0001 | < 0.0001 | 0.9690 | 0.7402 | – | 0.7840 |
| *lig-4Δ; cdc 77Δ* | 0.1574 | 0.0552 | < 0.0001 | < 0.0001 | 0.8807 | 0.9132 | 0.7840 | – |

**Table S17.**
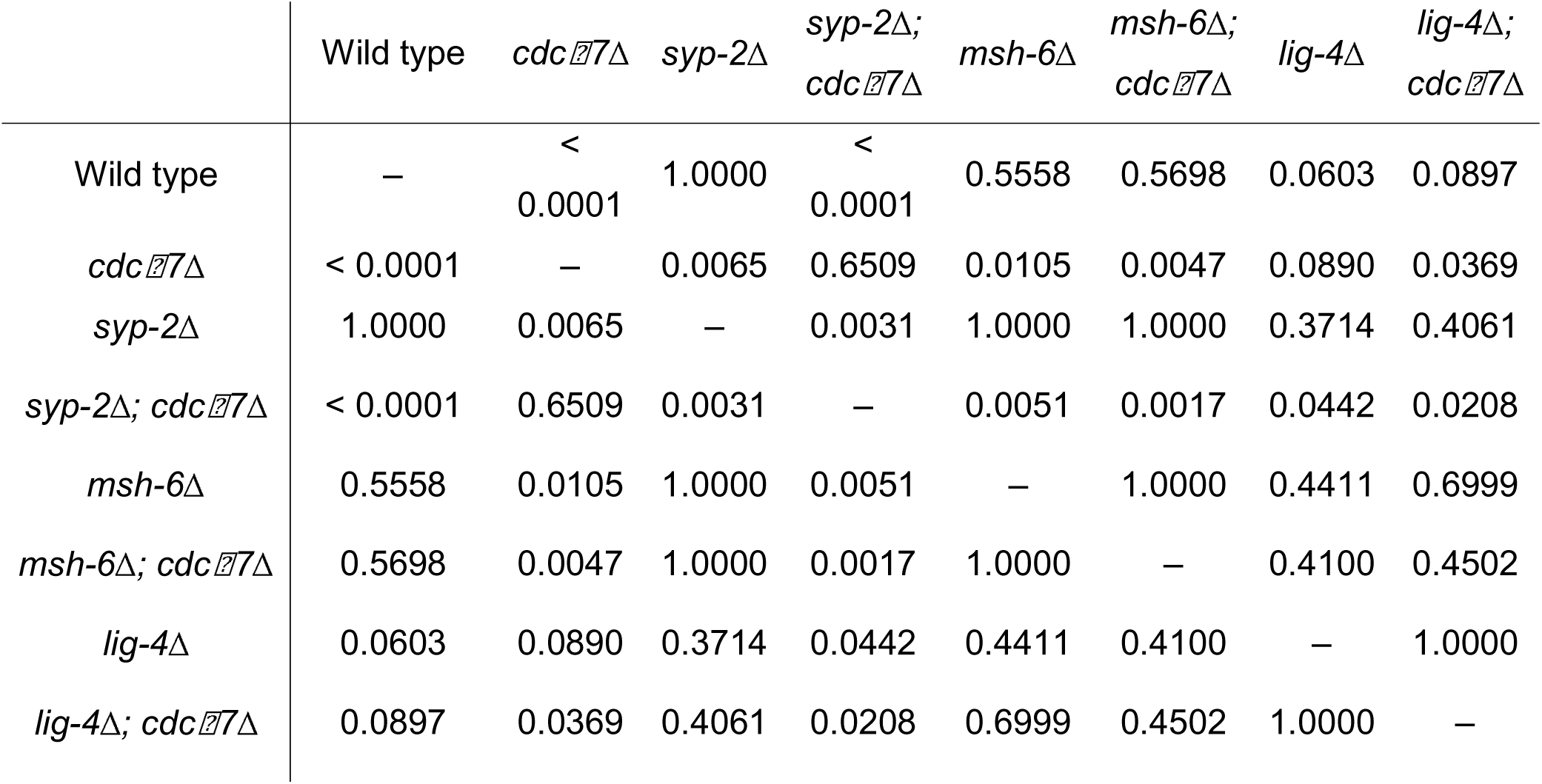
*P* values for pairwise Fisher’s exact tests for the proportion of nuclei with bridges at −1 diakinesis for the indicated genotypes.

|  | Wild type | *cdc 77Δ* | *syp-2Δ* | *syp-2Δ*;<br>*cdc 77Δ* | *msh-6Δ* | *msh-6Δ*;<br>*cdc 77Δ* | *lig-4Δ* | *lig-4Δ*;<br>*cdc 77Δ* |
| --- | --- | --- | --- | --- | --- | --- | --- | --- |
| Wild type | – | <<br>0.0001 | 1.0000 | <<br>0.0001 | 0.5558 | 0.5698 | 0.0603 | 0.0897 |
| *cdc 77Δ* | < 0.0001 | – | 0.0065 | 0.6509 | 0.0105 | 0.0047 | 0.0890 | 0.0369 |
| *syp-2Δ* | 1.0000 | 0.0065 | – | 0.0031 | 1.0000 | 1.0000 | 0.3714 | 0.4061 |
| *syp-2Δ*; *cdc 77Δ* | < 0.0001 | 0.6509 | 0.0031 | – | 0.0051 | 0.0017 | 0.0442 | 0.0208 |
| *msh-6Δ* | 0.5558 | 0.0105 | 1.0000 | 0.0051 | – | 1.0000 | 0.4411 | 0.6999 |
| *msh-6Δ*; *cdc 77Δ* | 0.5698 | 0.0047 | 1.0000 | 0.0017 | 1.0000 | – | 0.4100 | 0.4502 |
| *lig-4Δ* | 0.0603 | 0.0890 | 0.3714 | 0.0442 | 0.4411 | 0.4100 | – | 1.0000 |
| *lig-4Δ*; *cdc 77Δ* | 0.0897 | 0.0369 | 0.4061 | 0.0208 | 0.6999 | 0.4502 | 1.0000 | – |

**Table S18.** *P* values for the pairwise Chi-square test for the percentage of heterologous recombination events detected in the indicated genotypes.

|  | Wild<br>type | *cdc* 7<br>Δ | *msh*-<br>6Δ | *cdc* 7Δ<br>;<br>*msh*-6Δ | *lig*-4Δ | *cdc* 7Δ<br>;<br>*lig*-4Δ | *polq*-1 | *cdc* 7Δ<br>;<br>*polq*-1 |
| --- | --- | --- | --- | --- | --- | --- | --- | --- |
| Wild<br>type | – | 0.0005 | 0.0269 | 0.0002 | 0.016<br>6 | <<br>0.0001 |  | 0.0002 |
| *cdc* 7Δ | 0.0005 | – | 0.2204 | 0.7783 | 0.129<br>2 | 0.2192 | 0.004<br>2 | 0.7801 |
| *msh*-6Δ | 0.0269 | 0.2204 | – | 0.1657 | 0.846<br>9 | 0.0449 | 0.069<br>3 | 0.1691 |
| *cdc* 7Δ<br>;<br>*msh*-6Δ | 0.0002 | 0.7783 | 0.1657 | – | 0.085<br>3 | 0.3952 | 0.002<br>3 | 0.9948 |
| *lig*-4Δ | 0.0166 | 0.1292 | 0.8469 | 0.0853 | – | 0.0065 | 0.049<br>3 | 0.0918 |
| *cdc* 7Δ<br>;<br>*lig*-4Δ | <<br>0.0001 | 0.2192 | 0.0449 | 0.3952 | 0.006<br>5 | – | 0.000<br>3 | 0.4170 |
| *polq*-1 |  | 0.0042 | 0.0693 | 0.0023 | 0.049<br>3 | 0.0003 | – | 0.0023 |
| *cdc* 7Δ<br>;<br>*polq*-1 | 0.0002 | 0.7801 | 0.1691 | 0.9948 | 0.091<br>8 | 0.4170 | 0.002<br>3 | – |

**Table S19.** *P* values for the Mann–Whitney U test for DAPI body counts at −1 diakinesis for the indicated genotypes.

|  | Wild type | *cdc 7Δ* | *cdk 12*<sup>S30A</sup> | *cdc 7Δ*;<br>*cdk 12*<sup>S30A</sup> |
| --- | --- | --- | --- | --- |
| Wild type | – | 0.5690 | 0.0044 | 0.0990 |
| *cdc 7Δ* | 0.5690 | – | 0.0012 | 0.0354 |
| *cdk 12*<sup>S30A</sup> | 0.0044 | 0.0012 | – | 0.2732 |
| *cdc 7Δ*;<br>*cdk 12*<sup>S30A</sup> | 0.0990 | 0.0354 | 0.2732 | – |

**Table S20.** *P* values for pairwise Fisher’s exact tests for the proportion of nuclei with bridges at −1 diakinesis for the indicated genotypes.

|  | Wild type | *cdc 7Δ* | *cdk 12<sup>S30A</sup>* | *cdc 7Δ; cdk 12<sup>S30A</sup>* |
| --- | --- | --- | --- | --- |
| Wild type | – | < 0.0001 | 0.0002 | < 0.0001 |
| *cdc 7Δ* | < 0.0001 | – | 1.0000 | 0.8351 |
| *cdk 12<sup>S30A</sup>* | 0.0002 | 1.0000 | – | 1.0000 |
| *cdc 7Δ; cdk 12<sup>S30A</sup>* | < 0.0001 | 0.8351 | 1.0000 | – |

**Table S21.** *P* values for the Mann–Whitney U test for DAPI body counts at −1 diakinesis for the indicated genotypes.

|  | Wild type | *cdc 7Δ* | *cdk 12*<sup>S30D</sup> | *cdc 7Δ*;<br>*cdk 12*<sup>S30D</sup> |
| --- | --- | --- | --- | --- |
| Wild type | – | 0.5690 | 0.6718 | 1.0000 |
| *cdc 7Δ* | 0.5690 | – | 0.3969 | 0.6284 |
| *cdk 12*<sup>S30D</sup> | 0.6718 | 0.3969 | – | 0.7153 |
| *cdc 7Δ*;<br>*cdk 12*<sup>S30D</sup> | 1.0000 | 0.6284 | 0.7153 | – |

**Table S22.**
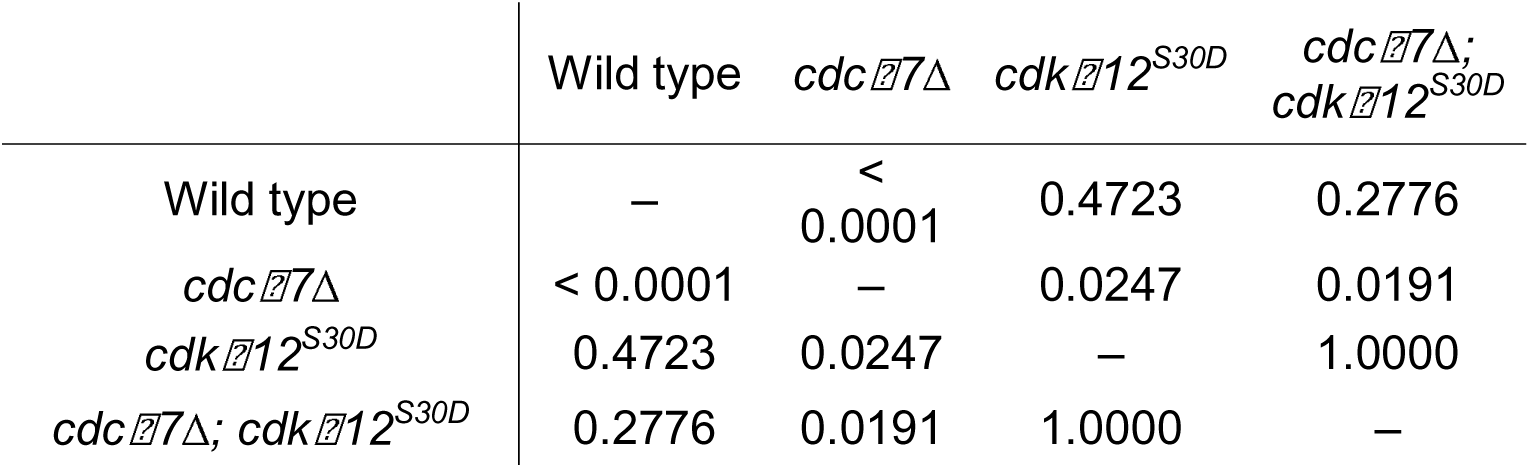
*P* values for pairwise Fisher’s exact tests for the proportion of nuclei with bridges at −1 diakinesis for the indicated genotypes.

**Table S23.** *P* values for the pairwise Chi-square test for the percentage of heterologous recombination events detected in the indicated genotypes.

|  | Wild<br>type | *cdc 7*Δ | *cdk 12*<sup>(S30D)</sup> | *cdc 7*;<br>*cdk 12*<sup>(S30D)</sup> |
| --- | --- | --- | --- | --- |
| Wild type | – | 0.0005 | 0.2800 | 0.0651 |
| *cdc 7*Δ | 0.0005 | – | 0.0052 | 0.0365 |
| *cdk 12*<sup>(S30D)</sup> | 0.2800 | 0.0052 | – | 0.3313 |
| *cdc 7*;<br>*cdk 12*<sup>(S30D)</sup> | 0.0651 | 0.0365 | 0.3313 | – |

**Table S24.** DNA–2 phosphosite assignments from an independent phosphoscreen. Identified phosphopeptides, phosphosite localization probabilities, and associated peptide-spectrum match statistics are listed. PEP denotes posterior error probability.

| Site | Phosphopeptide sequence | Localization probability | PEP |
| --- | --- | --- | --- |
| S46 | GTEHDGDSVDSFENFCPT(0.099)AS(0.451)S(0.451)PK | 0.45 | 1.30E-21 |
| S47 | GTEHDGDSVDSFENFCPT(0.099)AS(0.451)S(0.451)PK | 0.45 | 1.30E-21 |
| T383 | DGAS(0.02)HCVT(0.98)PR | 0.98 | 3.36E-03 |

**Table S25.** *P* values for the Mann–Whitney U test for DAPI body counts at −1 diakinesis for the indicated genotypes.

|  | Wild<br>type | *cdc 7*Δ | *cdk 12*<sup>S30A</sup> | *dna 2*<sup>S46A,</sup><br><sup>S47A</sup> | *cdk 12*<sup>S30A</sup>,<br>*dna 2*<sup>S46A,</sup><br><sup>S47A</sup> | *cdc 7*Δ;<br>*dna 2*<sup>S46A, S47A</sup> |
| --- | --- | --- | --- | --- | --- | --- |
| Wild type | – | 0.5690 | 0.0044 | 0.0009 | 0.0115 | 0.4135 |
| *cdc 7*Δ | 0.5690 | – | 0.0012 | 0.0002 | 0.0032 | 0.1987 |
| *cdk 12*<sup>S30A</sup> | 0.0044 | 0.0012 | – | 0.6601 | 0.7665 | 0.0689 |
| *dna 2*<sup>S46A, S47A</sup> | 0.0009 | 0.0002 | 0.6601 | – | 0.4540 | 0.0250 |
| *cdk 12*<sup>S30A</sup>,<br>*dna 2*<sup>S46A, S47A</sup> | 0.0115 | 0.0032 | 0.7665 | 0.4540 | – | 0.1297 |
| *cdc 7*Δ;<br>*dna 2*<sup>S46A, S47A</sup> | 0.4135 | 0.1987 | 0.0689 | 0.0250 | 0.1297 | – |

**Table S26.** *P* values for pairwise Fisher’s exact tests for the proportion of nuclei with bridges at −1 diakinesis for the indicated genotypes.

|  | Wild type | *cdc 7*Δ | *cdk 12*<sup>S30A</sup> | *dna 2*<sup>S46A, S47A</sup> | *cdk 12*<sup>S30A</sup>,<br>*dna 2*<sup>S46A, S47A</sup> | *cdc 7*Δ;<br>*dna 2*<sup>S46A, S47A</sup> |
| --- | --- | --- | --- | --- | --- | --- |
| Wild type | – | <<br>0.0001 | 0.0002 | 0.0001 | 0.0018 | 0.0091 |
| *cdc 7*Δ | < 0.0001 | – | 1.0000 | 1.0000 | 0.6550 | 0.3505 |
| *cdk 12*<sup>S30A</sup> | 0.0002 | 1.0000 | – | 1.0000 | 0.6061 | 0.4099 |
| *dna 2*<sup>S46A, S47A</sup> | 0.0001 | 1.0000 | 1.0000 | – | 0.6230 | 0.3041 |
| *cdk 12*<sup>S30A</sup>,<br>*dna 2*<sup>S46A, S47A</sup> | 0.0018 | 0.6550 | 0.6061 | 0.6230 | – | 0.7805 |
| *cdc 7*Δ;<br>*dna 2*<sup>S46A, S47A</sup> | 0.0091 | 0.3505 | 0.4099 | 0.3041 | 0.7805 | – |

**Table S27.** *P* values for the Mann–Whitney U test for DAPI body counts at −1 diakinesis for the indicated genotypes.

|  | Wild type | *cdc 7Δ* | *dna 2*<sup>S46D,</sup><br><sup>S47D</sup> | *cdk 12*<sup>S30A,</sup><br><sup>dna 2<sup>S46D,</sup></sup><br><sup><sup>S47D</sup></sup> | *cdc 7Δ;*<br>*dna 2*<sup>S46D, S47D</sup> |
| --- | --- | --- | --- | --- | --- |
| Wild type | – | 0.5690 | 0.0834 | 0.2374 | 0.1946 |
| *cdc 7Δ* | 0.5690 | – | 0.0309 | 0.1024 | 0.0796 |
| *dna 2*<sup>S46D, S47D</sup> | 0.0834 | 0.0309 | – | 0.6113 | 0.6439 |
| *cdk 12*<sup>S30A,</sup><br><sup>dna 2<sup>S46D, S47D</sup></sup> | 0.2374 | 0.1024 | 0.6113 | – | 0.9512 |
| *cdc 7Δ;*<br>*dna 2*<sup>S46D, S47D</sup> | 0.1946 | 0.0796 | 0.6439 | 0.9512 | – |

**Table S28.**
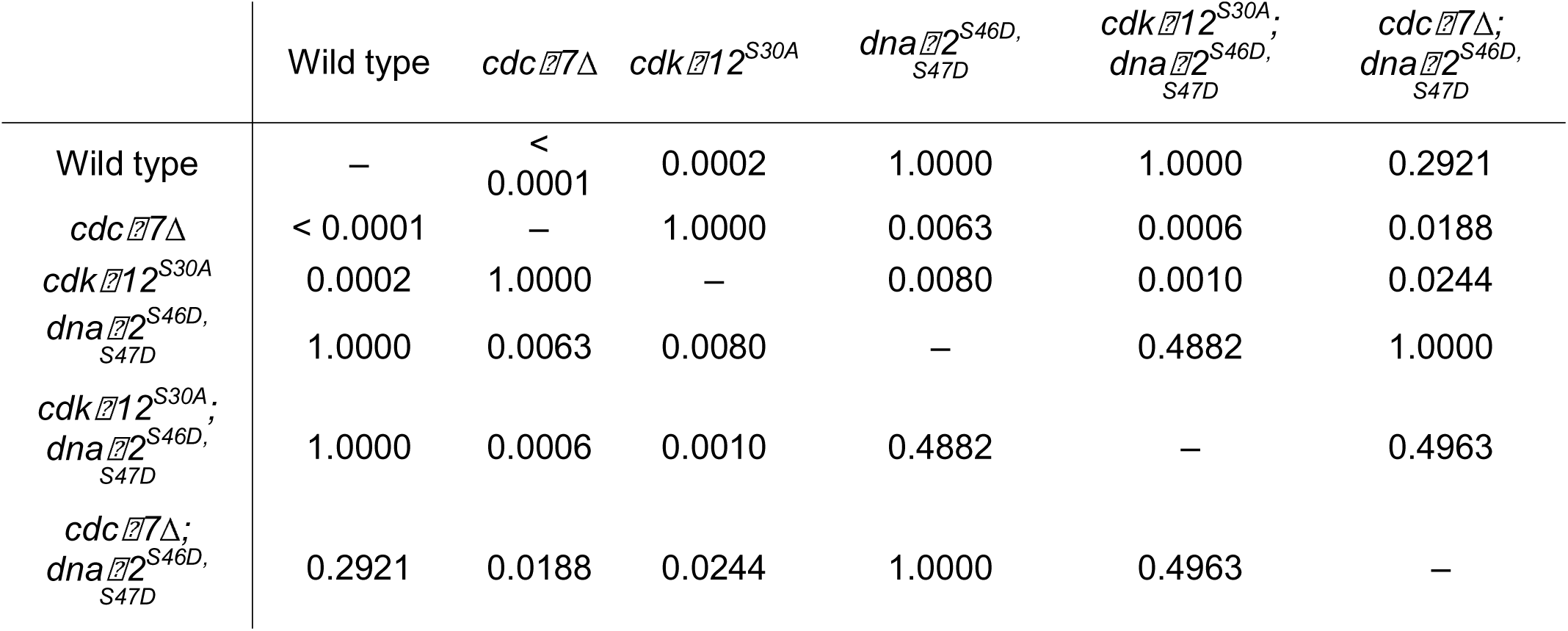
*P* values for pairwise Fisher’s exact tests for the proportion of nuclei with bridges at −1 diakinesis for the indicated genotypes.

|  | Wild type | *cdc 7Δ* | *cdk 12<sup>S30A</sup>* | *dna 2<sup>S46D, S47D</sup>* | *cdk 12<sup>S30A</sup>,*<br>*dna 2<sup>S46D, S47D</sup>* | *cdc 7Δ;*<br>*dna 2<sup>S46D, S47D</sup>* |
| --- | --- | --- | --- | --- | --- | --- |
| Wild type | – | <<br>0.0001 | 0.0002 | 1.0000 | 1.0000 | 0.2921 |
| *cdc 7Δ* | < 0.0001 | – | 1.0000 | 0.0063 | 0.0006 | 0.0188 |
| *cdk 12<sup>S30A</sup>* | 0.0002 | 1.0000 | – | 0.0080 | 0.0010 | 0.0244 |
| *dna 2<sup>S46D, S47D</sup>* | 1.0000 | 0.0063 | 0.0080 | – | 0.4882 | 1.0000 |
| *cdk 12<sup>S30A</sup>,*<br>*dna 2<sup>S46D, S47D</sup>* | 1.0000 | 0.0006 | 0.0010 | 0.4882 | – | 0.4963 |
| *cdc 7Δ;*<br>*dna 2<sup>S46D, S47D</sup>* | 0.2921 | 0.0188 | 0.0244 | 1.0000 | 0.4963 | – |

**Table S29.**
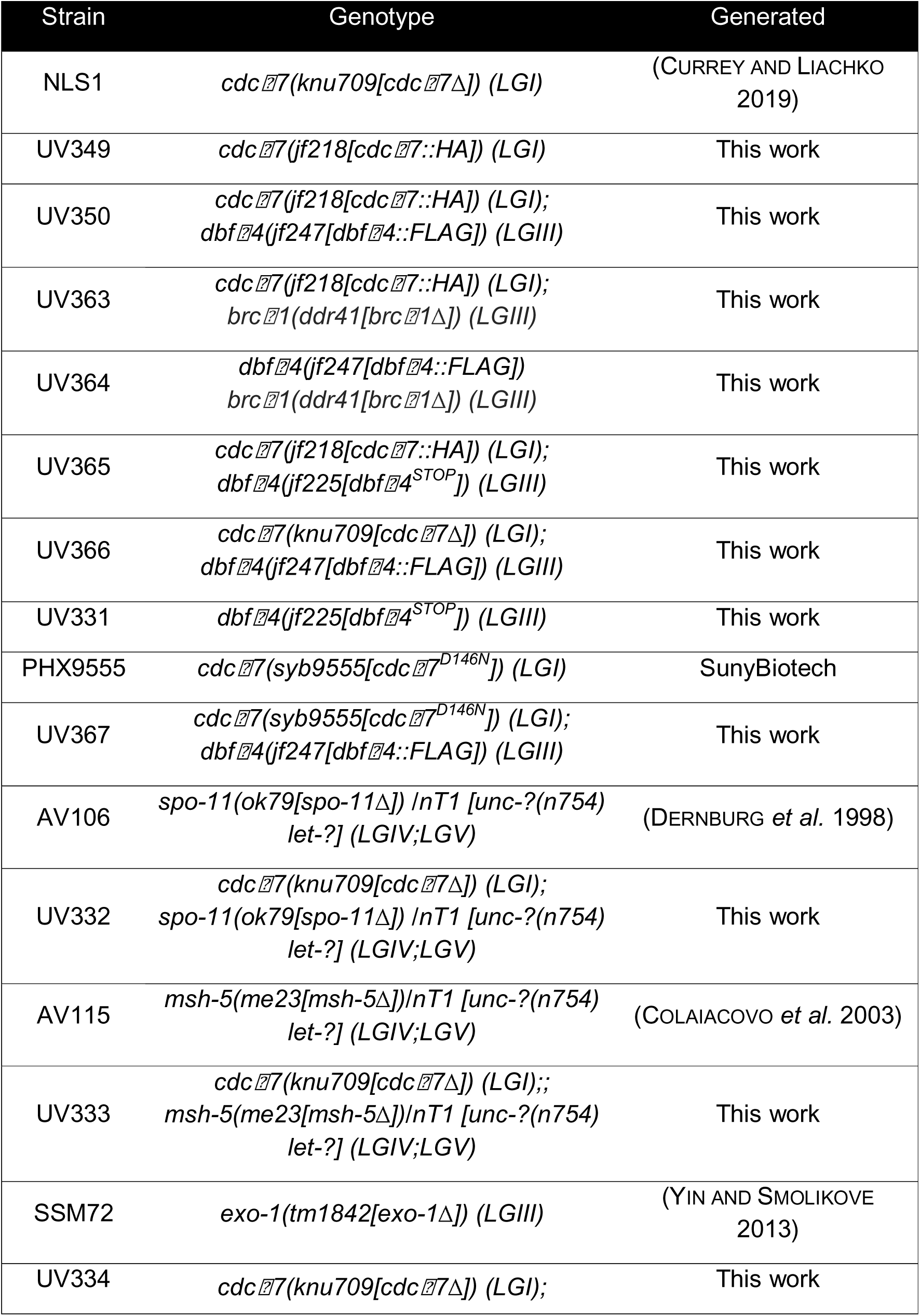
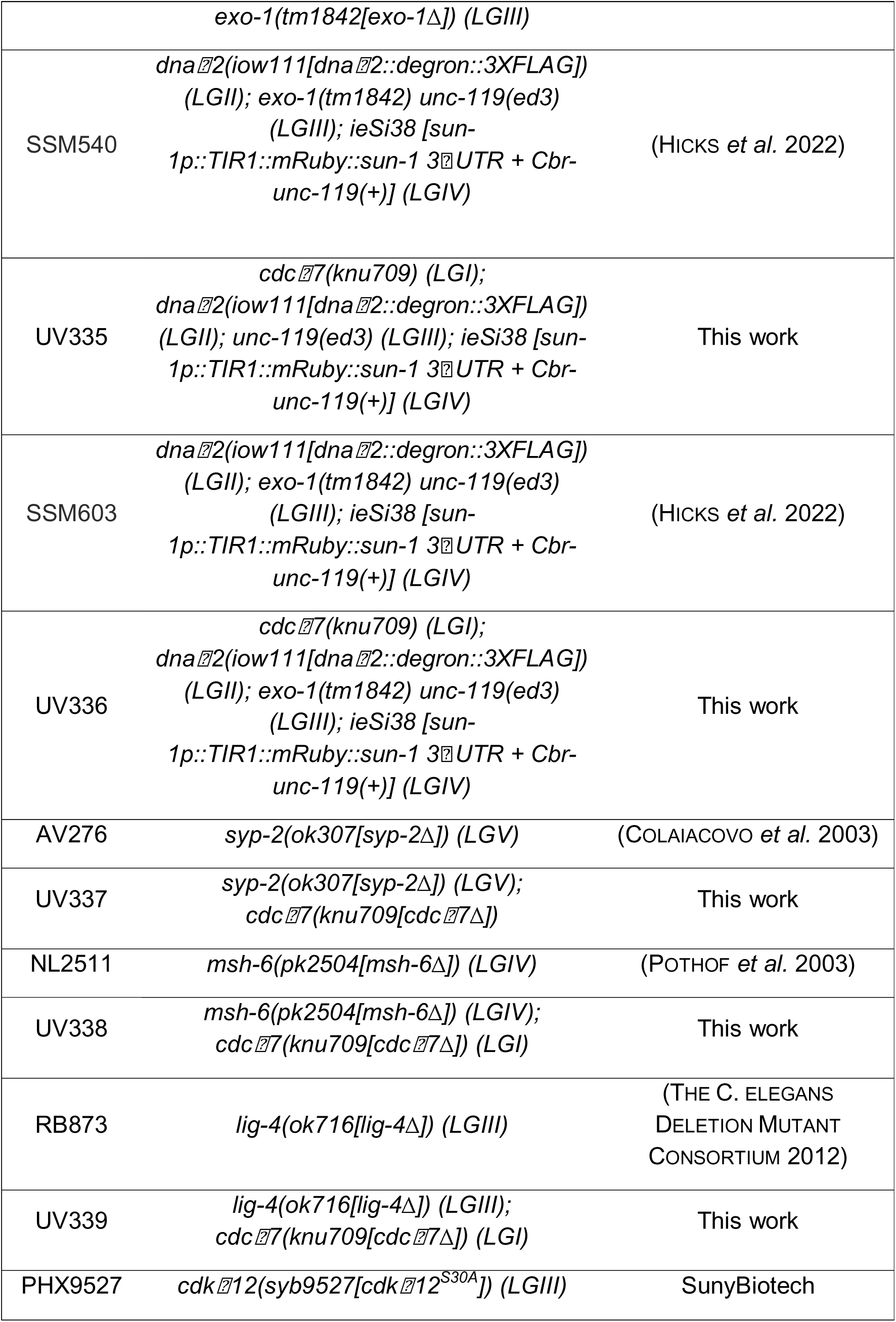
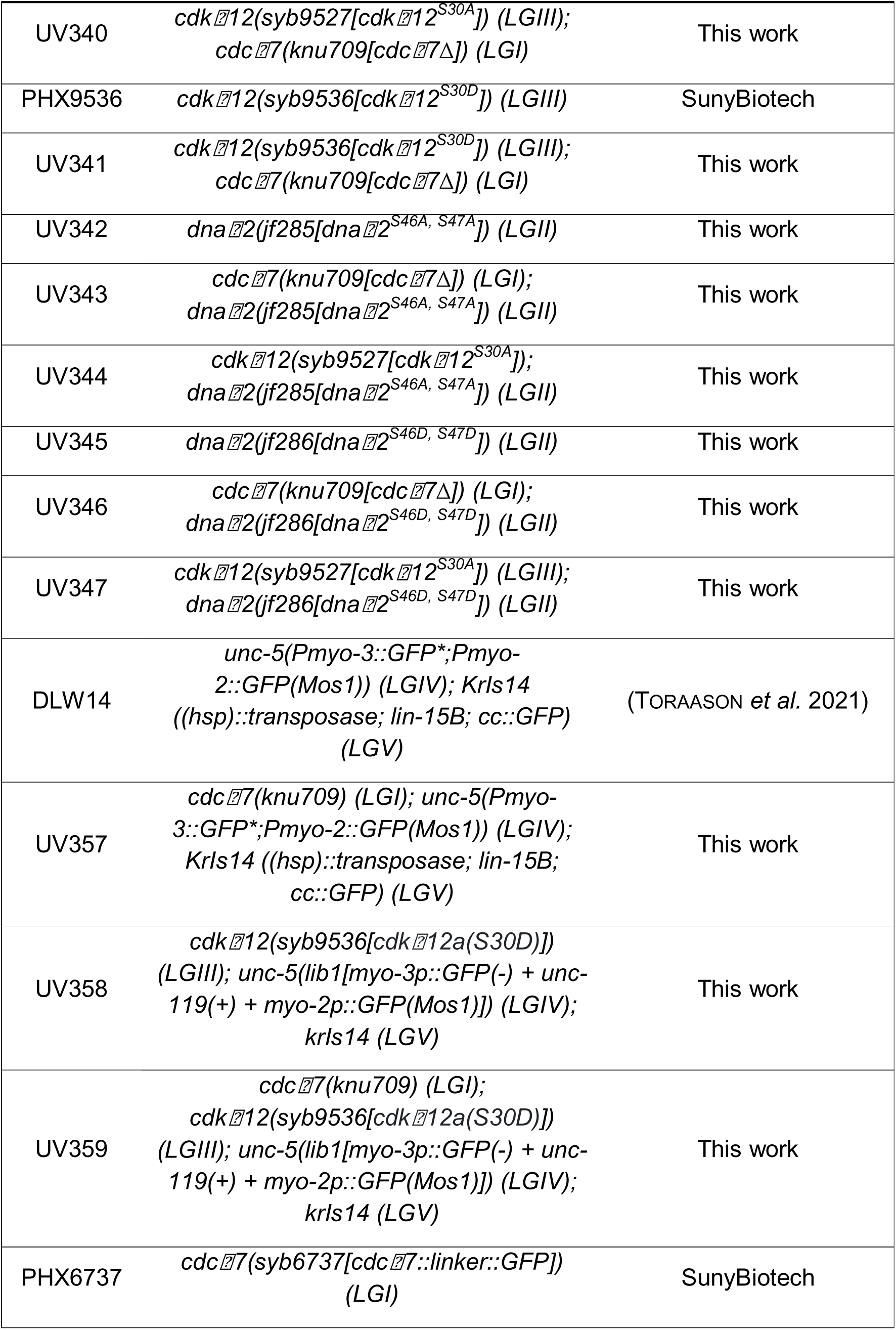
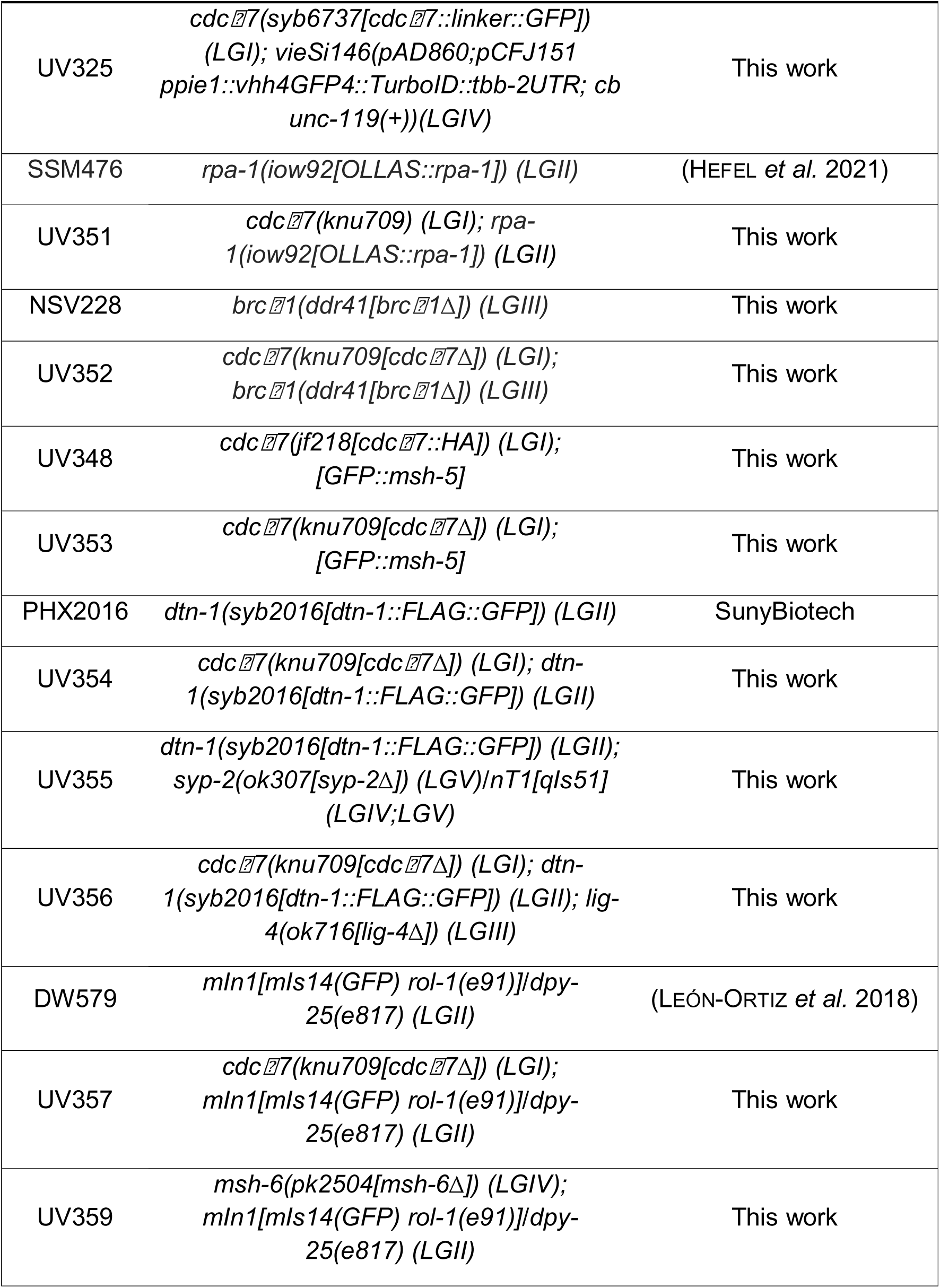
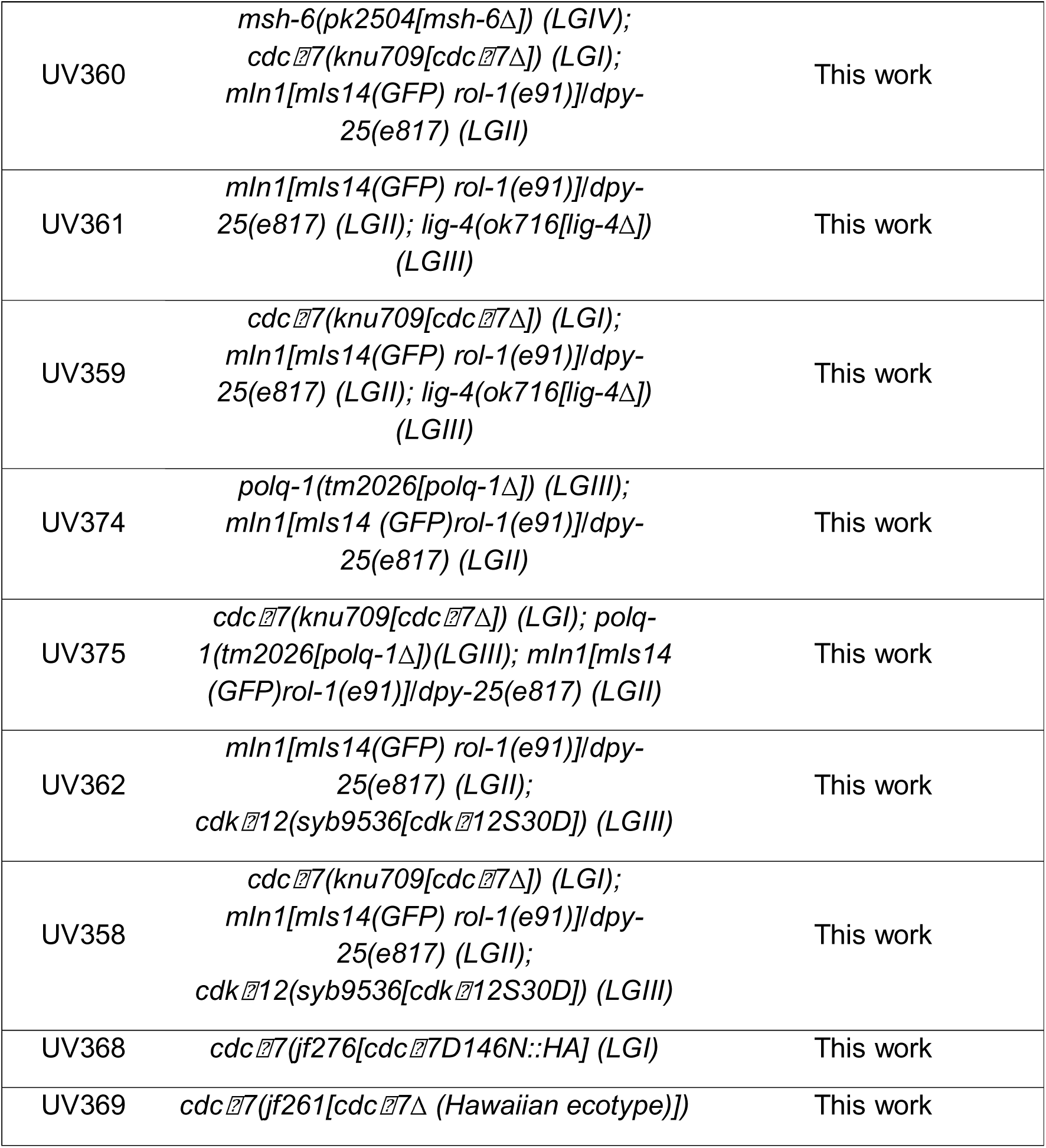
Strains generated and used for this study.

**Table S30.**
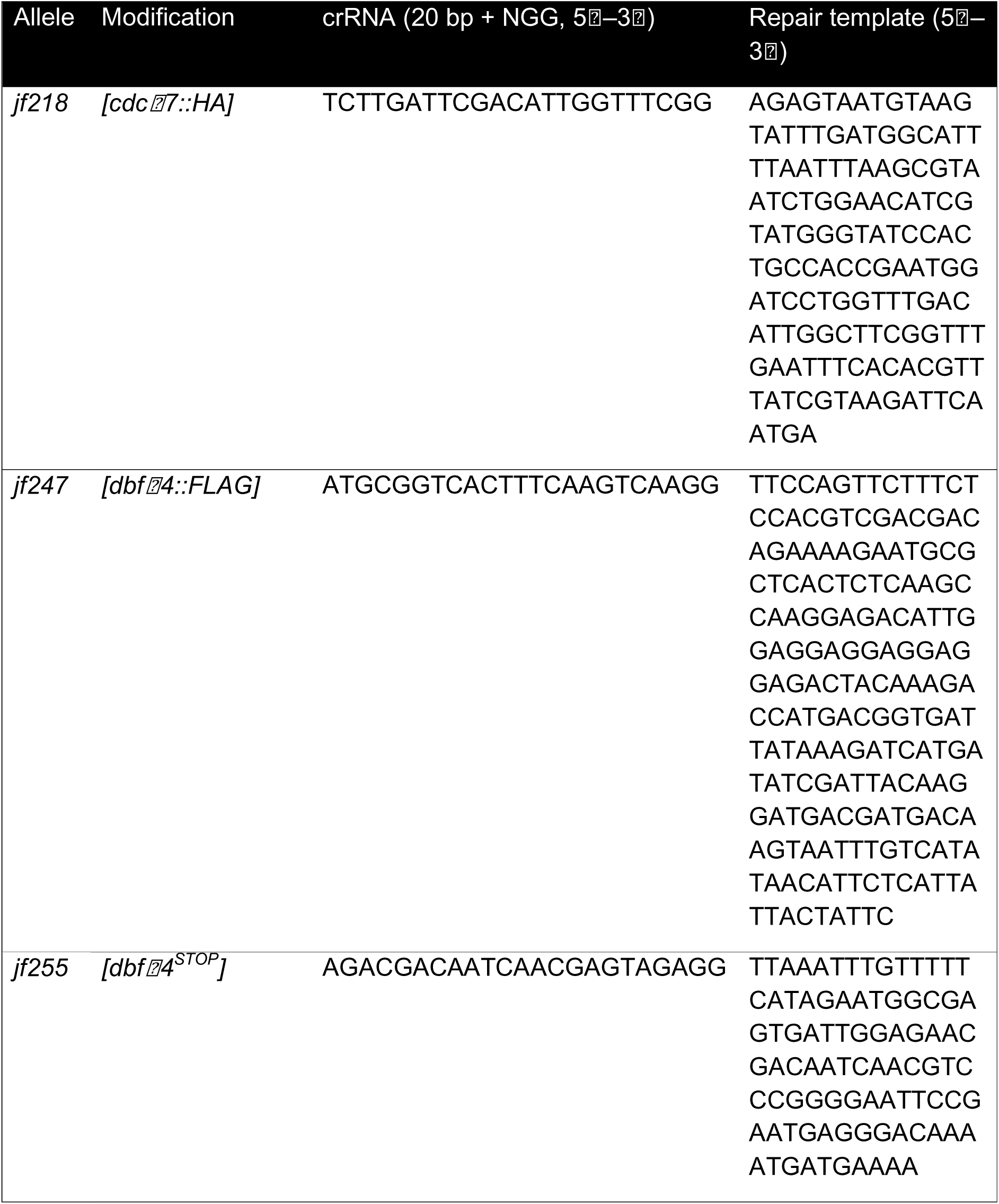
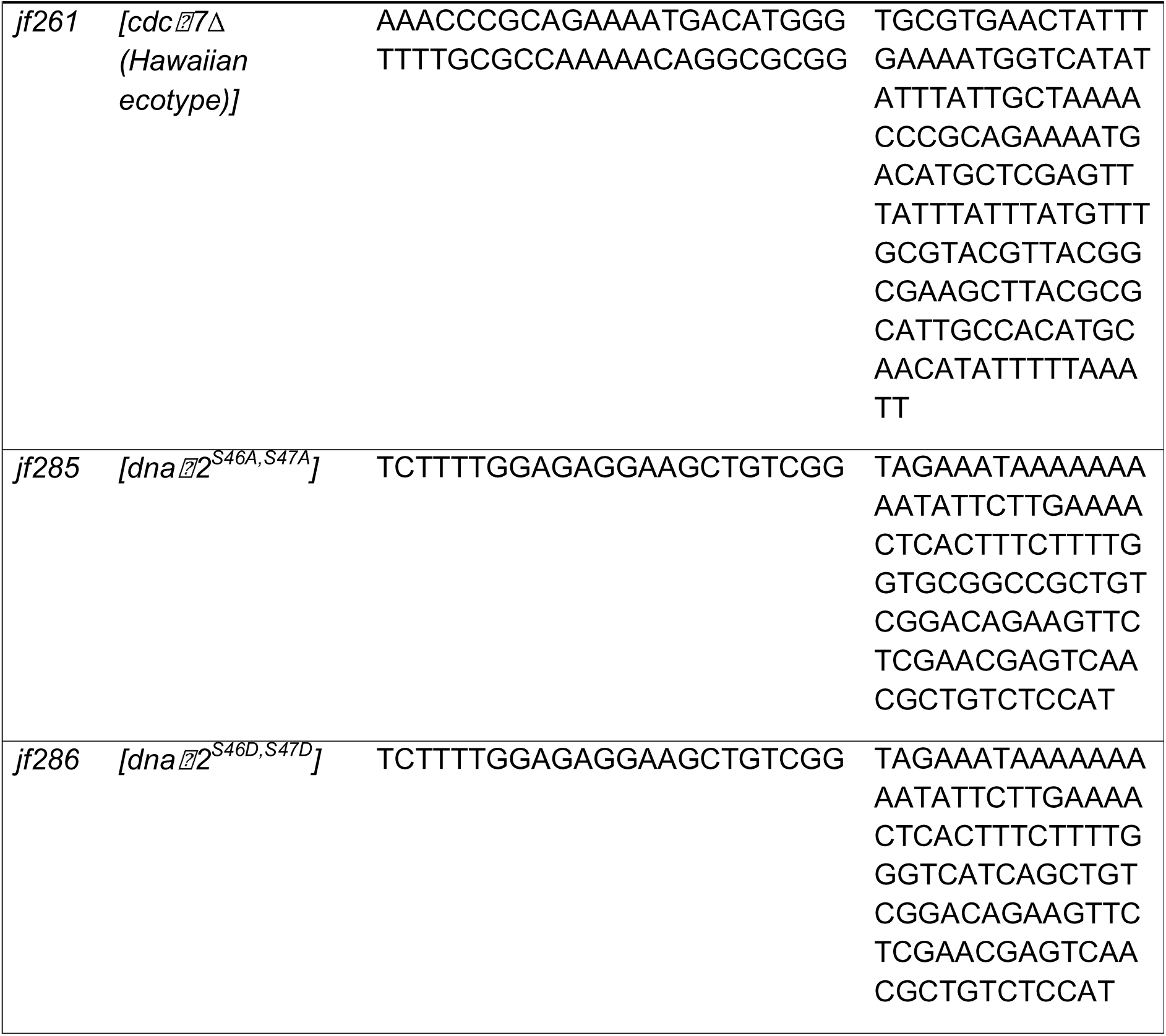
crRNAs, repair templates, and primers designed for this study.

**Table S31.**
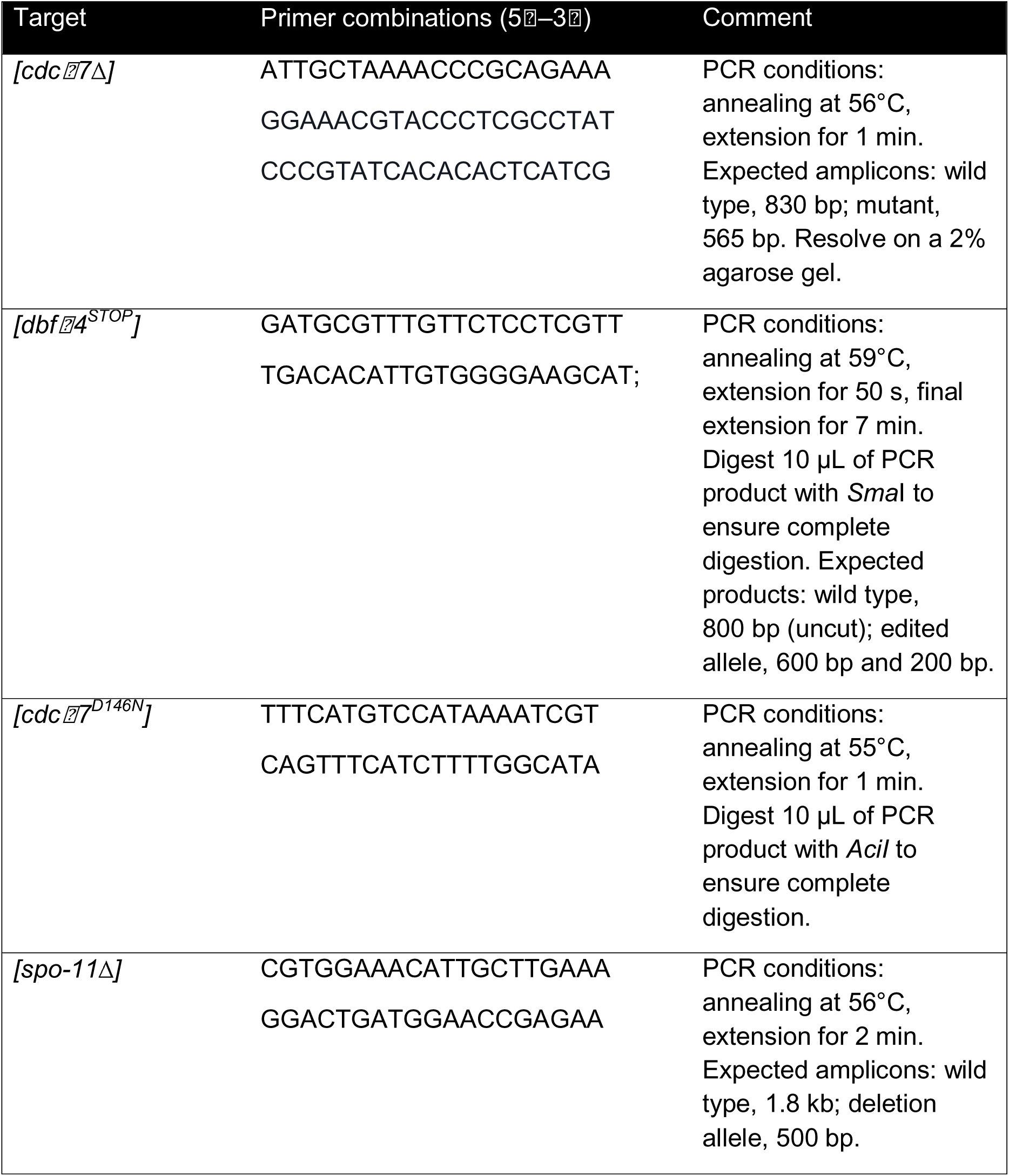
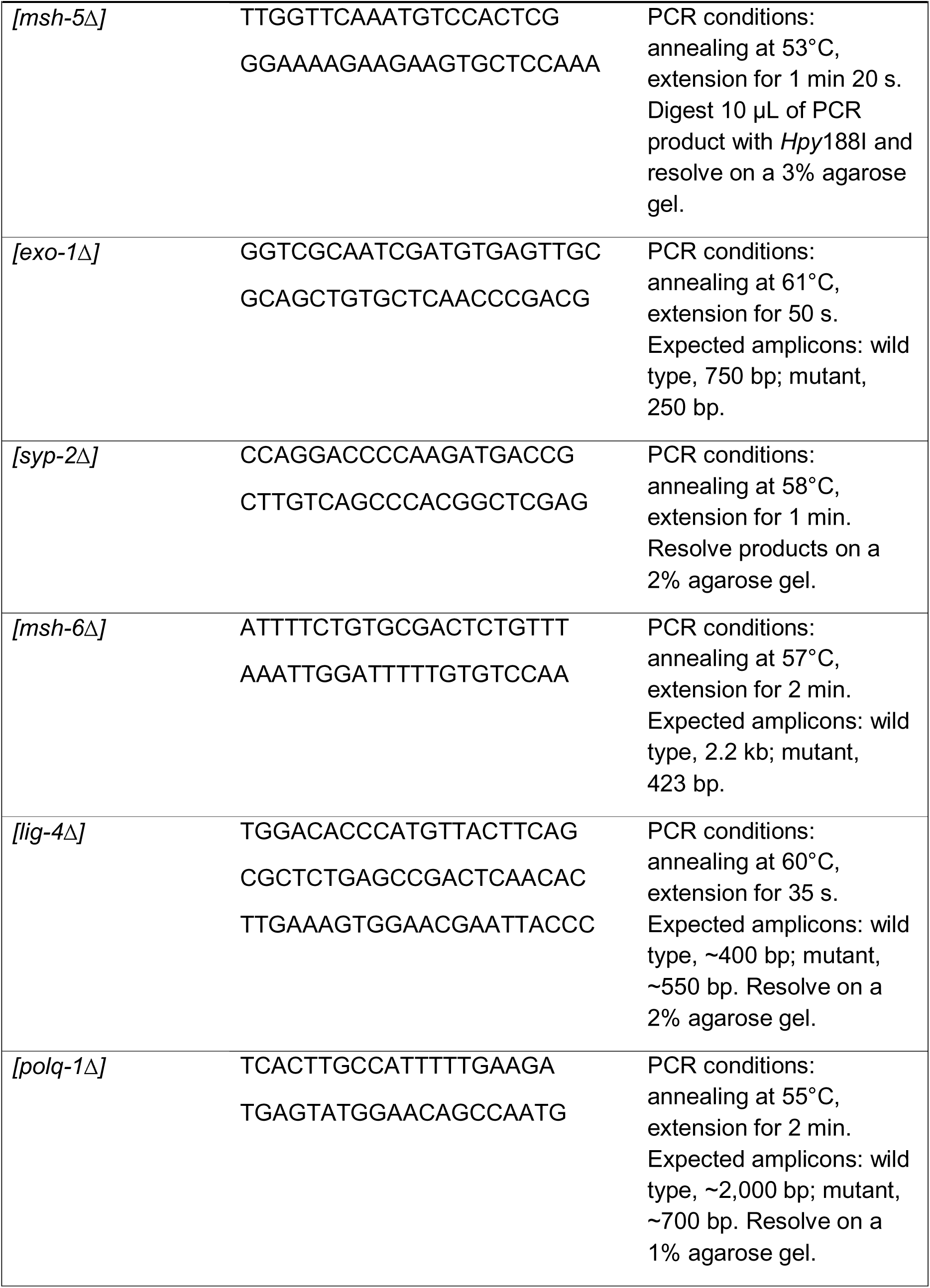
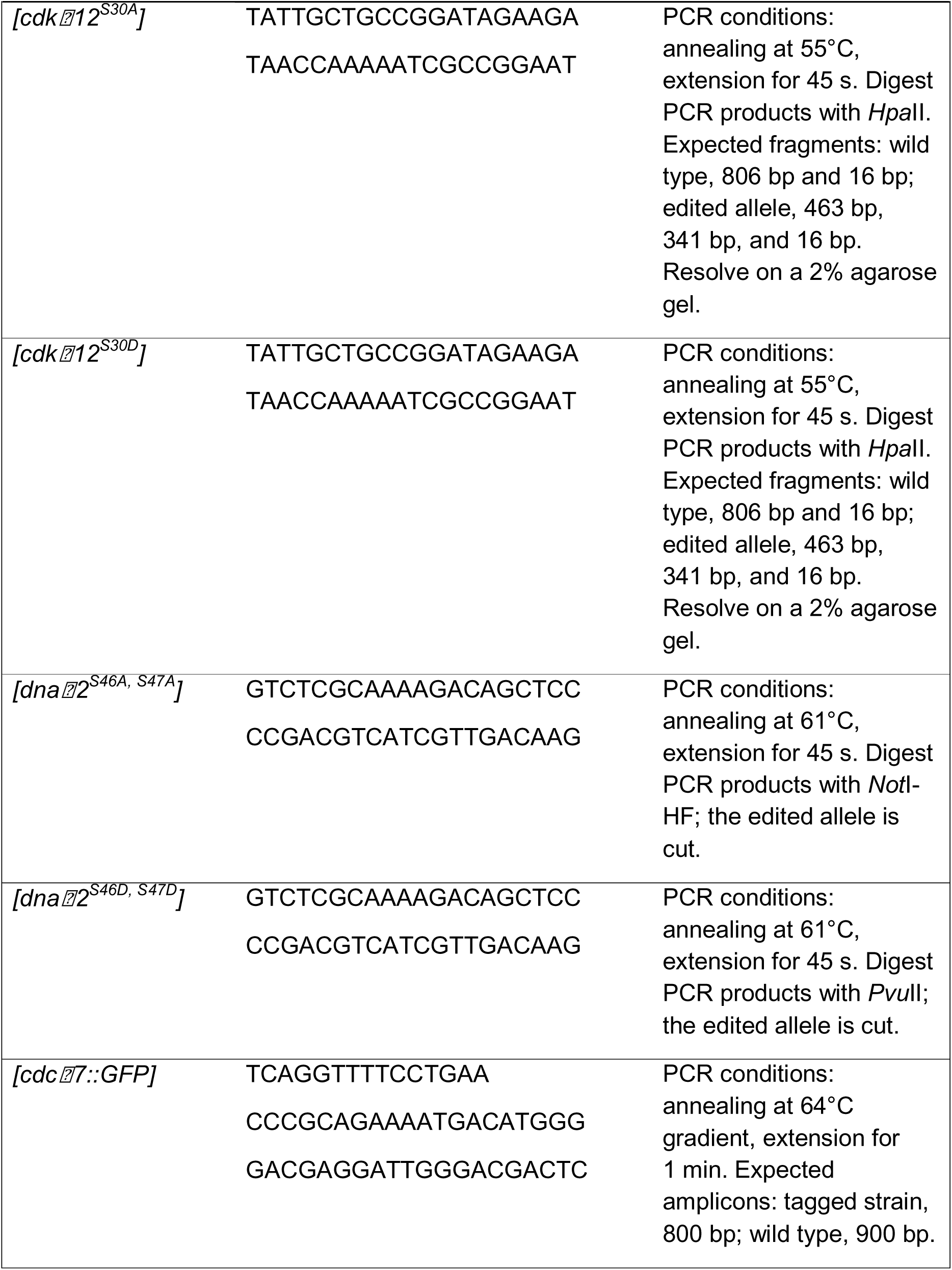
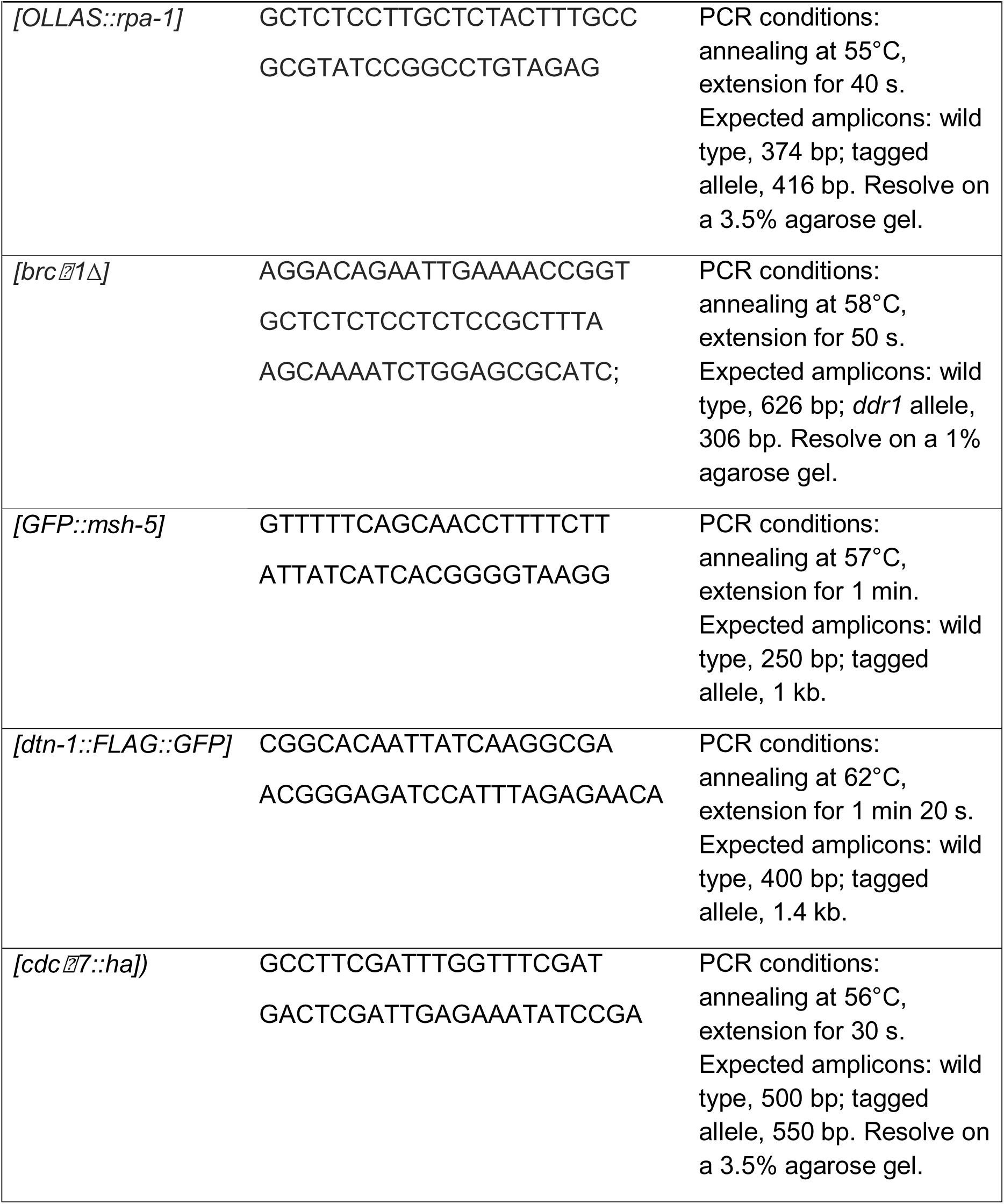
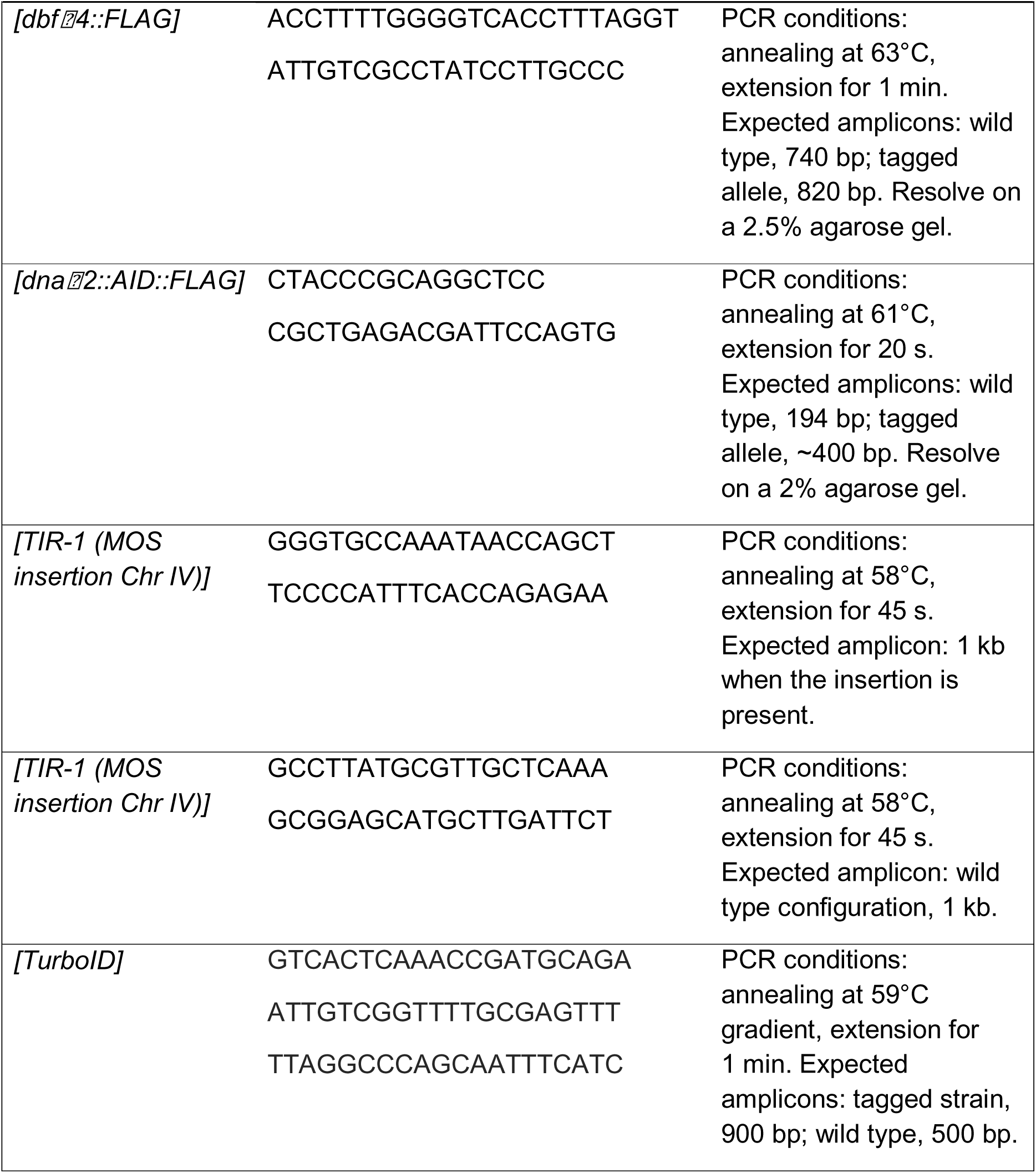
Primers used for this study.

**Table S32.**
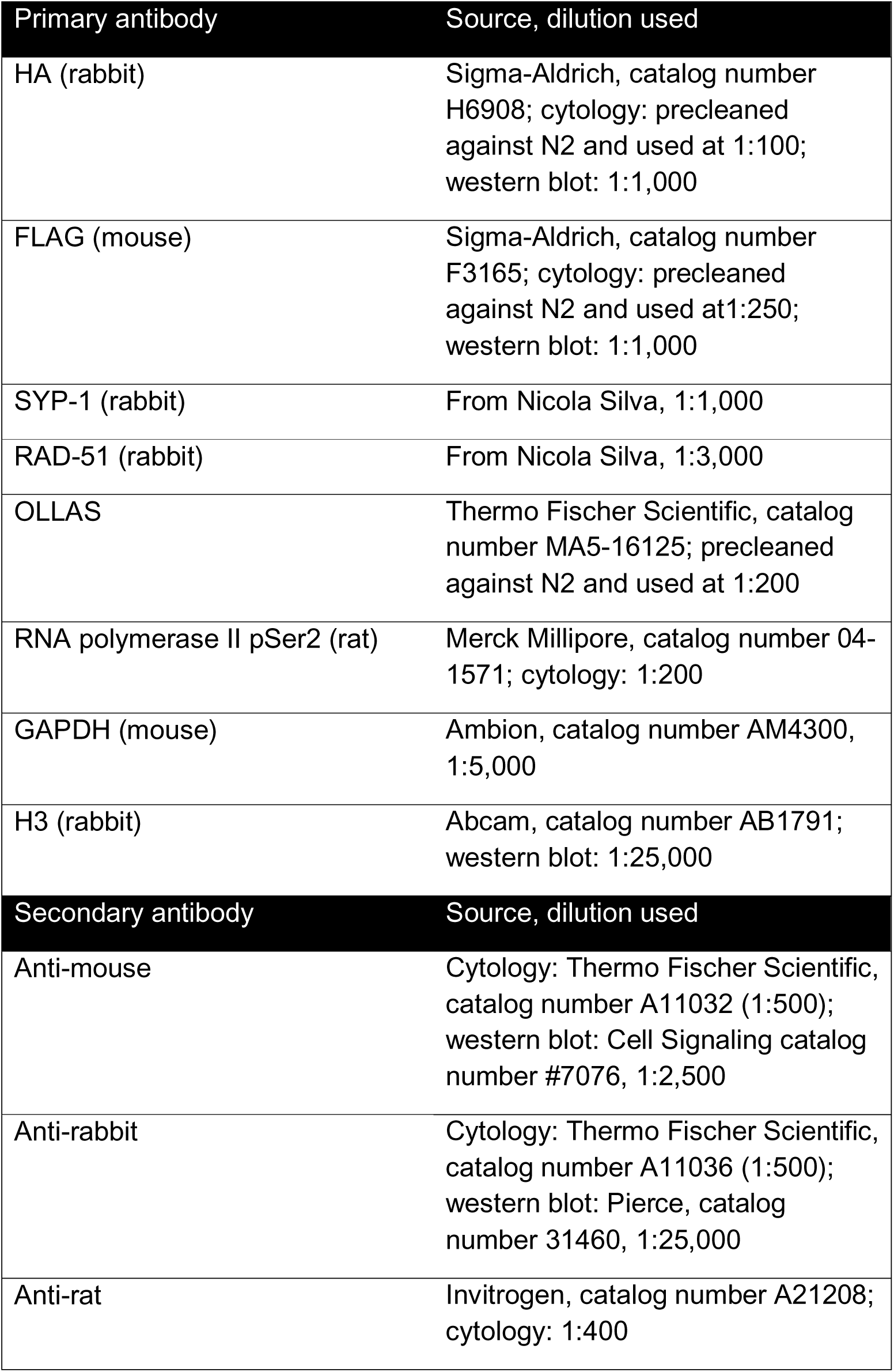
Antibodies used in this study.

| Primary antibody | Source, dilution used |
| --- | --- |
| HA (rabbit) | Sigma-Aldrich, catalog number H6908; cytology: precleaned against N2 and used at 1:100; western blot: 1:1,000 |
| FLAG (mouse) | Sigma-Aldrich, catalog number F3165; cytology: precleaned against N2 and used at 1:250; western blot: 1:1,000 |
| SYP-1 (rabbit) | From Nicola Silva, 1:1,000 |
| RAD-51 (rabbit) | From Nicola Silva, 1:3,000 |
| OLLAS | Thermo Fischer Scientific, catalog number MA5-16125; precleaned against N2 and used at 1:200 |
| RNA polymerase II pSer2 (rat) | Merck Millipore, catalog number 04-1571; cytology: 1:200 |
| GAPDH (mouse) | Ambion, catalog number AM4300, 1:5,000 |
| H3 (rabbit) | Abcam, catalog number AB1791; western blot: 1:25,000 |
| Secondary antibody | Source, dilution used |
| Anti-mouse | Cytology: Thermo Fischer Scientific, catalog number A11032 (1:500); western blot: Cell Signaling catalog number #7076, 1:2,500 |
| Anti-rabbit | Cytology: Thermo Fischer Scientific, catalog number A11036 (1:500); western blot: Pierce, catalog number 31460, 1:25,000 |
| Anti-rat | Invitrogen, catalog number A21208; cytology: 1:400 |

**Table S33.**
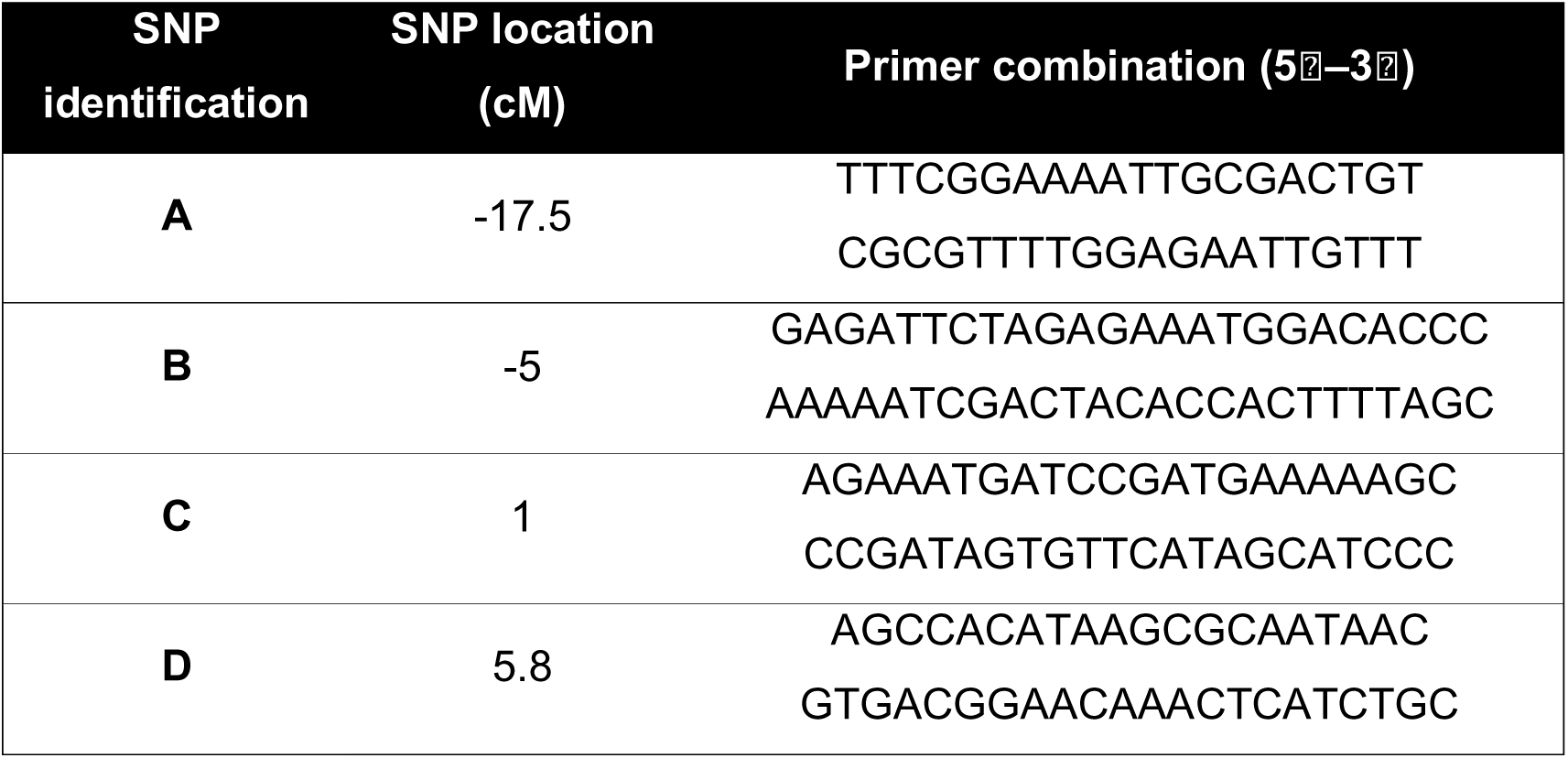
Primers and SNP locations on chromosome V used in the homologous recombination assay. All primers were designed to have a melting temperature (Tm) of 60□°C and a 1□min extension time. PCR products were digested with *Dra*I.

## References

Abramson, J., J. Adler, J. Dunger, R. Evans, T. Green et al., 2024 Accurate structure prediction of biomolecular interactions with AlphaFold 3. Nature 630: 493–500.

Achache, H., L. Laurent, Y. Hecker-Mimoun, H. Ishtayeh, Y. Rappaport et al., 2019 Progression of Meiosis Is Coordinated by the Level and Location of MAPK Activation Via OGR-2 in Caenorhabditis elegans. Genetics 212: 213–229.

Adamo, A., S. J. Collis, C. A. Adelman, N. Silva, Z. Horejsi et al., 2010 Preventing Nonhomologous End Joining Suppresses DNA Repair Defects of Fanconi Anemia. Molecular Cell 39: 25–35.

Alpi, A., P. Pasierbek, A. Gartner and J. Loidl, 2003 Genetic and cytological characterization of the recombination protein RAD-51 in Caenorhabditis elegans. Chromosoma 112: 6–16.

Altschul, S. F., T. L. Madden, A. A. Schäffer, J. Zhang, Z. Zhang et al., 1997 Gapped BLAST and PSI-BLAST: a new generation of protein database search programs. Nucleic Acids Research 25: 3389–3402.

Argunhan, B., T. Tsubouchi and H. Tsubouchi, 2018 Polo is not solo in meiosis. Cell Cycle 17: 273–274.

Arur, S., M. Ohmachi, M. Berkseth, S. Nayak, D. Hansen et al., 2011 MPK-1 ERK controls membrane organization in C. elegans oogenesis via a sex-determination module. Dev Cell 20: 677–688.

Baudrimont, A., and V. Jantsch, 2026 DDK meiotic resection analysis, pp. Zenodo.

Baumann, P., F. E. Benson and S. C. West, 1996 Human Rad51 Protein Promotes ATP-Dependent Homologous Pairing and Strand Transfer Reactions In Vitro. Cell 87: 757–766.

Belan, O., R. Anand and S. J. Boulton, 2021 Mechanism of mitotic recombination: insights from C. elegans. Curr Opin Genet Dev 71: 10–18.

Boulton, S. J., J. S. Martin, J. Polanowska, D. E. Hill, A. Gartner et al., 2004 BRCA1/BARD1 orthologs required for DNA repair in Caenorhabditis elegans. Curr Biol 14: 33–39.

Bowman, E. A., C. R. Bowman, J. H. Ahn and W. G. Kelly, 2013 Phosphorylation of RNA polymerase II is independent of P-TEFb in the C. elegans germline. Development 140: 3703–3713.

Brandt, J. N., K. A. Hussey and Y. Kim, 2020 Spatial and temporal control of targeting Polo-like kinase during meiotic prophase. J Cell Biol 219.

Brenner, S., 1974 The genetics of Caenorhabditis elegans. Genetics 77: 71–94.

Bruck, I., and D. Kaplan, 2009 Dbf4-Cdc7 phosphorylation of Mcm2 is required for cell growth. J Biol Chem 284: 28823–28831.

Cannavo, E., and P. Cejka, 2014 Sae2 promotes dsDNA endonuclease activity within Mre11–Rad50–Xrs2 to resect DNA breaks. Nature 514: 122–125.

Castellano-Pozo, M., S. Pacheco, G. Sioutas, A. L. Jaso-Tamame, M. H. Dore et al., 2020 Surveillance of cohesin-supported chromosome structure controls meiotic progression. Nat Commun 11: 4345.

Ceccaldi, R., and P. Cejka, 2025 Mechanisms and regulation of DNA end resection in the maintenance of genome stability. Nat Rev Mol Cell Biol 26: 586–599.

Ceccaldi, R., B. Rondinelli and A. D. D’Andrea, 2016 Repair Pathway Choices and Consequences at the Double-Strand Break. Trends in Cell Biology 26: 52–64.

Cejka, P., 2015 DNA End Resection: Nucleases Team Up with the Right Partners to Initiate Homologous Recombination *. Journal of Biological Chemistry 290: 22931–22938.

Ceppi, I., M. R. Dello Stritto, M. Mütze, S. Braunshier, V. Mengoli et al., 2024 Mechanism of BRCA1–BARD1 function in DNA end resection and DNA protection. Nature 634: 492–500.

Chen, X., H. Niu, W.-H. Chung, Z. Zhu, A. Papusha et al., 2011 Cell cycle regulation of DNA double-strand break end resection by Cdk1-dependent Dna2 phosphorylation. Nature Structural & Molecular Biology 18: 1015–1019.

Chen, X., R. T. Suhandynata, R. Sandhu, B. Rockmill, N. Mohibullah et al., 2015 Phosphorylation of the Synaptonemal Complex Protein Zip1 Regulates the Crossover/Noncrossover Decision during Yeast Meiosis. PLoS Biol 13: e1002329.

Chin, G. M., and A. M. Villeneuve, 2001 C. elegans mre-11 is required for meiotic recombination and DNA repair but is dispensable for the meiotic G2 DNA damage checkpoint. Genes & Development 15: 522–534.

Colaiacovo, M. P., A. J. MacQueen, E. Martinez-Perez, K. McDonald, A. Adamo et al., 2003 Synaptonemal complex assembly in C. elegans is dispensable for loading strand-exchange proteins but critical for proper completion of recombination. Dev Cell 5: 463–474.

Colaiácovo, M. P., A. J. MacQueen, E. Martinez-Perez, K. McDonald, A. Adamo et al., 2003 Synaptonemal Complex Assembly in *C. elegans* Is Dispensable for Loading Strand-Exchange Proteins but Critical for Proper Completion of Recombination. Developmental Cell 5: 463–474.

Cole, F., L. Kauppi, J. Lange, I. Roig, R. Wang et al., 2012 Homeostatic control of recombination is implemented progressively in mouse meiosis. Nature Cell Biology 14: 424–430.

Currey, H. N., and N. F. Liachko, 2019 A CRISPR/Cas9-generated cdc-7 loss of function mutation does not cause temperature-dependent fertility defects.

Davis, M. W., M. Hammarlund, T. Harrach, P. Hullett, S. Olsen et al., 2005 Rapid single nucleotide polymorphism mapping in C. elegans. BMC Genomics 6: 118.

Dello Stritto, M. R., N. Vojtassakova, M. Velkova, P. Hamminger, P. Ulm et al., 2022 The topoisomerase 3 zinc finger domain cooperates with the RMI1 scaffold to promote stable association of the BTR complex to recombination intermediates in the Caenorhabditis elegans germline. Nucleic Acids Res 50: 5652–5671.

Dereli, I., V. Telychko, F. Papanikos, K. Raveendran, J. Xu et al., 2024 Seeding the meiotic DNA break machinery and initiating recombination on chromosome axes. Nat Commun 15: 2941.

Dernburg, A. F., K. McDonald, G. Moulder, R. Barstead, M. Dresser et al., 1998 Meiotic recombination in C. elegans initiates by a conserved mechanism and is dispensable for homologous chromosome synapsis. Cell 94: 387–398.

Deshpande, Rajashree A., J.-H. Lee, S. Arora and Tanya T. Paull, 2016 Nbs1 Converts the Human Mre11/Rad50 Nuclease Complex into an Endo/Exonuclease Machine Specific for Protein-DNA Adducts. Molecular Cell 64: 593–606.

Doblmann, J., F. Dusberger, R. Imre, O. Hudecz, F. Stanek et al., 2019 apQuant: Accurate Label-Free Quantification by Quality Filtering. Journal of Proteome Research 18: 535–541.

Dorfer, V., P. Pichler, T. Stranzl, J. Stadlmann, T. Taus et al., 2014 MS Amanda, a Universal Identification Algorithm Optimized for High Accuracy Tandem Mass Spectra. Journal of Proteome Research 13: 3679–3684.

Ferrandiz, N., C. Barroso, O. Telecan, N. Shao, H. M. Kim et al., 2018 Spatiotemporal regulation of Aurora B recruitment ensures release of cohesion during C. elegans oocyte meiosis. Nat Commun 9: 834.

Galanti, L., M. Peritore, R. Gnügge, E. Cannavo, J. Heipke et al., 2024 Dbf4-dependent kinase promotes cell cycle controlled resection of DNA double-strand breaks and repair by homologous recombination. Nature Communications 15: 2890.

Garcia, V., S. E. L. Phelps, S. Gray and M. J. Neale, 2011 Bidirectional resection of DNA double-strand breaks by Mre11 and Exo1. Nature 479: 241–244.

Garcia-Muse, T., U. Galindo-Diaz, M. Garcia-Rubio, J. S. Martin, J. Polanowska et al., 2019 A Meiotic Checkpoint Alters Repair Partner Bias to Permit Inter-sister Repair of Persistent DSBs. Cell Rep 26: 775–787 e775.

Gartner, A., and J. Engebrecht, 2022 DNA repair, recombination, and damage signaling. Genetics 220.

Gerton, J. L., and R. S. Hawley, 2005 Homologous chromosome interactions in meiosis: diversity amidst conservation. Nat Rev Genet 6: 477–487.

Guo, H., E. L. Stamper, A. Sato-Carlton, M. A. Shimazoe, X. Li et al., 2022 Phosphoregulation of DSB-1 mediates control of meiotic double-strand break activity. Elife 11.

Harper, N. C., R. Rillo, S. Jover-Gil, Z. J. Assaf, N. Bhalla et al., 2011 Pairing centers recruit a Polo-like kinase to orchestrate meiotic chromosome dynamics in C. elegans. Dev Cell 21: 934–947.

Hartsuiker, E., K. Mizuno, M. Molnar, J. Kohli, K. Ohta et al., 2009 Ctp1CtIP and Rad32Mre11 Nuclease Activity Are Required for Rec12Spo11 Removal, but Rec12Spo11 Removal Is Dispensable for Other MRN-Dependent Meiotic Functions. Molecular and Cellular Biology 29: 1671–1681.

Haversat, J., A. Woglar, K. Klatt, C. C. Akerib, V. Roberts et al., 2022 Robust designation of meiotic crossover sites by CDK-2 through phosphorylation of the MutSgamma complex. Proc Natl Acad Sci U S A 119: e2117865119.

He, W., H. Rao, S. Tang, N. Bhagwat, D. S. Kulkarni et al., 2020 Regulated Proteolysis of MutSgamma Controls Meiotic Crossing Over. Mol Cell 78: 168–183 e165.

Hefel, A., M. Honda, N. Cronin, K. Harrell, P. Patel et al., 2021 RPA complexes in Caenorhabditis elegans meiosis; unique roles in replication, meiotic recombination and apoptosis. Nucleic Acids Research 49: 2005–2026.

Hicks, T., S. Trivedi, M. Eppert, R. Bowman, H. Tian et al., 2022 Continuous double-strand break induction and their differential processing sustain chiasma formation during Caenorhabditis elegans meiosis. Cell Reports 40: 111403.

Hodgkin, J., H. R. Horvitz and S. Brenner, 1979 NONDISJUNCTION MUTANTS OF THE NEMATODE CAENORHABDITIS ELEGANS. Genetics 91: 67–94.

Hollenstein, D., 2026 MsReport v0.0.34 - Hotfix: Multi-group LIMMA, pp. Zenodo. Hollenstein, D. M., and M. Hartl, 2025 hollenstein/xlsxreport, pp. Zenodo.

Hong, Y., R. Sonneville, B. Wang, V. Scheidt, B. Meier et al., 2018a LEM-3 is a midbody-tethered DNA nuclease that resolves chromatin bridges during late mitosis. Nature Communications 9: 728.

Hong, Y., M. Velkova, N. Silva, M. Jagut, V. Scheidt et al., 2018b The conserved LEM-3/Ankle1 nuclease is involved in the combinatorial regulation of meiotic recombination repair and chromosome segregation in Caenorhabditis elegans. PLOS Genetics 14: e1007453.

Huang, P.-C., S. Hong, H. F. Alnaser, E. P. Mimitou, K. P. Kim et al., 2025 Meiotic DNA break resection and recombination rely on chromatin remodeler Fun30. The EMBO Journal 44: 200–224.

Hunter, N., 2015 Meiotic Recombination: The Essence of Heredity. Cold Spring Harb Perspect Biol 7.

Hurlock, M. E., I. Čavka, L. E. Kursel, J. Haversat, M. Wooten et al., 2020 Identification of novel synaptonemal complex components in C. elegans. Journal of Cell Biology 219: e201910043.

Janisiw, E., M. R. Dello Stritto, V. Jantsch and N. Silva, 2018 BRCA1-BARD1 associate with the synaptonemal complex and pro-crossover factors and influence RAD-51 dynamics during Caenorhabditis elegans meiosis. PLoS Genet 14: e1007653.

Käll, L., J. D. Canterbury, J. Weston, W. S. Noble and M. J. MacCoss, 2007 Semi-supervised learning for peptide identification from shotgun proteomics datasets. Nature Methods 4: 923–925.

Kamp, J. A., R. van Schendel, I. W. Dilweg and M. Tijsterman, 2020 BRCA1-associated structural variations are a consequence of polymerase theta-mediated end-joining. Nat Commun 11: 3615.

Kar, F. M., and A. Hochwagen, 2021 Phospho-Regulation of Meiotic Prophase. Front Cell Dev Biol 9: 667073.

Katis, V. L., J. J. Lipp, R. Imre, A. Bogdanova, E. Okaz et al., 2010 Rec8 phosphorylation by casein kinase 1 and Cdc7-Dbf4 kinase regulates cohesin cleavage by separase during meiosis. Dev Cell 18: 397–409.

Kaur, T., and M. V. Rockman, 2014 Crossover Heterogeneity in the Absence of Hotspots in Caenorhabditis elegans. Genetics 196: 137–148.

Keelagher, R. E., V. E. Cotton, A. S. H. Goldman and R. H. Borts, 2011 Separable roles for Exonuclease I in meiotic DNA double-strand break repair. DNA Repair 10: 126–137.

Kelly, K. O., A. F. Dernburg, G. M. Stanfield and A. M. Villeneuve, 2000 Caenorhabditis elegans msh-5 Is Required for Both Normal and Radiation-Induced Meiotic Crossing Over but Not for Completion of Meiosis. Genetics 156: 617–630.

Kim, S., H. F. Alnaser, S. Keeney and H. Murakami, 2025a Insight into meiotic DNA end resection: Mechanisms and regulation. DNA Repair (Amst) 153: 103886.

Kim, S., S. Yamada, T. Li, C. Canasto-Chibuque, J. H. Kim et al., 2025b Mouse MRE11-RAD50-NBS1 is needed to start and extend meiotic DNA end resection. Nat Commun 16: 3613.

Kim, Y., N. Kostow and A. F. Dernburg, 2015 The Chromosome Axis Mediates Feedback Control of CHK-2 to Ensure Crossover Formation in C. elegans. Dev Cell 35: 247–261.

Kimble, M. T., M. J. Johnson, M. R. Nester and L. S. Symington, 2023 Long-range DNA end resection supports homologous recombination by checkpoint activation rather than extensive homology generation. eLife 12: e84322.

Kirkpatrick, D. T., J. R. Ferguson, T. D. Petes and L. S. Symington, 2000 Decreased Meiotic Intergenic Recombination and Increased Meiosis I Nondisjunction in exo1 Mutants of Saccharomyces cerevisiae. Genetics 156: 1549–1557.

Koury, E., K. Harrell and S. Smolikove, 2018 Differential RPA-1 and RAD-51 recruitment in vivo throughout the C. elegans germline, as revealed by laser microirradiation. Nucleic Acids Res 46: 748–764.

Kruisselbrink, E., V. Guryev, K. Brouwer, D. B. Pontier, E. Cuppen et al., 2008 Mutagenic Capacity of Endogenous G4 DNA Underlies Genome Instability in FANCJ-Defective C. elegans. Current Biology 18: 900–905.

Kung, J. E., and N. Jura, 2019 Prospects for pharmacological targeting of pseudokinases. Nature Reviews Drug Discovery 18: 501–526.

Kwon, A., S. Scott, R. Taujale, W. Yeung, K. J. Kochut et al., 2019 Tracing the origin and evolution of pseudokinases across the tree of life. Science Signaling 12: eaav3810.

Labella, S., A. Woglar, V. Jantsch and M. Zetka, 2011 Polo kinases establish links between meiotic chromosomes and cytoskeletal forces essential for homolog pairing. Dev Cell 21: 948–958.

Lascarez-Lagunas, L. I., S. Nadarajan, M. Martinez-Garcia, J. N. Quinn, E. Todisco et al., 2022 ATM/ATR kinases link the synaptonemal complex and DNA double-strand break repair pathway choice. Curr Biol 32: 4719–4726 e4714.

León-Ortiz, A. M., S. Panier, G. Sarek, J.-B. Vannier, H. Patel et al., 2018 A Distinct Class of Genome Rearrangements Driven by Heterologous Recombination. Molecular Cell 69: 292–305.e296.

Li, Q., T. T. Saito, M. Martinez-Garcia, A. J. Deshong, S. Nadarajan et al., 2018 The tumor suppressor BRCA1-BARD1 complex localizes to the synaptonemal complex and regulates recombination under meiotic dysfunction in Caenorhabditis elegans. PLOS Genetics 14: e1007701.

Lo, H. C., L. Wan, A. Rosebrock, B. Futcher and N. M. Hollingsworth, 2008 Cdc7-Dbf4 regulates NDT80 transcription as well as reductional segregation during budding yeast meiosis. Mol Biol Cell 19: 4956–4967.

Machovina, T. S., R. Mainpal, A. Daryabeigi, O. McGovern, D. Paouneskou et al., 2016 A Surveillance System Ensures Crossover Formation in C. elegans. Curr Biol 26: 2873–2884.

Martinez-Perez, E., M. Schvarzstein, C. Barroso, J. Lightfoot, A. F. Dernburg et al., 2008 Crossovers trigger a remodeling of meiotic chromosome axis composition that is linked to two-step loss of sister chromatid cohesion. Genes & Development 22: 2886–2901.

Martinez-Perez, E., and A. M. Villeneuve, 2005 HTP-1-dependent constraints coordinate homolog pairing and synapsis and promote chiasma formation during C. elegans meiosis. Genes Dev 19: 2727–2743.

Martini, E., R. L. Diaz, N. Hunter and S. Keeney, 2006 Crossover Homeostasis in Yeast Meiosis. Cell 126: 285–295.

Matos, J., J. J. Lipp, A. Bogdanova, S. Guillot, E. Okaz et al., 2008 Dbf4-dependent CDC7 kinase links DNA replication to the segregation of homologous chromosomes in meiosis I. Cell 135: 662–678.

Milman, N., E. Higuchi and G. R. Smith, 2009 Meiotic DNA Double-Strand Break Repair Requires Two Nucleases, MRN and Ctp1, To Produce a Single Size Class of Rec12 (Spo11)-Oligonucleotide Complexes. Molecular and Cellular Biology 29: 5998–6005.

Mimitou, E. P., and L. S. Symington, 2008 Sae2, Exo1 and Sgs1 collaborate in DNA double-strand break processing. Nature 455: 770–774.

Mimitou, E. P., S. Yamada and S. Keeney, 2017 A global view of meiotic double-strand break end resection. Science 355: 40–45.

Moreau, S., J. R. Ferguson and L. S. Symington, 1999 The Nuclease Activity of Mre11 Is Required for Meiosis but Not for Mating Type Switching, End Joining, or Telomere Maintenance. Molecular and Cellular Biology 19: 556–566.

Murakami, H., and S. Keeney, 2014 DDK links replication and recombination in meiosis. Cell Cycle 13: 3621–3622.

Nabeshima, K., A. M. Villeneuve and M. P. Colaiácovo, 2005 Crossing over is coupled to late meiotic prophase bivalent differentiation through asymmetric disassembly of the SC. Journal of Cell Biology 168: 683–689.

Nadarajan, S., T. J. Lambert, E. Altendorfer, J. Gao, M. D. Blower et al., 2017 Polo-like kinase-dependent phosphorylation of the synaptonemal complex protein SYP-4 regulates double-strand break formation through a negative feedback loop. Elife 6.

Nadarajan, S., F. Mohideen, Y. B. Tzur, N. Ferrandiz, O. Crawley et al., 2016 The MAP kinase pathway coordinates crossover designation with disassembly of synaptonemal complex proteins during meiosis. Elife 5: e12039.

Natsume, T., C. A. Muller, Y. Katou, R. Retkute, M. Gierlinski et al., 2013 Kinetochores coordinate pericentromeric cohesion and early DNA replication by Cdc7-Dbf4 kinase recruitment. Mol Cell 50: 661–674.

Neale, M. J., J. Pan and S. Keeney, 2005 Endonucleolytic processing of covalent protein-linked DNA double-strand breaks. Nature 436: 1053–1057.

Paix, A., A. Folkmann, D. Rasoloson and G. Seydoux, 2015 High Efficiency, Homology-Directed Genome Editing in Caenorhabditis elegans Using CRISPR-Cas9 Ribonucleoprotein Complexes. Genetics 201: 47–54.

Pattabiraman, D., B. Roelens, A. Woglar and A. M. Villeneuve, 2017 Meiotic recombination modulates the structure and dynamics of the synaptonemal complex during C. elegans meiosis. PLoS Genet 13: e1006670.

Perez-Riverol, Y., C. Bandla, Deepti J. Kundu, S. Kamatchinathan, J. Bai et al., 2025 The PRIDE database at 20 years: 2025 update. Nucleic Acids Research 53: D543–D553.

Pothof, J., G. van Haaften, K. Thijssen, R. S. Kamath, A. G. Fraser et al., 2003 Identification of genes that protect the C. elegans genome against mutations by genome-wide RNAi. Genes & Development 17: 443–448.

Ritchie, M. E., B. Phipson, D. Wu, Y. Hu, C. W. Law et al., 2015 limma powers differential expression analyses for RNA-sequencing and microarray studies. Nucleic Acids Research 43: e47–e47.

Rockman, M. V., and L. Kruglyak, 2009 Recombinational Landscape and Population Genomics of Caenorhabditis elegans. PLOS Genetics 5: e1000419.

Rosu, S., K. A. Zawadzki, E. L. Stamper, D. E. Libuda, A. L. Reese et al., 2013 The C. elegans DSB-2 protein reveals a regulatory network that controls competence for meiotic DSB formation and promotes crossover assurance. PLoS Genet 9: e1003674.

Salunkhe, S., J. M. Daley, H. Kaur, N. Tomimatsu, C. Xue et al., 2024 Promotion of DNA end resection by BRCA1–BARD1 in homologous recombination. Nature 634: 482–491.

Sarkar, G. C., U. Rautela, A. Goyala, S. Datta, N. Anand et al., 2023 DNA damage signals from somatic uterine tissue arrest oogenesis through activated DAF-16. Development 150: dev201472.

Sasanuma, H., K. Hirota, T. Fukuda, N. Kakusho, K. Kugou et al., 2008 Cdc7-dependent phosphorylation of Mer2 facilitates initiation of yeast meiotic recombination. Genes Dev 22: 398–410.

Sato-Carlton, A., C. Nakamura-Tabuchi, S. K. Chartrand, T. Uchino and P. M. Carlton, 2017 Phosphorylation of the synaptonemal complex protein SYP-1 promotes meiotic chromosome segregation. Journal of Cell Biology 217: 555–570.

Schneider, C. A., W. S. Rasband and K. W. Eliceiri, 2012 NIH Image to ImageJ: 25 years of image analysis. Nature Methods 9: 671–675.

Schwanhäusser, B., D. Busse, N. Li, G. Dittmar, J. Schuchhardt et al., 2011 Global quantification of mammalian gene expression control. Nature 473: 337–342.

Silva, N., N. Ferrandiz, C. Barroso, S. Tognetti, J. Lightfoot et al., 2014 The fidelity of synaptonemal complex assembly is regulated by a signaling mechanism that controls early meiotic progression. Dev Cell 31: 503–511.

Smyth, G. K., 2004 Linear Models and Empirical Bayes Methods for Assessing Differential Expression in Microarray Experiments. 3.

Sofroniew, N., T. Lambert, G. Bokota, J. Nunez-Iglesias, P. Sobolewski et al., 2026 napari: a multi-dimensional image viewer for Python (v0.7.0), pp. Zenodo.

Stamper, E. L., S. E. Rodenbusch, S. Rosu, J. Ahringer, A. M. Villeneuve et al., 2013 Identification of DSB-1, a protein required for initiation of meiotic recombination in Caenorhabditis elegans, illuminates a crossover assurance checkpoint. PLoS Genet 9: e1003679.

Stringer, C., T. Wang, M. Michaelos and M. Pachitariu, 2021 Cellpose: a generalist algorithm for cellular segmentation. Nature Methods 18: 100–106.

Sugiyama, T., and S. C. Kowalczykowski, 2002 Rad52 Protein Associates with Replication Protein A (RPA)-Single-stranded DNA to Accelerate Rad51-mediated Displacement of RPA and Presynaptic Complex Formation*. Journal of Biological Chemistry 277: 31663–31672.

Taus, T., T. Köcher, P. Pichler, C. Paschke, A. Schmidt et al., 2011 Universal and Confident Phosphorylation Site Localization Using phosphoRS. Journal of Proteome Research 10: 5354–5362.

The C. elegans Deletion Mutant Consortium, 2012 Large-Scale Screening for Targeted Knockouts in the Caenorhabditis elegans Genome. G3 Genes|Genomes|Genetics 2: 1415–1425.

The UniProt, C., 2023 UniProt: the Universal Protein Knowledgebase in 2023. Nucleic Acids Research 51: D523–D531.

Toraason, E., M. Glover, A. Horacek and D. E. Libuda, 2021 Detection of homolog-independent meiotic DNA repair events in C. elegans with the intersister/intrachromatid repair assay. STAR Protocols 2: 100801.

Toraason, E., A. Salagean, D. E. Almanzar, J. E. Brown, C. M. Richter et al., 2024 BRCA1/BRC-1 and SMC-5/6 regulate DNA repair pathway engagement during Caenorhabditis elegans meiosis. eLife 13: e80687.

Valentin, G., E. Schwob and F. Della Seta, 2006 Dual role of the Cdc7-regulatory protein Dbf4 during yeast meiosis. J Biol Chem 281: 2828–2834.

van Kempen, M., S. S. Kim, C. Tumescheit, M. Mirdita, J. Lee et al., 2024 Fast and accurate protein structure search with Foldseek. Nature Biotechnology 42: 243–246.

Varadi, M., S. Anyango, M. Deshpande, S. Nair, C. Natassia et al., 2022 AlphaFold Protein Structure Database: massively expanding the structural coverage of protein-sequence space with high-accuracy models. Nucleic Acids Research 50: D439–D444.

Velez-Aguilera, G., B. Ossareh-Nazari and L. Pintard, 2024 Dissecting the Multiple Functions of the Polo-Like Kinase 1 in the C. elegans Zygote, pp. 63–88 in Cell Cycle Control: Methods and Protocols, edited by A. Castro and B. Lacroix. Springer US, New York, NY.

Velkova, M., N. Silva, M. R. Dello Stritto, A. Schleiffer, P. Barraud et al., 2021 Caenorhabditis elegans RMI2 functional homolog-2 (RMIF-2) and RMI1 (RMH-1) have both overlapping and distinct meiotic functions within the BTR complex. PLoS Genet 17: e1009663.

Wan, L., H. Niu, B. Futcher, C. Zhang, K. M. Shokat et al., 2008 Cdc28-Clb5 (CDK-S) and Cdc7-Dbf4 (DDK) collaborate to initiate meiotic recombination in yeast. Genes Dev 22: 386–397.

Woglar, A., A. Daryabeigi, A. Adamo, C. Habacher, T. Machacek et al., 2013 Matefin/SUN-1 phosphorylation is part of a surveillance mechanism to coordinate chromosome synapsis and recombination with meiotic progression and chromosome movement. PLoS Genet 9: e1003335.

Wold, M. S., 1997 Replication protein A: a heterotrimeric, single-stranded DNA-binding protein required for eukaryotic DNA metabolism. Annual Review of Biochemistry 66: 61–92.

Xia, Y., R. Sonneville, M. Jenkyn-Bedford, L. Ji, C. Alabert et al., 2023 DNSN-1 recruits GINS for CMG helicase assembly during DNA replication initiation in Caenorhabditis elegans. Science 381: eadi4932.

Yin, Y., S. Donlevy and S. Smolikove, 2016 Coordination of Recombination with Meiotic Progression in the Caenorhabditis elegans Germline by KIN-18, a TAO Kinase That Regulates the Timing of MPK-1 Signaling. Genetics 202: 45–59.

Yin, Y., and S. Smolikove, 2013 Impaired resection of meiotic double-strand breaks channels repair to nonhomologous end joining in Caenorhabditis elegans. Mol Cell Biol 33: 2732–2747.

Yokoo, R., Karl A. Zawadzki, K. Nabeshima, M. Drake, S. Arur et al., 2012 COSA-1 Reveals Robust Homeostasis and Separable Licensing and Reinforcement Steps Governing Meiotic Crossovers. Cell 149: 75–87.

Yu, Z., H. J. Kim and A. F. Dernburg, 2023 ATM signaling modulates cohesin behavior in meiotic prophase and proliferating cells. Nat Struct Mol Biol 30: 436–450.

Zakharyevich, K., Y. Ma, S. Tang, P. Y.-H. Hwang, S. Boiteux et al., 2010 Temporally and Biochemically Distinct Activities of Exo1 during Meiosis: Double-Strand Break Resection and Resolution of Double Holliday Junctions. Molecular Cell 40: 1001–1015.

Zhang, L., W. T. Stauffer, J. S. Wang, F. Wu, Z. Yu et al., 2023 Recruitment of Polo-like kinase couples synapsis to meiotic progression via inactivation of CHK-2. Elife 12.

Zheng, G., M. Kanchwala, C. Xing and H. Yu, 2018 MCM2-7-dependent cohesin loading during S phase promotes sister-chromatid cohesion. Elife 7.

Zickler, D., and N. Kleckner, 2015 Recombination, Pairing, and Synapsis of Homologs during Meiosis. Cold Spring Harb Perspect Biol 7.

