## Supplementary material for "Dbf4-dependent kinase promotes meiotic DNA end resection through cyclin-dependent kinase 12 and DNA-2 in *Caenorhabditis elegans*": Suplement_text

**CDK-12 in the context of *cdc-7* mutants**

Given that CDK-12 is best known for phosphorylating the C-terminal domain (CTD) of RNA polymerase II (Bowman *et al.* 2013), we asked whether loss of *cdc-7* affects CDK-12 localization or its canonical activity. To this end, we first engineered a tagged version of CDK-12 using CRISPR/Cas9 to confirm that CDK-12 localization is not affected by *cdc-7* deletion and, second, probed the activity of CDK-12 in the germline using an antibody that specifically detects the phosphorylated form of the CTD (Figure S5).

Using a tagged CDK-12 allele (*syb10816*, hereafter *cdk-12::FLAG*), CDK-12::FLAG displayed a punctate and nucleoplasmic localization in wild-type germlines (Figure S5A). In *cdc-7∆* mutants, the signal appeared markedly brighter (Figure S5A). To quantify the difference, we measured the mean FLAG intensity in segmented nuclei (see Materials and Methods) across the germline (divided into seven zones of equal length; Figure S5B), and calculated the ratio of signal intensity between *cdc-7∆* mutants and the reference strain for each zone. Across the germline, CDK-12::FLAG levels were increased by approximately 4- to 8-fold in *cdc-7∆* mutants relative to the wild type (Figure S5C). These results indicate that loss of *cdc-7* leads to elevated CDK-12 levels.

We next assessed CDK-12 kinase activity by monitoring phosphorylation of serine 2 within the CTD of RNA polymerase II. Phospho-CTD staining appeared similar between wild type and *cdc-7∆* mutants (Figure S5D), and quantification using the same approach revealed differences of less than 2-fold across germline zones (Figure S5E,F). These results indicate that despite its increased levels, the canonical CTD kinase activity of CDK-12 is largely unchanged in *cdc-7∆* mutants.

Together, these results indicate that *cdc-7* loss leads to increased CDK-12 expression without altering the canonical CTD kinase activity, thus supporting a model in which CDK-12 performs an additional, DDK-dependent function during meiosis.
